# VesperFlow: An automated pipeline for decoding multisensor data from biologging tags

**DOI:** 10.64898/2026.10.12.758513

**Authors:** David Alexandre Silva, Mehran Eqbal, Lily Broos, Richard Vogg, Kaja Wierucka

## Abstract

Integrating multi-sensor data from biologging tags is key to understanding animal behavior, particularly in group-living species, where communication, movement, and spatial dynamics are closely linked. However, analyzing such data at scale is often hindered by proprietary binary formats or manual decoding workflows in existing software. We present VesperFlow, an open-source Python pipeline that automates the decoding of inertial measurement unit (IMU), audio, and GPS data from Vesper Wildlife tags. VesperFlow batch-processes large datasets from multiple individuals, synchronizes timestamps across sensors with sub-second precision, and removes hardware-induced artifacts such as spurious gyroscope spikes and audio pops. Outputs are exported as CSV and WAV files for direct use in common downstream analysis workflows. We validate the pipeline on field data from 10 ring-tailed lemurs (*Lemur catta*), showing that our pipeline reliably processes movement, vocal, and spatial data. By lowering technical barriers and automating processes, VesperFlow supports more scalable, multi-modal research on animal behavior. VesperFlow is freely available at https://github.com/dasilva98/vesperflow.

## Introduction

Animal behaviour is shaped by multiple, interacting sources of information. Animals may respond to signals and cues across several sensory modalities (e.g., acoustic or visual information) (1), as well as to the presence, movement, or activity of nearby individuals (2–4). These responses may depend not only on which individuals are present (including conspecifics and members of other species) (1), but also on various social factors such as social associations (5, 6), dominance status (7), or relatedness. These sources of information are dynamic and interdependent, and examining any one of them in isolation may provide only a partial account of the processes underlying individual and collective behaviour. A more complete understanding therefore requires the simultaneous measurement of multiple communication channels, behavioural states, social and spatial relationships. The interplay between these factors is highly complex and dynamic. Behavioral research thus increasingly relies on multi-sensor tags to capture fine-scale data across multiple dimensions.

Relatively few studies have integrated acoustic, behavioural, and spatial data from multiple individuals within a single empirical framework. Most examine one dimension in isolation (8–11) or look at the interaction of two dimensions, for example by linking communication or collective movement with behaviour (12–14). This gap is largely due to technological limitations in monitoring many individuals at once and integrating their vocal, spatial and behavioural activity simultaneously (15). Many available commercial biologging devices consist of only one or two integrated sensors, offer limited configuration options, or are designed for a restricted range of species. In-house systems can provide greater flexibility, but may lack detailed documentation, tutorials, or community support for data acquisition and processing. Consequently, adopting such tools often requires high levels of technical expertise on the part of the researcher, while the needed beta testing and troubleshooting can be time-consuming.

The Vesper Wildlife tag system (16) supports the simultaneous collection of acoustic, GPS, and inertial measurement unit (IMU) data. The system can also integrate a proximity sensor. It therefore provides an excellent platform for investigating individual and collective behaviour across a range of study designs. Data can be extracted and decoded using the accompanying VesperApp, which provides an accessible interface for processing tag recordings. However, workflows involving large numbers of files may benefit from additional automation and programmatic access. In particular, file-by-file processing, non-standardized file names, and outputs distributed across multiple locations can make large datasets more difficult to organize and process efficiently. Additional processing is also required to account for known features of the raw data, including gyroscope spikes associated with file-split delays, audio initialization artifacts, or non-compliant WAV headers.

Here, we introduce VesperFlow, a data-processing pipeline designed to streamline the decoding and organization of multisensor data collected from multiple animals using Vesper Wildlife tags (16). VesperFlow provides a local, serverless workflow that automatically identifies recording sessions and applies sensor-specific decoding procedures. It incorporates existing GPS decoders and provides open-source implementations for parsing IMU and audio data, allowing raw tag recordings to be integrated into reproducible analysis workflows with minimal manual intervention. Specifically, this paper provides A) a guide to VesperFlow, a new tool for decoding and flexibly assorting large quantities of multi-sensor data; B) a case study demonstrating the application of this system in a real-world research context. Part B also includes a practical deployment toolkit comprising a user guide for the tag, and a ready-to-print 3D collar model, and code for configuring the proximity sensor.

### VesperFlow

The processing of high-resolution bio-logging data requires robust, automated software capable of handling thousands of multi-modal recording files. VesperFlow employs a direct, headless Command Line Interface (CLI) architecture designed specifically for machine learning and database ingestion workflows. The pipeline streams and unpacks binary payloads in-memory, eliminating the I/O bottleneck of intermediate disk writes. Leveraging vectorized array operations, the tool routes data directly from the raw tag output into clean, analysis-ready formats. The system’s architecture is divided into three successive phases: (1) Data ingestion, (2) Binary decoding & parsing, and (3) Export (Figure 1).

**Fig. 1.**
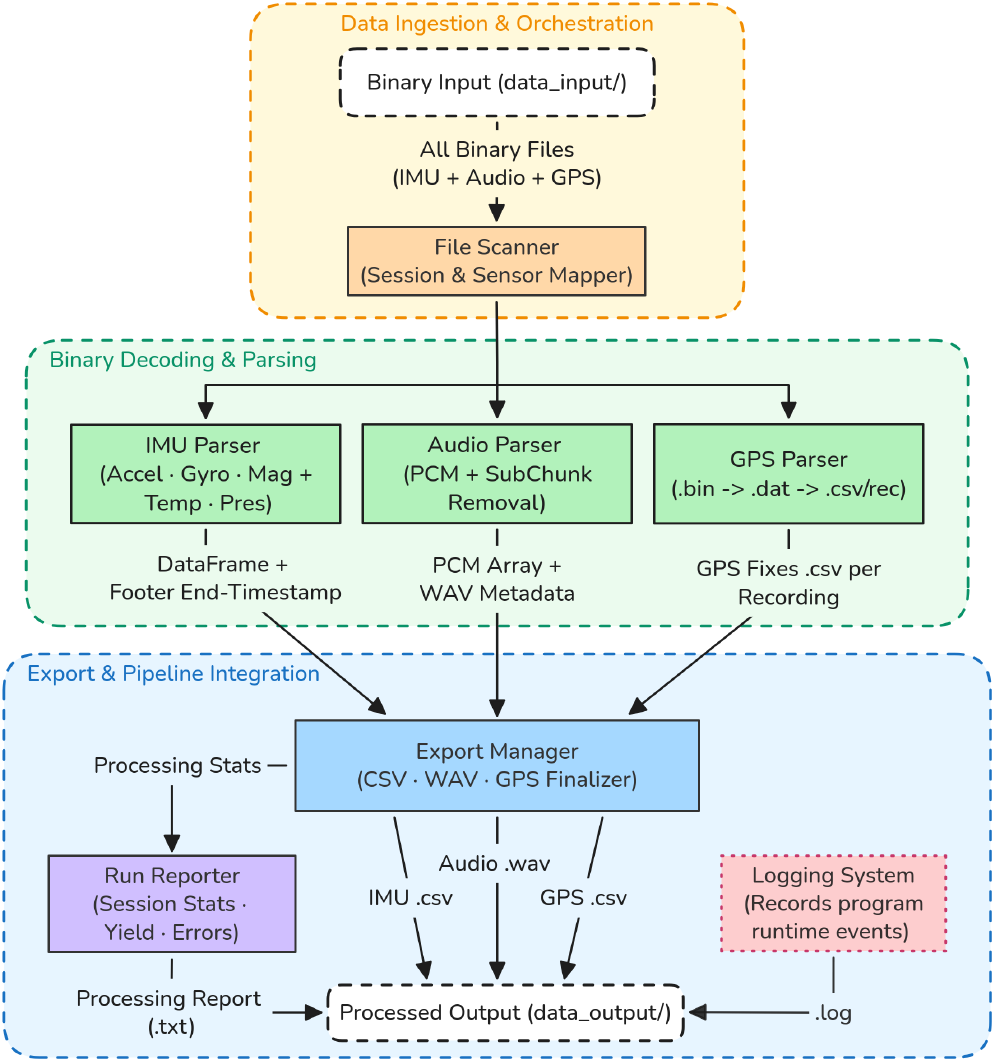
VesperFlow system architecture diagram. The pipeline sequentially processes data through three stages: batch ingestion & orchestration, sensor-specific binary decoding & stream parsing, and standardized export for database integration. Abbreviations: Accel=Accelerometer, Gyro=Gyroscope, Mag=Magnetometer, Temp=Temperature, Pres=Pressure and PCM=Pulse-Code Modulation.

#### (1) Data ingestion & orchestration

##### Headless CLI and batch processing

VesperFlow is implemented as a scriptable, batch-processing CLI designed for automated execution within large-scale telemetry pipelines. While traditional graphical user interface (GUI) extraction tools require manual navigation across individual deployment directories, this CLI operates unattended, allowing multi-deployment archives to be ingested directly into down-stream analytical workflows.

##### Session discovery and chunk flattening

The pipeline processes all binary files from a given tag deployment as a continuous, unified session. To accommodate FAT32 directory constraints and flash allocation limits, Vesper firmware partitions recordings across auto-incrementing subdirectories (e.g., `0`, `1`, `2`), capping each folder at 256 files. VesperFlow’s file scanner recursively crawls the deployment root, resolving directory fragmentation to assemble all chunks into a single chronological session.

##### Classification and chronological sorting

Files are classified using the manufacturer’s standardized single-letter file-name suffixes: `M.BIN` for IMU, `U.BIN` for audio, and `G.BIN` for GPS. Multi-file streams are sorted numerically by their leading sequence index (e.g., `0M.BIN`, `1M.BIN`, …) to guarantee strict temporal continuity before concatenation. Additionally, the file scanner filters out hidden operating system artefacts (such as `.DS_Store` or index files) to prevent non-payload data from triggering parser errors.

##### Orchestration and fault isolation

Once a session is mapped, payloads are routed independently through decoupled IMU, audio, and GPS parsing stages. This modular architecture enforces strict fault isolation: an I/O error in an audio block or an incomplete GPS snapshot buffer does not abort processing for the other modalities, ensuring that localized corruptions do not invalidate an entire deployment. During processing, the orchestrator logs an execution status for every file: SUCCESS, EMPTY (indicating a valid header without payload, typical of mid-recording power loss), or FAIL (indicating structural or parity corruption).

#### (2) Binary decoding & parsing

##### Decoding principles and metadata

Standardized media containers, such as the Resource Interchange File Format (RIFF), rely on self-describing Four-Character Codes (FourCC) like `fmt` and `data` (17, 18). Raw Vesper binary streams, by contrast, lack built-in syntactic descriptors (19). Consequently, to extract telemetry data, a compatible parser must rely on predefined schemas aligned to little-endian byte boundaries. To anchor the parser and validate structural integrity within the raw bitstream, the VesperFlow decoding engine employs magic numbers (fixed hexadecimal signatures) (20). Specifically, it scans for `0xC0DAAFDE` to mark the start of a file header, the sync pattern `0 × 5AA55AA5` for the block header, the inverse sync pattern `0xA55AA55A` for the block trailer, and `0xEFEFCDAB` to locate periodic sync markers across audio files.

Additionally, the tag’s on-board real-time clock (RTC) uses Binary-Coded Decimal (BCD) encoding (21). In this format, each decimal digit occupies a single 4-bit nibble; for example, the decimal value 25 is stored in memory as `0 × 25`. Because standard binary integer deserialization interprets this as decimal 37, VesperFlow decodes each byte into its corresponding decimal value using Equation 1:

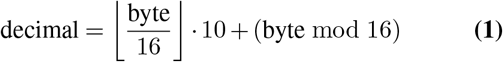

##### The 128-byte file header and 16-byte block header

VesperFlow mirrors the hardware’s on-device layout by parsing the stream into a discrete 128-byte file header followed by a 16-byte block header (Figure 2) (with the exception of GPS files, which contain only an initial 16-byte header). Every IMU and audio file begins with the 128-byte file header (Figure 2). This structure encodes the 32-bit device unique identifier (UID), a 16-byte ASCII sensor type (e.g., `IMU10` or `SPH0641`), the firmware version, the sampling rate, active bitmasks, and hardware configuration registers. Immediately following this is the initial 16-byte block header, marked by the `0 × 5AA55AA5` magic number, which acts as a sync pattern (20). This block header contains sub-second resolution metrics alongside hardware date and time fields, generated natively by the microcontroller’s RTC in BCD format.

**Fig. 2.**
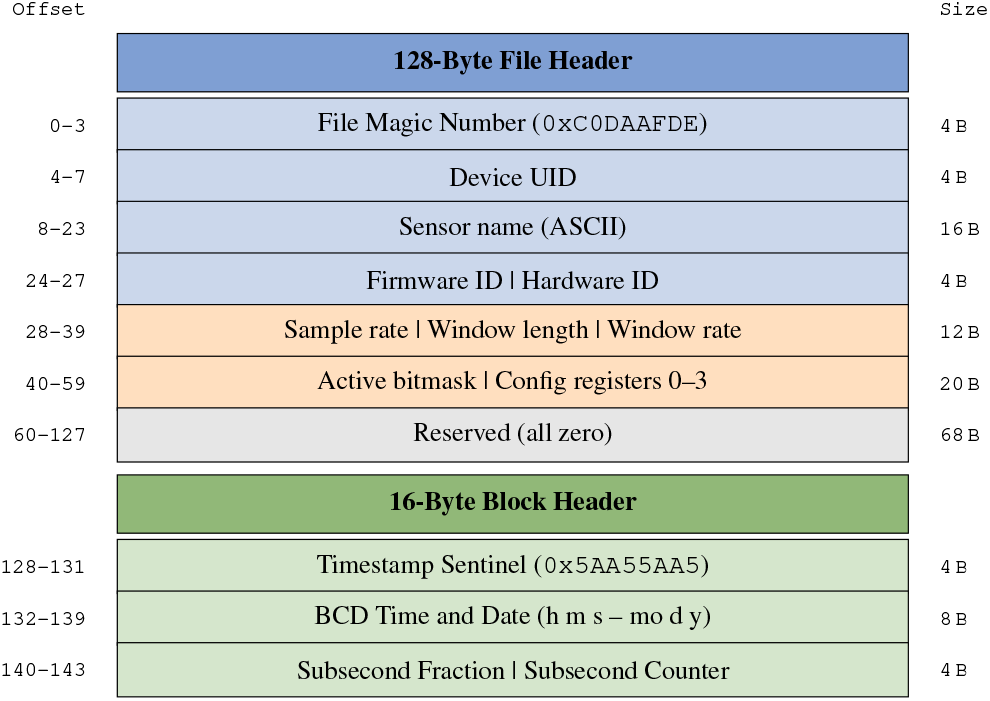
Schematic layout of the Vesper binary structure. The 128-byte global File Header establishes identity and sensor configuration, while the 16-byte Block Header provides precise temporal synchronization. Blue: identity; orange: configuration; gray: reserved; green: timestamp region.

##### Sub-second start time

Standard BCD calendar registers provide a maximum resolution of one second, which is insufficient for time-series data sampled at frequencies between 50 Hz and 48 kHz. To achieve precise temporal alignment, the pipeline extracts the final four bytes of the block header, which encode the RTC’s internal subsecond timer state (21). These bytes represent two 16-bit integers: the *subsecond fraction* (*S*_frac_, representing the hardware timer prescaler frequency minus one, e.g., 1023 for a 1024 Hz clock source) and the *sub-second counter* (*S*_cnt_, the down-counter tick value when the file is initialized). The millisecond offset from the preceding whole-second boundary is calculated as:

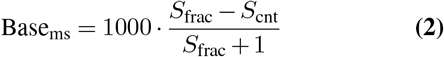

##### Sensor-specific stream parsing

Once header metadata is validated, the payload is routed to one of three dedicated VesperFlow sensor parsers. To handle the large number of recording sessions without the performance overhead of scalar Python loops, these parsers execute vectorized operations over contiguous NumPy memory buffers.

##### IMU stream and true-time synchronization

VesperFlow parses IMU payload structured as contiguous sequences of 42-byte binary packets (or 46 bytes when optional pressure and temperature channels are enabled). Each 42-byte packet contains nine IEEE 754 little-endian single-precision floats (`Float32`) comprising tri-axial angular velocity (degrees per second, °*/*s), tri-axial acceleration (milli-*g*, m*g*), and triaxial magnetic flux density (milligauss, mG), followed by a hardware sample counter. Per-sample timestamps are generated across the array using a vectorized index mapping:

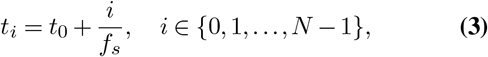

where *t*_0_ represents the hardware start time anchored to the RTC sub-second registers (21), and *f*_*s*_ denotes the nominal sensor sampling rate.

The physical tag firmware exhibits an undocumented I/O delay bug during boundary transitions. When the device closes an active file and allocates a subsequent FAT32 cluster chain, write latency halts CPU execution. Consequently, the firmware duplicates the timestamp row and freezes the gyroscope register value, producing a static artifact value (*™* 8.75 °*/*s). While localized, these duplicate rows can distort downstream feature extraction and cause state misclassifications. VesperFlow resolves this by detecting boundary duplicates and replacing them with `NaN` using vectorized masks directly at the array level.

##### Audio stream and strict RIFF compliance

Audio data is stored as 16-bit signed pulse-code modulation (PCM) sampled at *f*_*s*_ = 48 kHz and written to flash in *≈* 64 KB pages. At each page boundary, the firmware injects a 16-byte sync marker comprising a 4-byte magic number (`0xEFEFCDAB`) (20), an 8-byte BCD timestamp, and a 4-byte subsecond tick register. If ingested directly into the audio buffer, these metadata bytes are misinterpreted as full-scale acoustic samples, producing periodic clipping artifacts every *≈* 0.68 s (65,536 bytes*/*(2 bytes/sample *·* 48,000 Hz)). To resolve this, VesperFlow locates each sync marker magic number and strips the 16 metadata bytes, preserving two-byte word alignment without introducing phase drift over long recordings. Additionally, the parser discards an initial *≈* 17 ms (*≈* 1,000 samples) negative full-scale DC spike (`0 × 8000`), or *™* 32,768) caused by microphone hardware initialization. Finally, the continuous PCM array is encapsulated into a compliant RIFF container by synthesizing a standard 44-byte WAVE header containing valid `RIFF`, `WAVE`, `fmt`, and `data` FourCC chunks (17, 18, 22).

##### GPS snapshots and data alignment

To minimize power consumption on the tags, the hardware bypasses on-board satellite decoding and position calculation. Instead, it captures periodic raw Intermediate Frequency (IF) radio snapshots for post-processed software-defined radio (SDR) solving (23). VesperFlow extracts these captures by first stripping a 1,008-byte null-padded buffer injected into the first session file (`0G.BIN`) while the GNSS front-end phase-locked loop (PLL) stabilizes. To format the raw captures for downstream tracking algorithms, the parser performs a vectorized 16-bit halfword swap across each 32-bit word:

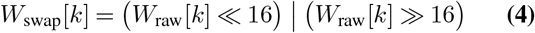

where *W*_raw_[*k*] *∈* U_32_ represents the *k*-th unsigned 32-bit sample word and | denotes the bitwise OR operator. This permutation reorders the lower and upper 16-bit halfwords into the memory layout expected by the GNSS baseband processor. Failing to perform this transposition inverts the order of interleaved In-phase and Quadrature (*I/Q*) radio samples, preventing downstream SDR correlators from synchronizing with satellite pseudo-random noise (PRN) codes.

#### (3) Export & pipeline integration

##### Output structure

Tag firmware partitions recording across sequential files (`0M.BIN`, `1M.BIN`, etc.), fragmenting multi-hour deployments and complicating batch ingestion. The Export Manager consolidates these fragments into a structured, sensor-indexed directory hierarchy (`imu/`, `aud/`, `gps/`). Each output file is named deterministically using the session’s temporal boundaries and hardware identifier:

~~~
YYYYMMDD_HHMMSS-YYYYMMDD_HHMMSS_DEVICEID.ext
~~~

By deriving filenames directly from payload metadata rather than arbitrary sequence counters, completed sessions can ingested into relational databases or automated machine learning pipelines without intermediate renaming.

Depending on the hardware configuration, exported streams appear as follows:

- **IMU (**.**csv**): High-resolution timestamps (`YYYY-MM-DD HH:MM:SS.mmm`) alongside legacy split temporal fields (minute, second, millisecond), calibrated tri-axial acceleration (m*g*), angular velocity (°/s), magnetic flux density (mG), and optional environmental channels (temperature in °C and barometric pressure in hPa).
- **Audio (**.**wav**): Single-channel, 16-bit signed PCM audio sampled at 48 kHz, structured as a RIFF-compliant WAVE container with hardware initialization spikes and boundary sync markers excised.
- **GPS (**.**csv**): Resolved positional telemetry compiled from the SDR post-processing of GNSS snapshots. These records include estimated latitude, longitude, altitude, UTC timestamp, and fix quality metrics.

Each IMU and audio export is paired with a `.txt` metadata sidecar file containing the raw hardware configuration registers extracted from the 128-byte file header, preserving parameter traceability and ensuring data provenance.

##### External decoder integration

Once the `.DAT` snapshots are written, a small wrapper script sends them to `GeoTag.exe`, the manufacturer’s command-line tool, and collects the coordinate file it produces. The external GeoTag.exe is restricted to Windows-only. The executable is unavailable when running the tool on a Linux server or a high-performance computing cluster. The tool currently handles this situation smoothly without halting or crashing. A missing executable message shows up as a critical error in the run report, and everything else (IMU, audio, and the GPS snapshots themselves) is decoded and exported as usual. However, while the current stable release relies on this external Windows tool for the GPS parser module, we intend to swap this dependency with a native cross-platform GPS parser which is currently under development and will be available soon.

##### Quality control and run reporting

To ensure a verifiable data trail for research purposes, the pipeline employs a `RunReporter` module to centrally manage execution metrics. Rather than relying on unstructured console output, the reporter continuously tracks the processing status of every file and compiles a comprehensive plain text summary report (`.txt`) that is saved alongside the exported data (Figure 3). The module collects and aggregates this data across four specific categories:

**Fig. 3.**
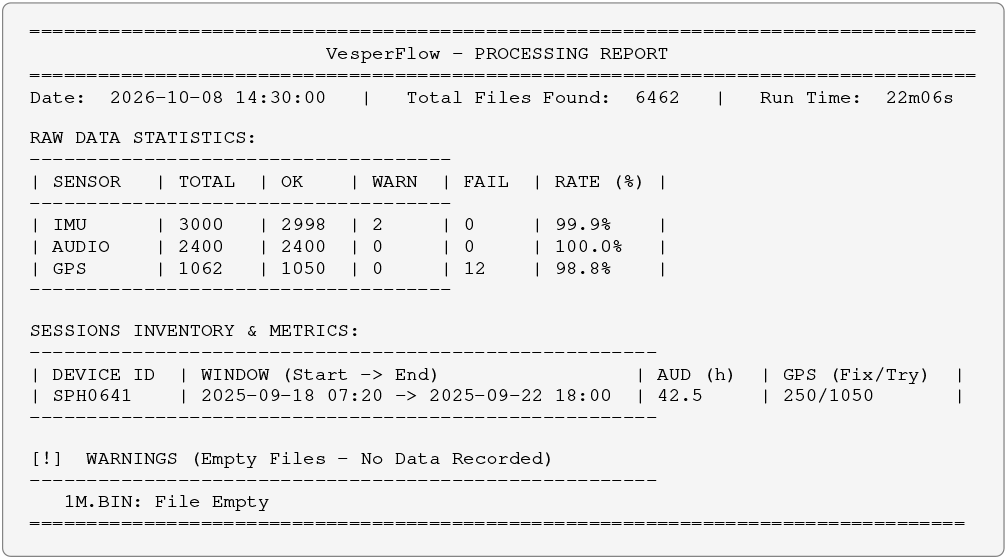
Example excerpt of the generated plaintext processing report (`.txt`). The report consolidates file health, session boundaries, and true scientific yield, ensuring automated pipeline execution is fully documented for downstream analysis.

1. **Raw Data Statistics:** The pipeline tracks all files across three statuses: SUCCESS, WARN, and FAIL. Crucially, files that contain a valid 128-byte header but no data payload, are logged as WARN (*Empty*) rather than FAIL. This prevents natural hardware shutdowns from skewing the pipeline’s failure rate.
2. **Sessions Inventory:** For each tag deployment, the reporter logs the device UID, the absolute temporal window (earliest start to latest end), the actual audio duration in hours, and the GPS fix-to-attempt ratio.
3. **True Data Yield:** Bio-logging tags are frequently programmed with intermittent duty cycles or sleep schedulers to conserve battery. Subtracting the session’s final timestamp from its initial timestamp would erroneously report sleep intervals as active data. The reporter bypasses this by calculating the true duration from the extracted arrays 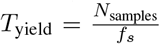. For instance, 47 min 12 s of continuous IMU data sampled at 50 Hz corresponds to exactly 141,600 rows in the output CSV, providing a precise scientific yield.
4. **Errors and Warnings:** The module logs exact file paths for corrupted binary payloads and gracefully catches missing external dependencies. Furthermore, GPS success is reported mathematically as *η*_GPS_ = *N*_fixes_*/N*_attempts_. Low percentage values commonly occur under dense canopy or in mountainous terrain and reflect physical field conditions rather than software faults.

To support execution on headless environments, the reporter implements a simple console truncation. When processing massive datasets, it limits standard terminal output to approximately a few dozen lines to prevent console spam, while ensuring that all CRITICAL errors and WARN logs bypass the truncation and remain visible.

### Modification and Extension

The software is released under GPLv3 to encourage community contributions. Apart from its application to decode data recorded with Vesper Wildlife tags, the `binary_decoder.py` can be adapted for other proprietary binary formats used in wildlife telemetry. The decoupled nature of the parsers allows developers to add new sensor types (e.g., humidity) by simply adding a new parser class and updating the file scanner. Contributors can reach the authors via the GitHub repository or the provided email addresses.

Support is provided via the GitHub *Issues* tab for bug reporting and feature requests. Comprehensive technical documentation and a user manual are included in the repository.

### Case study: ring-tailed lemurs (*Lemur catta*)

#### Data collection

We deployed collars containing Vesper Wildlife tags on 10 ring-tailed lemurs (*Lemur catta*) at Affenwald Straußberg, Germany, as part of a separate research project. Ethical approval (E1-26) was issued by the Animal Welfare Committee and the Animal Welfare Officer at the German Primate Center, which is registered and authorised by the local and regional veterinary governmental authorities (Ref. no. 122910.3311900, PK 36674). Because the study population was habituated to human presence, collars could be fitted without capture or anesthesia. A food lure was used to guide each animal into position, after which the collar was quickly secured. Following a deployment of 3 days, collars were removed by approaching the animals with a food lure and releasing the snap fastener. This procedure enabled rapid deployment and retrieval while minimizing handling and disturbance to the animals.

#### Case and collar

The tag housing and collar were designed as a single, integrated system consisting of two main components secured by four interlocking keys, with a snap fastener used to close the collar loop. Constructed primarily from flexible yet durable thermoplastic polyurethane (TPU; collar and cap) with reinforcing polylactic acid (PLA) or Acrylonitrile Styrene Acrylate (ASA; case and keys) components, the system is lightweight, streamlined, and comfortable for the animal. The tag (and battery) case measures 55 × 35 mm and is sized to fit a 18500 Li-Ion battery. The design is weather-resistant and has performed reliably under variable field conditions, with no instances of loss or failure during deployments lasting up to one week. The complete 3D model file, ready for printing, is provided in the Supplementary Materials.

#### Tag settings

The GPS unit was configured to search for satellites with a snap size of 512 ms and a sample rate of 1500 ms. The audio recorder was set to a sampling rate of 48,000 Hz with a file size of 125,829,120 bytes (approximately 22.5 minutes). A digital filter was enabled to reduce background noise, and the programmable gain was set to 30.1 dB. The IMU was configured to sample at a rate of 50 Hz. The gyroscope and accelerometer, as well as the barometer were enabled, and the accelerometer and gyroscope dynamic ranges were set to ±2g and ±250 dps, respectively, which is suitable for most terrestrial animals. The exact settings and tag deployment protocols can be found in the Supplementary Materials.

The Proxit tag is a standalone, Bluetooth-based proximity sensor that shares a battery with the Vesper Wildlife tag but operates independently. Tag configuration and data retrieval were managed using a custom Python script (see Supplementary Materials). When a signal from another Proxit tag is detected, a new interaction record is created if either (1) more than 18 min has elapsed since the most recent record involving the same tag or (2) the received signal strength indicator (RSSI) has changed by more than 10 dB, providing a proxy for a change in distance between individuals. Bluetooth transmission power was set to -3 dBm (decibel-milliwatts). In our test environment, this setting enabled the detection of interactions at distances of up to approximately 10 m, with greater sensitivity to distance changes within 5 m. Data were downloaded using a Thunderboard EFR32BG22 dongle connected to a Windows tablet. The custom Python script functions as a Network Co-Processor host application, scanning for nearby Proxit devices, establishing Bluetooth connections, and checking each device for stored records. The effective connection range was approximately 30 m, although this varied with environmental conditions. During each connection, the script synchronized the device clock to Coordinated Universal Time (UTC) to support temporal alignment with the Vesper data. Available records, including device identifiers, interaction status, RSSI, and timestamps, were then downloaded and written to a CSV file. Each record was verified as saved before the next record was retrieved, reducing the risk of data loss if an animal moved out of range during transfer. Successfully transferred records were subsequently deleted from the device. Each Proxit tag can store up to 448 interaction records. Where animals range over a large sampling area, multiple download hubs may therefore be required at suitable locations to support regular data retrieval and prevent the device memory from reaching capacity.

#### Results

Data were successfully recovered from all 10 deployed collars. The collars remained attached throughout deployment, with no observed loss or structural failure under variable field conditions. Vesper Wildlife tag files were downloaded and processed using VesperFlow, which successfully decoded the GPS, audio, and IMU data. Interaction records from the Proxit tags were downloaded separately and temporally aligned with the Vesper data using synchronized UTC timestamps. The resulting four data types, GPS locations, audio recordings, IMU measurements, and proximity interactions, were visualized for inspection (Figure 4). The decoded data showed the expected structure and temporal patterns for each sensor type, and no apparent decoding or synchronization errors were identified. This case study therefore demonstrates that VesperFlow can be used to process multimodal Vesper Wildlife tag data collected during field deployments on free-ranging, ring-tailed lemurs. With appropriate adjustments to collar dimensions, device mass, and species-specific welfare requirements, the tag can be readily adapted for use with other medium- to large-bodied mammals.

**Fig. 4.**
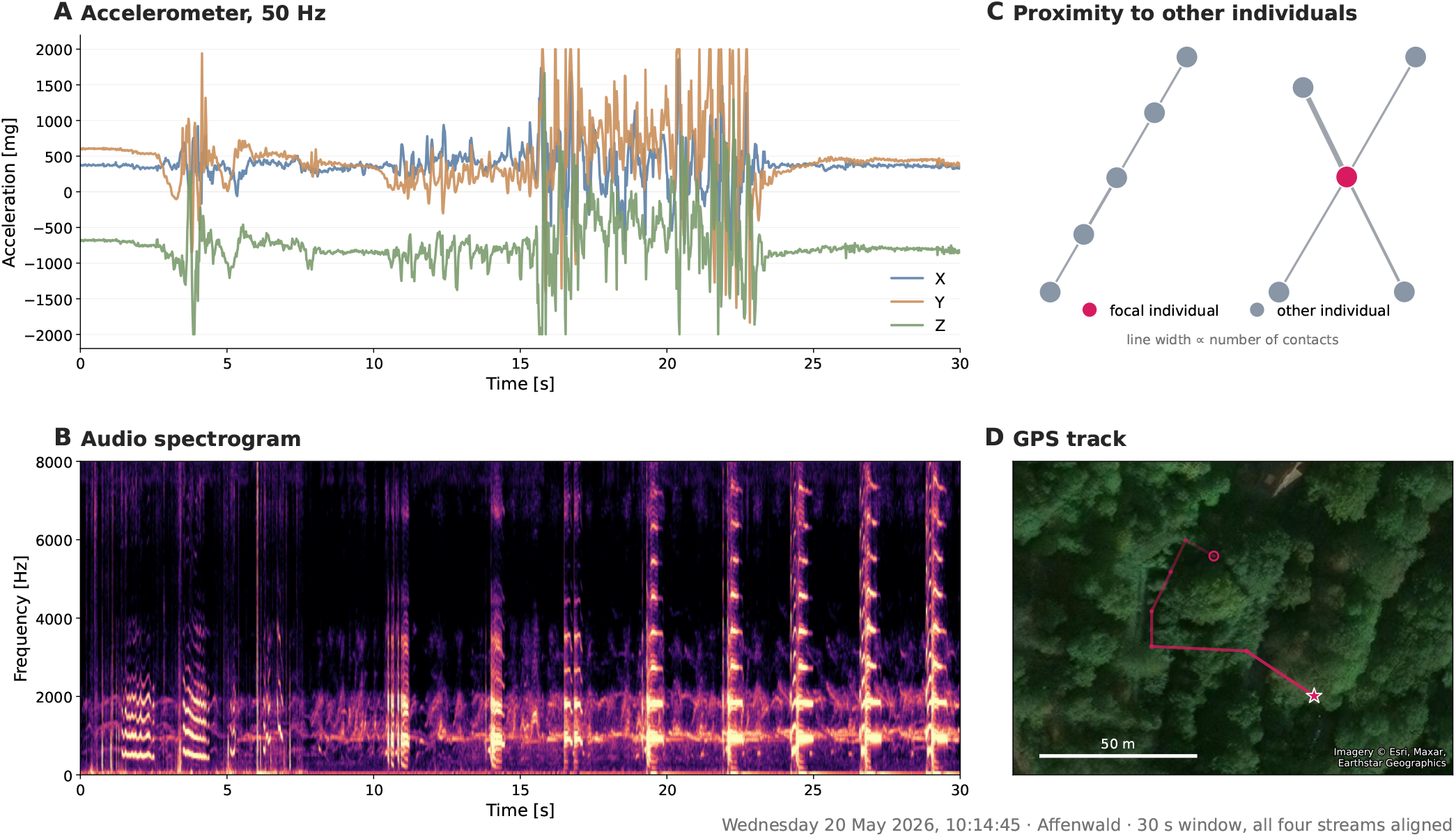
Representative outputs from animal-mounted tags deployed on ring-tailed lemurs (*Lemur catta*). A) Tri-axial accelerometer data, B) Spectrogram derived from the acoustic recordings, C) Close-range interactions between tagged individuals recorded by the Proxit sensors, D) GPS tracks showing the recorded movements of tagged individuals. Vesper Wildlife tag data were decoded and processed using VesperFlow, while Proxit data were downloaded and processed separately.

#### Validation and benchmarking

Functional validation was achieved by benchmarking the tool’s outputs directly against those produced by the proprietary manufacturer tool (Vesper-App). Using real-world, multi-modal field datasets, the extracted sensor arrays were cross-examined. The tool achieves 100% parity in data extraction while actively improving data quality by mitigating hardware artifacts (e.g., gyroscope file-split bug and strictly compliant RIFF WAV headers).

Furthermore, because the software utilizes in-memory vectorized array operations (via `NumPy` and `pandas`) rather than the proprietary tool’s multi-phase intermediate file approach (`.MBN`/`.UBN` disk writes), it processes large batch deployments significantly faster and with drastically reduced disk I/O bottlenecks.

## ACKNOWLEDGEMENTS

We thank Alexander Schwartz and the Alexander Schwartz Developments team for their support in addressing technical questions about the Vesper Wildlife tag system and VesperApp. We are also grateful to Silvio Dietzel and Affenwald, Straussberg staff for providing access to the animals.

Funding for the project was supported by a Deutsche Forschungsgemeinschaft grant (WI 6390/2-1 to KW).

