## Supplementary Materials for "VesperFlow: An automated pipeline for decoding multisensor data from biologging tags"

### S1. Tag settings and configuration

#### S1.1. Vesper Wildlife Tag description

The Vesper Wildlife tagging system is a modular system, with various sensors that can be added or removed. For the purpose of this research, we used a system containing the main unit (with GPS and an audio sensor, `https://asd-tech.com/product/vt04-vesper-64/`), an IMU unit (`https://asd-tech.com/product/vt04-imu10`), and a Proxtit unit (`https://asd-tech.com/product/proxtit/`). The main unit and IMU unit are both operated via the VesperApp (Version 1.0.27.0, ASD, n.d.), for which the tag needs to be connected to a laptop via a docking station (`https://asd-tech.com/product/vt04-dockv4/`), and are controlled using a configuration file that is created in the VesperApp. The proxtit tag only takes energy from the battery but is a completely standalone unit in everything else. It is controlled with Python code and connects via the dongle (Thunderboard EFR32BG22).

#### S1.2. General tag functioning

The tag can be operated with a magnet. It has four modes, which are shown using light flashes: (1) idle (continuous blinking), allows communication with the docking station and VesperApp; (2) armed sleep (two long slow blinks), ready for switch to sampling state; (3) sampling state (one blink for wake-up, then 2 blinks for start sampling), executing configuration file; (4) non-armed sleep (one long blink), reserves battery. Errors or issues are also shown using blinks. One long blink, then multiple rapid blinks indicates either a battery issue or a missing configuration file.

#### S1.3. Tag configuration

The Vesper tag was configured to record GPS location, audio, and motion data according to a pre-defined schedule. All settings were managed through the configuration file (config.json, see below), which is created in the VesperApp and uploaded to the tag. Using the settings below, the Vesper tag with a storage of 58 GB is able to collect data for 91 hours before the storage is fully used. The following settings were used for the configuration.

##### S1.3.1. GPS (ACLYS)

The GPS unit was configured to search for satellites with a snap size of 512 ms and a sample rate of 1500 ms. The file size was set to its default size, being 1,000,000 bytes. These settings were chosen to maximize the frequency and stability of location fixes in a forested environment, where satellite connection is limited.

##### S1.3.2. Audio (SPH0641)

The audio recorder was set to a sampling rate of 48,000 Hz with a file size of 125,829,120 bytes (approximately 22.5 minutes). A digital filter was enabled to reduce background noise, and the programmable gain was set to 30.1 dB. The high-pass filter was disabled, as this is only necessary for ultrasonic recordings (e.g., bats).

##### S1.3.3. Motion (IMU10)

The IMU was configured to sample at a rate of 50 Hz. The gyroscope and accelerometer, as well as the barometer, were enabled, and the accelerometer and gyroscope dynamic ranges were set to ±2g and ±250 dps, respectively, which is suitable for most terrestrial animals.

##### S1.3.4. Proxtit

The Proxtit tag was used to record close-range interactions between individuals, with a maximum range of 10 meters. The tag operates by continuously listening for Bluetooth signals from other Proxtit tags. Upon detecting a signal, it records a new interaction if (1) more than eighteen minutes have elapsed since the last recorded interaction with that specific tag, or (2) the signal strength (RSSI) changes by more than 10 decibels, indicating a significant change in the relative distance between the individuals. The proxtit tags have storage capacity for up to 448 logs, i.e. connections to other tags. The system was set to a TX power of -3 to detect close-range interactions of up to 10 meters, with higher resolution within 5 meters.

**Table S1.** Configuration file (`config.json`) uploaded to all Vesper tags. Line breaks were added for readability.


```
{
  "name": "Vesper",
  "minhw": "4.0",
  "cdrift": 32999,
  "scheduleType": 0,
  "magnetOff": true,
  "battery": 60,
  "poweron": "",
  "schedule": [
    {"time": "2000-08-25 03:30:00", "config": 1},
    {"time": "2000-08-25 19:00:00", "config": 0}
  ],
  "devices": [
    {"name": "ACLYS", "sampleRate": [0, 1500, 0], "wLen": [0, 0, 0], "wRate": [0, 0, 0], "bitmask": 2,
        "snapSize": 512, "fileSize": 0},
    {"name": "SPH0641", "sampleRate": [0, 48000, 0], "wLen": [0, 0, 0], "wRate": [0, 0, 0], "bitmask":
        0, "fileSize": 125829120, "thresup": 0, "thresdn": 0, "cic4": false, "filter": 257, "hpf": 0,
        "gain": 10},
    {"name": "LED", "sampleRate": [0, 0, 0], "wLen": [0, 0, 0], "wRate": [0, 0, 0], "bitmask": 0,
        "fileSize": 0},
    {"name": "IMU10", "sampleRate": [0, 50, 0], "wLen": [0, 0, 0], "wRate": [0, 0, 0], "bitmask": 15,
        "imuRange": 1, "fileSize": 524288}
  ]
}
```


### S2. Deployment protocol

The deployment is performed according to the following sequence:

#### S2.1. Preparation of the tags

1. Open the VesperApp, connect a docking station to the computer and attach it to the VesperApp using the ‘Connect Dock’ button. Then click ‘Enable Device’ so that the circle turns green.

2. Connect the tag to the docking station with the wires and sensor units facing forward, and switch it to idle using a magnet. It will show up on the VesperApp. Click the tag on the VesperApp and click ‘Connect Device’. Now the software version and current time on the tag can be seen.

3. Before deployment, format each tag using the format function in File Explorer and the default settings. This reduces clock drift and removes faulty data writes.

4. Re-upload the config.json file to the tag, and recalibrate the tag by pressing ‘Set Time’ in the VesperApp. Make sure the app is set to UTC.

5. To decouple the tag, press ‘Arm Sleep’. The tag will blink longer a couple of times and then the light turns off, indicating the tag is ready to be deployed.

#### S2.2. Deployment and retrieval procedure

1. Turn on tags by passing the magnet over the tag once; it will blink three times upon turn-on (one blink for wake-up, then 2 blinks for start sampling), then blink continuously with one-second intervals while recording data.

   (a) If the tag blinks rapidly and then turns off, this signals an issue in the configuration or with the battery. Configuration issues can be resolved by re-uploading the configuration file to the tag or formatting the tag. Connect the tags to the laptop again and reupload the config file. In case of battery issues, check for wire breaks or oxidation of the battery, and repair as needed.

2. Close the case by inserting the four pins.

3. Manually do a time calibration in the range of all tags (explained below).

4. Using a food lure, distract the animal. Then, another person attaches the collar to the animal’s neck by closing the snap fastenings. The zigzag pattern should be facing up.

5. Do a time calibration daily.

6. Retrieve the collar after a maximum of 4 days, as the storage will be full after 91 hours of active sampling. Do this using a food lure and unclipping the snap fastening. Manually do a time calibration in the range of all tags.

7. Switch each tag to non-armed sleep by passing the magnet over the tag once; the tag will blink once for two seconds.

8. Keep track of the date and time the collar was attached and removed from the animal, and note down the Vesper ID and the animal ID.

#### S2.3. Proximity system setup

Data from the Proxtit tags were downloaded continuously using a dongle (Thunderboard EFR32BG22) and a custom Python script (see Supplementary Material S5). The script was run from a Windows tablet, which activated the dongle to scan for and connect to all available Proxtit tags within a range of about 30 meters, depending on the environment. We placed two tablets with dongles in the forest to ensure good download coverage of the area.

1. Place each Proxtit hub in a strategic area and attach the dongle using a USB cable.

2. Open Command Prompt

3. Call: "py" and then direct the code to the app.py file. E.g.: `py ~/proxtit-downloader-py-master/downloader/app.py`. Press Enter.

4. The program will connect to the dongle, then start scanning for nearby tags and start downloading and removing their data entries. After the connection to one tag is closed, it automatically starts searching for another tag.

5. It is good practice to download the data from each tag before deployment, to ensure their storage is empty before use.

### S3. Time calibration

To test clock drift and align the different tags temporally, we performed daily manual time calibrations using two mobile apps: TalkingTime (version 1.6.0, https://play.google.com/store/apps/details?id=apps.sadim.talking_time) and Atomic Clock (version 2.1.22, https://apps.microsoft.com/detail/9wzdncrfj0cf?hl=nl-NL&gl=DE).

1. In TalkingTime, turn on Background mode and Periodic Announcements. Set the interval to every minute.

2. In Atomic Clock, turn on Acoustic ticking and set the Greenwich time signal to every minute. This will allow for an acoustic time signal each minute: five beeps at the last five seconds of a minute, with the last beep having a longer duration.

3. With maximum volume, and while keeping the Atomic clock open, make sure every tag is within hearing range of the Greenwich signal and follows the acoustic time announcement at least once.

4. Keep a record of the time period and date of the time calibration.

### S4. Proxtit code

The app.py script is an NCP host application that scans for PRX devices using Bluetooth, connects to them and checks for available logs on the device. It then synchronises the device clock to UTC to ensure temporal alignment with the Vesper tag data, and sets the device Bluetooth transmission power to TX=-3dBm. Then, all available logs, including the status, device IDs, signal strength, and interaction timestamps, are downloaded and written into a .csv file, after which it is deleted from the device. The script ensures that each log is saved to the .csv file before another is downloaded, to not lose data when the animal moves out of range.

The app.py script works together with the util.py script, which manages device connections, initializes Bluetooth and Bluetooth Mesh applications, handles errors and monitors device responsiveness. It also encodes commands that configure device settings, such as the real-time clock and transmission power, and decodes raw byte data from device logs into readable information.

We altered both the original app.py and util.py source code provided by Silicon Laboratories Inc. to meet our specific requirements. The complete folder structure needed for the code is provided alongside this paper (proxtit-downloader-py-master).
