## Supplementary material for "VesperFlow: An automated pipeline for decoding multisensor data from biologging tags": CHANGELOG.md

# Changelog

The releases in this repo are based on particular [Gecko SDK](https://github.com/SiliconLabs/gecko_sdk/releases)
and [Simplicity SDK](https://github.com/SiliconLabs/simplicity_sdk/releases) releases.
The version numbers here reflect the GSDK/SiSDK version that the release is based on.

## [2024.12.0] - 2024-12-19

### Changed
- Update BGAPI version to 9.0.0.
- Keep SL status codes aligned with the SiSDK

### Fixed
- Indication not stopped after connection close in the bt_thermometer example

## [2024.6.0] - 2024-06-10

### Changed
- Update BGAPI version to 8.0.0.
- Keep SL status codes aligned with the SiSDK (former GSDK)
- Replace the monolithic `event_handler` method with dedicated event handler methods in all examples
- Refactor connection handling in the bt_thermometer_client example

### Fixed
- Various GUI issues in mesh light and switch examples on Mac and Linux
- Most of the pylint errors in mesh examples

## [4.4.0] - 2023-12-13

### Changed
- Device reset is not connector type dependent anymore, it is performed by a watchdog at startup.
- Update BGAPI version to 7.0.0.
- Represent BGAPI event and response parameters with `errorcode` type as `Status` objects with human readable string representation

### Added
- Robust connector option. Requires at least pyBGAPI v1.3.0.
- New features in bt_roaming example: RSSI sniffing, network visualization
- NCP test example
- Module with SL status values

### Removed
- CPC connector has been moved to the pyBGAPI package. Requires at least pyBGAPI v1.3.0.

## [4.3.0] - 2023-06-09

### Fixed
- Various issues with CPC reset

### Changed
- Update BGAPI version to 6.0.0.

### Removed
- libcpc_wrapper.py. Install libcpc module from the [cpc-daemon](https://github.com/SiliconLabs/cpc-daemon) repo.

## [4.2.0] - 2022-12-15

### Added
- bt_roaming example

### Changed
- Update BGAPI version to 5.0.0.
- `GenericApp` class derived from `threading.Thread`, therefore it's easier to
  start `GenericApp` instances in their own thread.
- Events can be handled in their dedicated event callback methods in addition to
  the generic `event_handler` method of the `GenericApp`.
- `ConnectorApp` class removed, its functionality moved to `ArgumentParser` class
  and`get_connector` functions.
- `BluetoothApp` and `BtmeshApp` classes derived directly from `GenericApp`,
  they need a connector instance in their constructor.
- `PeriodicTimer` replaced with threading objects in `bt_thermometer` example.
- Timestamp added to logging messages.
- Libcpc_wrapper script updated.

### Removed
- `PeriodicTimer` helper class

## [4.1.2] - 2022-09-30
### Added
- btmesh_light and btmesh_switch examples.
- CPC support (Linux only).
- 'bt_' prefix for BLE example names.
- Readme files in the example folders.
- This change log.

### Changed
- Update BGAPI version to 4.2.0.

### Fixed
- Float conversion issue with bt_thermometer_client example.

## [4.0.0] - 2021-12-20
### Changed
- Update BGAPI version to 3.3.0.
- Node reset removed from Bluetooth mesh reset.

## 3.2.2 - 2021-09-14
### Added
- Initial public release.

[2024.12.0]: https://github.com/SiliconLabs/simplicity_sdk/releases/tag/v2024.12.0
[2024.6.0]: https://github.com/SiliconLabs/simplicity_sdk/releases/tag/v2024.6.0
[4.4.0]: https://github.com/SiliconLabs/gecko_sdk/releases/tag/v4.4.0
[4.3.0]: https://github.com/SiliconLabs/gecko_sdk/releases/tag/v4.3.0
[4.2.0]: https://github.com/SiliconLabs/gecko_sdk/releases/tag/v4.2.0
[4.1.2]: https://github.com/SiliconLabs/gecko_sdk/releases/tag/v4.1.2
[4.0.0]: https://github.com/SiliconLabs/gecko_sdk/releases/tag/v4.0.0
