## Supplementary material for "VesperFlow: An automated pipeline for decoding multisensor data from biologging tags": btmesh_models.py

```
"""
Module for bluetooth mesh models
"""

# Copyright 2024 Silicon Laboratories Inc. www.silabs.com
#
# SPDX-License-Identifier: Zlib
#
# The licensor of this software is Silicon Laboratories Inc.
#
# This software is provided 'as-is', without any express or implied
# warranty. In no event will the authors be held liable for any damages
# arising from the use of this software.
#
# Permission is granted to anyone to use this software for any purpose,
# including commercial applications, and to alter it and redistribute it
# freely, subject to the following restrictions:
#
# 1. The origin of this software must not be misrepresented; you must not
#    claim that you wrote the original software. If you use this software
#    in a product, an acknowledgment in the product documentation would be
#    appreciated but is not required.
# 2. Altered source versions must be plainly marked as such, and must not be
#    misrepresented as being the original software.
# 3. This notice may not be removed or altered from any source distribution.

BTMESH_CONF_CONFIG_SERVER_MODEL_ID = 0x0000
BTMESH_CONF_CONFIG_CLIENT_MODEL_ID = 0x0001
BTMESH_CONF_HEALTH_SERVER_MODEL_ID = 0x0002
BTMESH_CONF_HEALTH_CLIENT_MODEL_ID = 0x0003
BTMESH_REMOTE_PROV_SERVER_MODEL_ID = 0x0004
BTMESH_REMOTE_PROV_CLIENT_MODEL_ID = 0x0005
BTMESH_DIRECTED_FORWARD_CONF_SERVER_MODEL_ID = 0x0006
BTMESH_DIRECTED_FORWARD_CONF_CLIENT_MODEL_ID = 0x0007
BTMESH_BRIDGE_CONF_SERVER_MODEL_ID = 0x0008
BTMESH_BRIDGE_CONF_CLIENT_MODEL_ID = 0x0009
BTMESH_PRIVATE_BEACON_SERVER_MODEL_ID = 0x000A
BTMESH_PRIVATE_BEACON_CLIENT_MODEL_ID = 0x000B
BTMESH_ON_DEMAND_PRIVATE_PROXY_SERVER_MODEL_ID = 0x000C
BTMESH_ON_DEMAND_PRIVATE_PROXY_CLIENT_MODEL_ID = 0x000D
BTMESH_SAR_CONFIG_SERVER_MODEL_ID = 0x000E
BTMESH_SAR_CONFIG_CLIENT_MODEL_ID = 0x000F
BTMESH_OPCODE_AGGREGATOR_SERVER_MODEL_ID = 0x0010
BTMESH_OPCODE_AGGREGATOR_CLIENT_MODEL_ID = 0x0011
BTMESH_LCD_SERVER_MODEL_ID = 0x0012
BTMESH_LCD_CLIENT_MODEL_ID = 0x0013
BTMESH_SOLICITATION_PDU_RPL_CONFIG_SERVER_MODEL_ID = 0x0014
BTMESH_SOLICITATION_PDU_RPL_CONFIG_CLIENT_MODEL_ID  = 0x0015
BTMESH_GENERIC_ON_OFF_SERVER_MODEL_ID = 0x1000
BTMESH_GENERIC_ON_OFF_CLIENT_MODEL_ID = 0x1001
BTMESH_GENERIC_LEVEL_SERVER_MODEL_ID = 0x1002
BTMESH_GENERIC_LEVEL_CLIENT_MODEL_ID = 0x1003
BTMESH_GENERIC_TRANSITION_TIME_SERVER_MODEL_ID = 0x1004
BTMESH_GENERIC_TRANSITION_TIME_CLIENT_MODEL_ID = 0x1005
BTMESH_GENERIC_POWER_ON_OFF_SERVER_MODEL_ID = 0x1006
BTMESH_GENERIC_POWER_ON_OFF_SETUP_SERVER_MODEL_ID = 0x1007
BTMESH_GENERIC_POWER_ON_OFF_CLIENT_MODEL_ID = 0x1008
BTMESH_GENERIC_POWER_LEVEL_SERVER_MODEL_ID = 0x1009
BTMESH_GENERIC_POWER_LEVEL_SETUP_SERVER_MODEL_ID = 0x100A
BTMESH_GENERIC_POWER_LEVEL_CLIENT_MODEL_ID = 0x100B
BTMESH_GENERIC_BATTERY_SERVER_MODEL_ID = 0x100C
BTMESH_GENERIC_BATTERY_CLIENT_MODEL_ID = 0x100D
BTMESH_GENERIC_LOCATION_SERVER_MODEL_ID = 0x100E
BTMESH_GENERIC_LOCATION_SETUP_SERVER_MODEL_ID = 0x100F
BTMESH_GENERIC_LOCATION_CLIENT_MODEL_ID = 0x1010
BTMESH_GENERIC_ADMIN_PROPERTY_SERVER_MODEL_ID = 0x1011
BTMESH_GENERIC_MANUF_PROPERTY_SERVER_MODEL_ID = 0x1012
BTMESH_GENERIC_USER_PROPERTY_SERVER_MODEL_ID = 0x1013
BTMESH_GENERIC_CLIENT_PROPERTY_SERVER_MODEL_ID = 0x1014
BTMESH_GENERIC_PROPERTY_CLIENT_MODEL_ID = 0x1015
BTMESH_SENSOR_SERVER_MODEL_ID = 0x1100
BTMESH_SENSOR_SETUP_SERVER_MODEL_ID = 0x1101
BTMESH_SENSOR_CLIENT_MODEL_ID = 0x1102
BTMESH_TIME_SERVER_MODEL_ID = 0x1200
BTMESH_TIME_SETUP_SERVER_MODEL_ID = 0x1201
BTMESH_TIME_CLIENT_MODEL_ID = 0x1202
BTMESH_SCENE_SERVER_MODEL_ID = 0x1203
BTMESH_SCENE_SETUP_SERVER_MODEL_ID = 0x1204
BTMESH_SCENE_CLIENT_MODEL_ID = 0x1205
BTMESH_SCHEDULER_SERVER_MODEL_ID = 0x1206
BTMESH_SCHEDULER_SETUP_SERVER_MODEL_ID = 0x1207
BTMESH_SCHEDULER_CLIENT_MODEL_ID = 0x1208
BTMESH_LIGHTING_LIGHTNESS_SERVER_MODEL_ID = 0x1300
BTMESH_LIGHTING_LIGHTNESS_SETUP_SERVER_MODEL_ID = 0x1301
BTMESH_LIGHTING_LIGHTNESS_CLIENT_MODEL_ID = 0x1302
BTMESH_LIGHTING_CTL_SERVER_MODEL_ID = 0x1303
BTMESH_LIGHTING_CTL_SETUP_SERVER_MODEL_ID = 0x1304
BTMESH_LIGHTING_CTL_CLIENT_MODEL_ID = 0x1305
BTMESH_LIGHTING_CTL_TEMPERATURE_SERVER_MODEL_ID = 0x1306
BTMESH_LIGHTING_HSL_SERVER_MODEL_ID = 0x1307
BTMESH_LIGHTING_HSL_SETUP_SERVER_MODEL_ID = 0x1308
BTMESH_LIGHTING_HSL_CLIENT_MODEL_ID = 0x1309
BTMESH_LIGHTING_HSL_HUE_SERVER_MODEL_ID = 0x130A
BTMESH_LIGHTING_HSL_SATURATION_SERVER_MODEL_ID = 0x130B
BTMESH_LIGHTING_XYL_SERVER_MODEL_ID = 0x130C
BTMESH_LIGHTING_XYL_SETUP_SERVER_MODEL_ID = 0x130D
BTMESH_LIGHTING_XYL_CLIENT_MODEL_ID = 0x130E
BTMESH_LC_SERVER_MODEL_ID = 0x130F
BTMESH_LC_SETUP_SERVER_MODEL_ID = 0x1310
BTMESH_LC_CLIENT_MODEL_ID = 0x1311
BTMESH_MBT_SERVER_MODEL_ID = 0x1400
BTMESH_MBT_CLIENT_MODEL_ID = 0x1401
BTMESH_DFU_SERVER_MODEL_ID = 0x1402
BTMESH_DFU_CLIENT_MODEL_ID = 0x1403
BTMESH_DFU_DISTRIBUTOR_SERVER_MODEL_ID = 0x1404
BTMESH_DFU_DISTRIBUTOR_CLIENT_MODEL_ID = 0x1405
```
