## Supplementary material for "VesperFlow: An automated pipeline for decoding multisensor data from biologging tags": conversion.py

```
"""
Module for data conversion classes
"""

# Copyright 2021 Silicon Laboratories Inc. www.silabs.com
#
# SPDX-License-Identifier: Zlib
#
# The licensor of this software is Silicon Laboratories Inc.
#
# This software is provided 'as-is', without any express or implied
# warranty. In no event will the authors be held liable for any damages
# arising from the use of this software.
#
# Permission is granted to anyone to use this software for any purpose,
# including commercial applications, and to alter it and redistribute it
# freely, subject to the following restrictions:
#
# 1. The origin of this software must not be misrepresented; you must not
#    claim that you wrote the original software. If you use this software
#    in a product, an acknowledgment in the product documentation would be
#    appreciated but is not required.
# 2. Altered source versions must be plainly marked as such, and must not be
#    misrepresented as being the original software.
# 3. This notice may not be removed or altered from any source distribution.

import decimal

class Ieee11073Float(float):
    '''IEEE 11073 32 bit float converter class.'''
    @classmethod
    def from_bytes(cls, src):
        '''Return the IEEE 11073 32 bit float represented by the given array of bytes.'''
        mantissa = int().from_bytes(src[0:3], byteorder='little', signed=True)
        exponent = int().from_bytes([src[3]], byteorder='little', signed=True)
        val = mantissa * pow(10, exponent)
        return float(val)

    def to_bytes(self):
        '''Return an array of bytes representing an IEEE 11073 32 bit float.'''
        (sign, digits, exponent) = decimal.Decimal(str(self)).as_tuple()
        # Convert digits to a mantissa string
        mantissa = ''.join([str(i) for i in digits])
        # Remove trailing zeros
        if len(mantissa) > 1:
            mantissa = mantissa.rstrip('0')
        # Adjust exponent according to removed zeros
        exponent = exponent + (len(digits) - len(mantissa))
        # Convert mantissa to integer
        mantissa = int(mantissa)
        if sign == 1:
            mantissa = -1 * mantissa
        return mantissa.to_bytes(3, byteorder='little', signed=True) + exponent.to_bytes(1, byteorder='little', signed=True)
```
