## Supplementary material for "VesperFlow: An automated pipeline for decoding multisensor data from biologging tags": status.py

```
#!/usr/bin/env python3
"""
SL status codes derived from sl_status.h
"""

# Copyright 2023 Silicon Laboratories Inc. www.silabs.com
#
# SPDX-License-Identifier: Zlib
#
# The licensor of this software is Silicon Laboratories Inc.
#
# This software is provided 'as-is', without any express or implied
# warranty. In no event will the authors be held liable for any damages
# arising from the use of this software.
#
# Permission is granted to anyone to use this software for any purpose,
# including commercial applications, and to alter it and redistribute it
# freely, subject to the following restrictions:
#
# 1. The origin of this software must not be misrepresented; you must not
#    claim that you wrote the original software. If you use this software
#    in a product, an acknowledgment in the product documentation would be
#    appreciated but is not required.
# 2. Altered source versions must be plainly marked as such, and must not be
#    misrepresented as being the original software.
# 3. This notice may not be removed or altered from any source distribution.

import os.path
import re
import sys

class Status(int):
    """Representation for SL status codes"""
    def __new__(cls, value, base=0):
        if isinstance(value, str):
            value = value.upper()
            if value.startswith("SL_STATUS_"):
                # remove prefix
                value = value[len("SL_STATUS_"):]
            if value in globals():
                value = globals()[value]
            else:
                # Try to convert string value to a valid integer. Setting the default base to 0
                # allows to detect the base automatically based on an optional prefix. Note that
                # the int constructor uses base=10 by default.
                value = int(value, base)
        return super().__new__(cls, value)

    def __repr__(self):
        names = [var for var in globals() if var.isupper()]
        for name in names:
            if globals()[name] == self:
                return f"SL_STATUS_{name}"
        return f"SL_STATUS_{self:#06x}"

    def __str__(self):
        if self in _doc:
            return _doc[self]
        return f"Unknown status: {self:#06x}"

def compare(header):
    """Compare status values in this module to the status values in the header"""
    header_values = set()
    pattern = r"#define SL_STATUS_(?P<name>\w+)\s+\(\(sl_status_t\)(?P<value>0x[0-9A-Fa-f]+)\)\s+///< (?P<doc>.+)"
    with open(header, encoding="utf-8") as hdr:
        parse = False
        for line in hdr:
            if not parse:
                # Skip space defines
                if "Status Defines" in line:
                    parse = True
                continue
            match = re.match(pattern, line)
            if not match:
                continue
            value = int(match.group("value"), base=0)
            header_values.add((match.group("name"), value, match.group("doc")))
    local_values = {(name, int(value), _doc[value]) for name, value in globals().items() if name.isupper()}
    diff = local_values ^ header_values
    if diff:
        # Sort difference based on the integer representation (2nd item in the tuple)
        diff = sorted(diff, key=lambda x: x[1])
        for d in diff:
            where = "+" if d in local_values else "-"
            print(where, d)
    else:
        print("Header and local values are identical.")

def main():
    """Module test"""
    if len(sys.argv) != 2:
        print("Usage:")
        print(f"{sys.argv[0]} <value>               - Interpret <value> as SL status.")
        print(f"{sys.argv[0]} <path/to/sl_status.h> - Compare status values in the input header to the status values in this module.")
    else:
        if os.path.exists(sys.argv[1]):
            compare(sys.argv[1])
        else:
            s = Status(sys.argv[1])
            print(f"{repr(s)}: {s}")

# Generic Errors
OK = Status(0x0000)
"""No error."""
FAIL = Status(0x0001)
"""Generic error."""

# State Errors
INVALID_STATE = Status(0x0002)
"""Generic invalid state error."""
NOT_READY = Status(0x0003)
"""Module is not ready for requested operation."""
BUSY = Status(0x0004)
"""Module is busy and cannot carry out requested operation."""
IN_PROGRESS = Status(0x0005)
"""Operation is in progress and not yet complete (pass or fail)."""
ABORT = Status(0x0006)
"""Operation aborted."""
TIMEOUT = Status(0x0007)
"""Operation timed out."""
PERMISSION = Status(0x0008)
"""Operation not allowed per permissions."""
WOULD_BLOCK = Status(0x0009)
"""Non-blocking operation would block."""
IDLE = Status(0x000A)
"""Operation/module is Idle, cannot carry requested operation."""
IS_WAITING = Status(0x000B)
"""Operation cannot be done while construct is waiting."""
NONE_WAITING = Status(0x000C)
"""No task/construct waiting/pending for that action/event."""
SUSPENDED = Status(0x000D)
"""Operation cannot be done while construct is suspended."""
NOT_AVAILABLE = Status(0x000E)
"""Feature not available due to software configuration."""
NOT_SUPPORTED = Status(0x000F)
"""Feature not supported."""
INITIALIZATION = Status(0x0010)
"""Initialization failed."""
NOT_INITIALIZED = Status(0x0011)
"""Module has not been initialized."""
ALREADY_INITIALIZED = Status(0x0012)
"""Module has already been initialized."""
DELETED = Status(0x0013)
"""Object/construct has been deleted."""
ISR = Status(0x0014)
"""Illegal call from ISR."""
NETWORK_UP = Status(0x0015)
"""Illegal call because network is up."""
NETWORK_DOWN = Status(0x0016)
"""Illegal call because network is down."""
NOT_JOINED = Status(0x0017)
"""Failure due to not being joined in a network."""
NO_BEACONS = Status(0x0018)
"""Invalid operation as there are no beacons."""

# Allocation/ownership Errors
ALLOCATION_FAILED = Status(0x0019)
"""Generic allocation error."""
NO_MORE_RESOURCE = Status(0x001A)
"""No more resource available to perform the operation."""
EMPTY = Status(0x001B)
"""Item/list/queue is empty."""
FULL = Status(0x001C)
"""Item/list/queue is full."""
WOULD_OVERFLOW = Status(0x001D)
"""Item would overflow."""
HAS_OVERFLOWED = Status(0x001E)
"""Item/list/queue has been overflowed."""
OWNERSHIP = Status(0x001F)
"""Generic ownership error."""
IS_OWNER = Status(0x0020)
"""Already/still owning resource."""

# Invalid Parameters Errors
INVALID_PARAMETER = Status(0x0021)
"""Generic invalid argument or consequence of invalid argument."""
NULL_POINTER = Status(0x0022)
"""Invalid null pointer received as argument."""
INVALID_CONFIGURATION = Status(0x0023)
"""Invalid configuration provided."""
INVALID_MODE = Status(0x0024)
"""Invalid mode."""
INVALID_HANDLE = Status(0x0025)
"""Invalid handle."""
INVALID_TYPE = Status(0x0026)
"""Invalid type for operation."""
INVALID_INDEX = Status(0x0027)
"""Invalid index."""
INVALID_RANGE = Status(0x0028)
"""Invalid range."""
INVALID_KEY = Status(0x0029)
"""Invalid key."""
INVALID_CREDENTIALS = Status(0x002A)
"""Invalid credentials."""
INVALID_COUNT = Status(0x002B)
"""Invalid count."""
INVALID_SIGNATURE = Status(0x002C)
"""Invalid signature / verification failed."""
NOT_FOUND = Status(0x002D)
"""Item could not be found."""
ALREADY_EXISTS = Status(0x002E)
"""Item already exists."""

# IO/Communication Errors
IO = Status(0x002F)
"""Generic I/O failure."""
IO_TIMEOUT = Status(0x0030)
"""I/O failure due to timeout."""
TRANSMIT = Status(0x0031)
"""Generic transmission error."""
TRANSMIT_UNDERFLOW = Status(0x0032)
"""Transmit underflowed."""
TRANSMIT_INCOMPLETE = Status(0x0033)
"""Transmit is incomplete."""
TRANSMIT_BUSY = Status(0x0034)
"""Transmit is busy."""
RECEIVE = Status(0x0035)
"""Generic reception error."""
OBJECT_READ = Status(0x0036)
"""Failed to read on/via given object."""
OBJECT_WRITE = Status(0x0037)
"""Failed to write on/via given object."""
MESSAGE_TOO_LONG = Status(0x0038)
"""Message is too long."""

# EEPROM/Flash Errors
EEPROM_MFG_VERSION_MISMATCH = Status(0x0039)
"""EEPROM MFG version mismatch."""
EEPROM_STACK_VERSION_MISMATCH = Status(0x003A)
"""EEPROM Stack version mismatch."""
FLASH_WRITE_INHIBITED = Status(0x003B)
"""Flash write is inhibited."""
FLASH_VERIFY_FAILED = Status(0x003C)
"""Flash verification failed."""
FLASH_PROGRAM_FAILED = Status(0x003D)
"""Flash programming failed."""
FLASH_ERASE_FAILED = Status(0x003E)
"""Flash erase failed."""

# MAC Errors
MAC_NO_DATA = Status(0x003F)
"""MAC no data."""
MAC_NO_ACK_RECEIVED = Status(0x0040)
"""MAC no ACK received."""
MAC_INDIRECT_TIMEOUT = Status(0x0041)
"""MAC indirect timeout."""
MAC_UNKNOWN_HEADER_TYPE = Status(0x0042)
"""MAC unknown header type."""
MAC_ACK_HEADER_TYPE = Status(0x0043)
"""MAC ACK unknown header type."""
MAC_COMMAND_TRANSMIT_FAILURE = Status(0x0044)
"""MAC command transmit failure."""

# CLI_STORAGE Errors
CLI_STORAGE_NVM_OPEN_ERROR = Status(0x0045)
"""Error in open NVM"""

# Security status codes
SECURITY_IMAGE_CHECKSUM_ERROR = Status(0x0046)
"""Image checksum is not valid."""
SECURITY_DECRYPT_ERROR = Status(0x0047)
"""Decryption failed"""

# Command status codes
COMMAND_IS_INVALID = Status(0x0048)
"""Command was not recognized"""
COMMAND_TOO_LONG = Status(0x0049)
"""Command or parameter maximum length exceeded"""
COMMAND_INCOMPLETE = Status(0x004A)
"""Data received does not form a complete command"""

# Misc Errors
BUS_ERROR = Status(0x004B)
"""Bus error, e.g. invalid DMA address"""

# Unified MAC Errors
CCA_FAILURE = Status(0x004C)
"""CCA failure."""

# Scan errors
MAC_SCANNING = Status(0x004D)
"""MAC scanning."""
MAC_INCORRECT_SCAN_TYPE = Status(0x004E)
"""MAC incorrect scan type."""
INVALID_CHANNEL_MASK = Status(0x004F)
"""Invalid channel mask."""
BAD_SCAN_DURATION = Status(0x0050)
"""Bad scan duration."""

# MAC transmit related status
MAC_TRANSMIT_QUEUE_FULL = Status(0x0053)
"""The MAC transmit queue is full"""
TRANSMIT_SCHEDULER_FAIL = Status(0x0054)
"""The transmit attempt failed because the radio scheduler could not find a slot to transmit this packet in or a higher priority event interrupted it"""
TRANSMIT_INVALID_CHANNEL = Status(0x0055)
"""An unsupported channel setting was specified"""
TRANSMIT_INVALID_POWER = Status(0x0056)
"""An unsupported power setting was specified"""
TRANSMIT_ACK_RECEIVED = Status(0x0057)
"""The expected ACK was received after the last transmission"""
TRANSMIT_BLOCKED = Status(0x0058)
"""The transmit attempt was blocked from going over the air. Typically this is due to the Radio Hold Off (RHO) or Coexistence plugins as they can prevent transmits based on external signals."""

# NVM3 specific errors
NVM3_ALIGNMENT_INVALID = Status(0x0059)
"""The initialization was aborted as the NVM3 instance is not aligned properly in memory"""
NVM3_SIZE_TOO_SMALL = Status(0x005A)
"""The initialization was aborted as the size of the NVM3 instance is too small"""
NVM3_PAGE_SIZE_NOT_SUPPORTED = Status(0x005B)
"""The initialization was aborted as the NVM3 page size is not supported"""
NVM3_TOKEN_INIT_FAILED = Status(0x005C)
"""The application that there was an error initializing some of the tokens"""
NVM3_OPENED_WITH_OTHER_PARAMETERS = Status(0x005D)
"""The initialization was aborted as the NVM3 instance was already opened with other parameters"""
NVM3_NO_VALID_PAGES = Status(0x005E)
"""Initialization aborted, no valid page found"""
NVM3_OBJECT_SIZE_NOT_SUPPORTED = Status(0x005F)
"""The object size is not supported"""
NVM3_OBJECT_IS_NOT_DATA = Status(0x0060)
"""Trying to access a data object which is currently a counter object"""
NVM3_OBJECT_IS_NOT_A_COUNTER = Status(0x0061)
"""Trying to access a counter object which is currently a data object"""
NVM3_WRITE_DATA_SIZE = Status(0x0062)
"""The object is too large"""
NVM3_READ_DATA_SIZE = Status(0x0063)
"""Trying to read with a length different from actual object size"""
NVM3_INIT_WITH_FULL_NVM = Status(0x0064)
"""The module was opened with a full NVM"""
NVM3_RESIZE_PARAMETER = Status(0x0065)
"""Illegal parameter"""
NVM3_RESIZE_NOT_ENOUGH_SPACE = Status(0x0066)
"""Not enough NVM to complete resize"""
NVM3_ERASE_COUNT_ERROR = Status(0x0067)
"""Erase counts are not valid"""
NVM3_NVM_ACCESS = Status(0x0068)
"""A NVM function call was failing"""
NVM3_CRYPTO_INIT_FAILED = Status(0x0069)
"""Crypto initialization failed"""
NVM3_ENCRYPTION_KEY_ERROR = Status(0x006A)
"""Error in obtaining encryption key"""
NVM3_RANDOM_NUM_GENERATION_FAILED = Status(0x006B)
"""Error in obtaining random number"""
NVM3_ENCRYPTION_FAILED = Status(0x006C)
"""Encryption failed"""
NVM3_WRITE_TO_NOT_ERASED = Status(0x006D)
"""Write to memory that is not erased"""
NVM3_INVALID_ADDR = Status(0x006E)
"""Invalid NVM address"""
NVM3_KEY_MISMATCH = Status(0x006F)
"""Key validation failure"""
NVM3_SIZE_ERROR = Status(0x0070)
"""Size mismatch error"""
NVM3_EMULATOR = Status(0x0071)
"""Emulator error"""
NVM3_SECURITY_INIT_FAILED = Status(0x0072)
"""Security init failed"""
NVM3_GET_REGION_LOCATION_FAILED = Status(0x0073)
"""Get data region location failed"""

# Bluetooth status codes
BT_OUT_OF_BONDS = Status(0x0402)
"""Bonding procedure can't be started because device has no space left for bond."""
BT_UNSPECIFIED = Status(0x0403)
"""Unspecified error"""
BT_HARDWARE = Status(0x0404)
"""Hardware failure"""
BT_NO_BONDING = Status(0x0406)
"""The bonding does not exist."""
BT_CRYPTO = Status(0x0407)
"""Error using crypto functions"""
BT_DATA_CORRUPTED = Status(0x0408)
"""Data was corrupted."""
BT_INVALID_SYNC_HANDLE = Status(0x040A)
"""Invalid periodic advertising sync handle"""
BT_INVALID_MODULE_ACTION = Status(0x040B)
"""Bluetooth cannot be used on this hardware"""
BT_RADIO = Status(0x040C)
"""Error received from radio"""
BT_L2CAP_REMOTE_DISCONNECTED = Status(0x040D)
"""Returned when remote disconnects the connection-oriented channel by sending disconnection request."""
BT_L2CAP_LOCAL_DISCONNECTED = Status(0x040E)
"""Returned when local host disconnect the connection-oriented channel by sending disconnection request."""
BT_L2CAP_CID_NOT_EXIST = Status(0x040F)
"""Returned when local host did not find a connection-oriented channel with given destination CID."""
BT_L2CAP_LE_DISCONNECTED = Status(0x0410)
"""Returned when connection-oriented channel disconnected due to LE connection is dropped."""
BT_L2CAP_FLOW_CONTROL_VIOLATED = Status(0x0412)
"""Returned when connection-oriented channel disconnected due to remote end send data even without credit."""
BT_L2CAP_FLOW_CONTROL_CREDIT_OVERFLOWED = Status(0x0413)
"""Returned when connection-oriented channel disconnected due to remote end send flow control credits exceed 65535."""
BT_L2CAP_NO_FLOW_CONTROL_CREDIT = Status(0x0414)
"""Returned when connection-oriented channel has run out of flow control credit and local application still trying to send data."""
BT_L2CAP_CONNECTION_REQUEST_TIMEOUT = Status(0x0415)
"""Returned when connection-oriented channel has not received connection response message within maximum timeout."""
BT_L2CAP_INVALID_CID = Status(0x0416)
"""Returned when local host received a connection-oriented channel connection response with an invalid destination CID."""
BT_L2CAP_WRONG_STATE = Status(0x0417)
"""Returned when local host application tries to send a command which is not suitable for L2CAP channel's current state."""
BT_PS_STORE_FULL = Status(0x041B)
"""Flash reserved for PS store is full"""
BT_PS_KEY_NOT_FOUND = Status(0x041C)
"""PS key not found"""
BT_APPLICATION_MISMATCHED_OR_INSUFFICIENT_SECURITY = Status(0x041D)
"""Mismatched or insufficient security level"""
BT_APPLICATION_ENCRYPTION_DECRYPTION_ERROR = Status(0x041E)
"""Encryption/decryption operation failed."""

# Bluetooth controller status codes
BT_CTRL_UNKNOWN_CONNECTION_IDENTIFIER = Status(0x1002)
"""Connection does not exist, or connection open request was cancelled."""
BT_CTRL_AUTHENTICATION_FAILURE = Status(0x1005)
"""Pairing or authentication failed due to incorrect results in the pairing or authentication procedure. This could be due to an incorrect PIN or Link Key"""
BT_CTRL_PIN_OR_KEY_MISSING = Status(0x1006)
"""Pairing failed because of missing PIN, or authentication failed because of missing Key"""
BT_CTRL_MEMORY_CAPACITY_EXCEEDED = Status(0x1007)
"""Controller is out of memory."""
BT_CTRL_CONNECTION_TIMEOUT = Status(0x1008)
"""Link supervision timeout has expired."""
BT_CTRL_CONNECTION_LIMIT_EXCEEDED = Status(0x1009)
"""Controller is at limit of connections it can support."""
BT_CTRL_SYNCHRONOUS_CONNECTION_LIMIT_EXCEEDED = Status(0x100A)
"""The Synchronous Connection Limit to a Device Exceeded error code indicates that the Controller has reached the limit to the number of synchronous connections that can be achieved to a device."""
BT_CTRL_ACL_CONNECTION_ALREADY_EXISTS = Status(0x100B)
"""The ACL Connection Already Exists error code indicates that an attempt to create a new ACL Connection to a device when there is already a connection to this device."""
BT_CTRL_COMMAND_DISALLOWED = Status(0x100C)
"""Command requested cannot be executed because the Controller is in a state where it cannot process this command at this time."""
BT_CTRL_CONNECTION_REJECTED_DUE_TO_LIMITED_RESOURCES = Status(0x100D)
"""The Connection Rejected Due To Limited Resources error code indicates that an incoming connection was rejected due to limited resources."""
BT_CTRL_CONNECTION_REJECTED_DUE_TO_SECURITY_REASONS = Status(0x100E)
"""The Connection Rejected Due To Security Reasons error code indicates that a connection was rejected due to security requirements not being fulfilled, like authentication or pairing."""
BT_CTRL_CONNECTION_REJECTED_DUE_TO_UNACCEPTABLE_BD_ADDR = Status(0x100F)
"""The Connection was rejected because this device does not accept the BD_ADDR. This may be because the device will only accept connections from specific BD_ADDRs."""
BT_CTRL_CONNECTION_ACCEPT_TIMEOUT_EXCEEDED = Status(0x1010)
"""The Connection Accept Timeout has been exceeded for this connection attempt."""
BT_CTRL_UNSUPPORTED_FEATURE_OR_PARAMETER_VALUE = Status(0x1011)
"""A feature or parameter value in the HCI command is not supported."""
BT_CTRL_INVALID_COMMAND_PARAMETERS = Status(0x1012)
"""Command contained invalid parameters."""
BT_CTRL_REMOTE_USER_TERMINATED = Status(0x1013)
"""User on the remote device terminated the connection."""
BT_CTRL_REMOTE_DEVICE_TERMINATED_CONNECTION_DUE_TO_LOW_RESOURCES = Status(0x1014)
"""The remote device terminated the connection because of low resources"""
BT_CTRL_REMOTE_POWERING_OFF = Status(0x1015)
"""Remote Device Terminated Connection due to Power Off"""
BT_CTRL_CONNECTION_TERMINATED_BY_LOCAL_HOST = Status(0x1016)
"""Local device terminated the connection."""
BT_CTRL_REPEATED_ATTEMPTS = Status(0x1017)
"""The Controller is disallowing an authentication or pairing procedure because too little time has elapsed since the last authentication or pairing attempt failed."""
BT_CTRL_PAIRING_NOT_ALLOWED = Status(0x1018)
"""The device does not allow pairing. This can be for example, when a device only allows pairing during a certain time window after some user input allows pairing"""
BT_CTRL_UNSUPPORTED_REMOTE_FEATURE = Status(0x101A)
"""The remote device does not support the feature associated with the issued command."""
BT_CTRL_INVALID_LL_PARAMETERS = Status(0x101E)
"""Indicates that some LMP PDU / LL Control PDU parameters were invalid"""
BT_CTRL_UNSPECIFIED_ERROR = Status(0x101F)
"""No other error code specified is appropriate to use."""
BT_CTRL_LL_RESPONSE_TIMEOUT = Status(0x1022)
"""Connection terminated due to link-layer procedure timeout."""
BT_CTRL_LL_PROCEDURE_COLLISION = Status(0x1023)
"""LL procedure has collided with the same transaction or procedure that is already in progress."""
BT_CTRL_ENCRYPTION_MODE_NOT_ACCEPTABLE = Status(0x1025)
"""The requested encryption mode is not acceptable at this time."""
BT_CTRL_LINK_KEY_CANNOT_BE_CHANGED = Status(0x1026)
"""Link key cannot be changed because a fixed unit key is being used."""
BT_CTRL_INSTANT_PASSED = Status(0x1028)
"""LMP PDU or LL PDU that includes an instant cannot be performed because the instant when this would have occurred has passed."""
BT_CTRL_PAIRING_WITH_UNIT_KEY_NOT_SUPPORTED = Status(0x1029)
"""It was not possible to pair as a unit key was requested and it is not supported."""
BT_CTRL_DIFFERENT_TRANSACTION_COLLISION = Status(0x102A)
"""LMP transaction was started that collides with an ongoing transaction."""
BT_CTRL_CHANNEL_ASSESSMENT_NOT_SUPPORTED = Status(0x102E)
"""The Controller cannot perform channel assessment because it is not supported."""
BT_CTRL_INSUFFICIENT_SECURITY = Status(0x102F)
"""The HCI command or LMP PDU sent is only possible on an encrypted link."""
BT_CTRL_PARAMETER_OUT_OF_MANDATORY_RANGE = Status(0x1030)
"""A parameter value requested is outside the mandatory range of parameters for the given HCI command or LMP PDU."""
BT_CTRL_SIMPLE_PAIRING_NOT_SUPPORTED_BY_HOST = Status(0x1037)
"""The IO capabilities request or response was rejected because the sending Host does not support Secure Simple Pairing even though the receiving Link Manager does."""
BT_CTRL_HOST_BUSY_PAIRING = Status(0x1038)
"""The Host is busy with another pairing operation and unable to support the requested pairing. The receiving device should retry pairing again later."""
BT_CTRL_CONNECTION_REJECTED_DUE_TO_NO_SUITABLE_CHANNEL_FOUND = Status(0x1039)
"""The Controller could not calculate an appropriate value for the Channel selection operation."""
BT_CTRL_CONTROLLER_BUSY = Status(0x103A)
"""Operation was rejected because the controller is busy and unable to process the request."""
BT_CTRL_UNACCEPTABLE_CONNECTION_INTERVAL = Status(0x103B)
"""Remote device terminated the connection because of an unacceptable connection interval."""
BT_CTRL_ADVERTISING_TIMEOUT = Status(0x103C)
"""Advertising for a fixed duration completed or, for directed advertising, that advertising completed without a connection being created."""
BT_CTRL_CONNECTION_TERMINATED_DUE_TO_MIC_FAILURE = Status(0x103D)
"""Connection was terminated because the Message Integrity Check (MIC) failed on a received packet."""
BT_CTRL_CONNECTION_FAILED_TO_BE_ESTABLISHED = Status(0x103E)
"""LL initiated a connection but the connection has failed to be established. Controller did not receive any packets from remote end."""
BT_CTRL_MAC_CONNECTION_FAILED = Status(0x103F)
"""The MAC of the 802.11 AMP was requested to connect to a peer, but the connection failed."""
BT_CTRL_COARSE_CLOCK_ADJUSTMENT_REJECTED_BUT_WILL_TRY_TO_ADJUST_USING_CLOCK_DRAGGING = Status(0x1040)
"""The master, at this time, is unable to make a coarse adjustment to the piconet clock, using the supplied parameters. Instead the master will attempt to move the clock using clock dragging."""
BT_CTRL_UNKNOWN_ADVERTISING_IDENTIFIER = Status(0x1042)
"""A command was sent from the Host that should identify an Advertising or Sync handle, but the Advertising or Sync handle does not exist."""
BT_CTRL_LIMIT_REACHED = Status(0x1043)
"""Number of operations requested has been reached and has indicated the completion of the activity (e.g., advertising or scanning)."""
BT_CTRL_OPERATION_CANCELLED_BY_HOST = Status(0x1044)
"""A request to the Controller issued by the Host and still pending was successfully canceled."""
BT_CTRL_PACKET_TOO_LONG = Status(0x1045)
"""An attempt was made to send or receive a packet that exceeds the maximum allowed packet length."""
BT_CTRL_TOO_LATE = Status(0x1046)
"""Information was provided too late to the controller."""
BT_CTRL_TOO_EARLY = Status(0x1047)
"""Information was provided too early to the controller."""
BT_CTRL_INSUFFICIENT_CHANNELS = Status(0x1048)
"""Indicates that the result of the requested operation would yield too few physical channels."""

# Bluetooth attribute status codes
BT_ATT_INVALID_HANDLE = Status(0x1101)
"""The attribute handle given was not valid on this server"""
BT_ATT_READ_NOT_PERMITTED = Status(0x1102)
"""The attribute cannot be read"""
BT_ATT_WRITE_NOT_PERMITTED = Status(0x1103)
"""The attribute cannot be written"""
BT_ATT_INVALID_PDU = Status(0x1104)
"""The attribute PDU was invalid"""
BT_ATT_INSUFFICIENT_AUTHENTICATION = Status(0x1105)
"""The attribute requires authentication before it can be read or written."""
BT_ATT_REQUEST_NOT_SUPPORTED = Status(0x1106)
"""Attribute Server does not support the request received from the client."""
BT_ATT_INVALID_OFFSET = Status(0x1107)
"""Offset specified was past the end of the attribute"""
BT_ATT_INSUFFICIENT_AUTHORIZATION = Status(0x1108)
"""The attribute requires authorization before it can be read or written."""
BT_ATT_PREPARE_QUEUE_FULL = Status(0x1109)
"""Too many prepare writes have been queued"""
BT_ATT_ATT_NOT_FOUND = Status(0x110A)
"""No attribute found within the given attribute handle range."""
BT_ATT_ATT_NOT_LONG = Status(0x110B)
"""The attribute cannot be read or written using the Read Blob Request"""
BT_ATT_INSUFFICIENT_ENC_KEY_SIZE = Status(0x110C)
"""The Encryption Key Size used for encrypting this link is insufficient."""
BT_ATT_INVALID_ATT_LENGTH = Status(0x110D)
"""The attribute value length is invalid for the operation"""
BT_ATT_UNLIKELY_ERROR = Status(0x110E)
"""The attribute request that was requested has encountered an error that was unlikely, and therefore could not be completed as requested."""
BT_ATT_INSUFFICIENT_ENCRYPTION = Status(0x110F)
"""The attribute requires encryption before it can be read or written."""
BT_ATT_UNSUPPORTED_GROUP_TYPE = Status(0x1110)
"""The attribute type is not a supported grouping attribute as defined by a higher layer specification."""
BT_ATT_INSUFFICIENT_RESOURCES = Status(0x1111)
"""Insufficient Resources to complete the request"""
BT_ATT_OUT_OF_SYNC = Status(0x1112)
"""The server requests the client to rediscover the database."""
BT_ATT_VALUE_NOT_ALLOWED = Status(0x1113)
"""The attribute parameter value was not allowed."""
BT_ATT_APPLICATION = Status(0x1180)
"""When this is returned in a BGAPI response, the application tried to read or write the value of a user attribute from the GATT database."""
BT_ATT_WRITE_REQUEST_REJECTED = Status(0x11FC)
"""The requested write operation cannot be fulfilled for reasons other than permissions."""
BT_ATT_CLIENT_CHARACTERISTIC_CONFIGURATION_DESCRIPTOR_IMPROPERLY_CONFIGURED = Status(0x11FD)
"""The Client Characteristic Configuration descriptor is not configured according to the requirements of the profile or service."""
BT_ATT_PROCEDURE_ALREADY_IN_PROGRESS = Status(0x11FE)
"""The profile or service request cannot be serviced because an operation that has been previously triggered is still in progress."""
BT_ATT_OUT_OF_RANGE = Status(0x11FF)
"""The attribute value is out of range as defined by a profile or service specification."""

# Bluetooth Security Manager Protocol status codes
BT_SMP_PASSKEY_ENTRY_FAILED = Status(0x1201)
"""The user input of passkey failed, for example, the user cancelled the operation"""
BT_SMP_OOB_NOT_AVAILABLE = Status(0x1202)
"""Out of Band data is not available for authentication"""
BT_SMP_AUTHENTICATION_REQUIREMENTS = Status(0x1203)
"""The pairing procedure cannot be performed as authentication requirements cannot be met due to IO capabilities of one or both devices"""
BT_SMP_CONFIRM_VALUE_FAILED = Status(0x1204)
"""The confirm value does not match the calculated compare value"""
BT_SMP_PAIRING_NOT_SUPPORTED = Status(0x1205)
"""Pairing is not supported by the device"""
BT_SMP_ENCRYPTION_KEY_SIZE = Status(0x1206)
"""The resultant encryption key size is insufficient for the security requirements of this device"""
BT_SMP_COMMAND_NOT_SUPPORTED = Status(0x1207)
"""The SMP command received is not supported on this device"""
BT_SMP_UNSPECIFIED_REASON = Status(0x1208)
"""Pairing failed due to an unspecified reason"""
BT_SMP_REPEATED_ATTEMPTS = Status(0x1209)
"""Pairing or authentication procedure is disallowed because too little time has elapsed since last pairing request or security request"""
BT_SMP_INVALID_PARAMETERS = Status(0x120A)
"""The Invalid Parameters error code indicates: the command length is invalid or a parameter is outside of the specified range."""
BT_SMP_DHKEY_CHECK_FAILED = Status(0x120B)
"""Indicates to the remote device that the DHKey Check value received doesn't match the one calculated by the local device."""
BT_SMP_NUMERIC_COMPARISON_FAILED = Status(0x120C)
"""Indicates that the confirm values in the numeric comparison protocol do not match."""
BT_SMP_BREDR_PAIRING_IN_PROGRESS = Status(0x120D)
"""Indicates that the pairing over the LE transport failed due to a Pairing Request sent over the BR/EDR transport in process."""
BT_SMP_CROSS_TRANSPORT_KEY_DERIVATION_GENERATION_NOT_ALLOWED = Status(0x120E)
"""Indicates that the BR/EDR Link Key generated on the BR/EDR transport cannot be used to derive and distribute keys for the LE transport."""
BT_SMP_KEY_REJECTED = Status(0x120F)
"""Indicates that the device chose not to accept a distributed key."""

# Bluetooth Mesh status codes
BT_MESH_ALREADY_EXISTS = Status(0x0501)
"""Returned when trying to add a key or some other unique resource with an ID which already exists"""
BT_MESH_DOES_NOT_EXIST = Status(0x0502)
"""Returned when trying to manipulate a key or some other resource with an ID which does not exist"""
BT_MESH_LIMIT_REACHED = Status(0x0503)
"""Returned when an operation cannot be executed because a pre-configured limit for keys, key bindings, elements, models, virtual addresses, provisioned devices, or provisioning sessions is reached"""
BT_MESH_INVALID_ADDRESS = Status(0x0504)
"""Returned when trying to use a reserved address or add a "pre-provisioned" device using an address already used by some other device"""
BT_MESH_MALFORMED_DATA = Status(0x0505)
"""In a BGAPI response, the user supplied malformed data; in a BGAPI event, the remote end responded with malformed or unrecognized data"""
BT_MESH_ALREADY_INITIALIZED = Status(0x0506)
"""An attempt was made to initialize a subsystem that was already initialized."""
BT_MESH_NOT_INITIALIZED = Status(0x0507)
"""An attempt was made to use a subsystem that wasn't initialized yet. Call the subsystem's init function first."""
BT_MESH_NO_FRIEND_OFFER = Status(0x0508)
"""Returned when trying to establish a friendship as a Low Power Node, but no acceptable friend offer message was received."""
BT_MESH_PROV_LINK_CLOSED = Status(0x0509)
"""Provisioning link was unexpectedly closed before provisioning was complete."""
BT_MESH_PROV_INVALID_PDU = Status(0x050A)
"""An unrecognized provisioning PDU was received."""
BT_MESH_PROV_INVALID_PDU_FORMAT = Status(0x050B)
"""A provisioning PDU with wrong length or containing field values that are out of bounds was received."""
BT_MESH_PROV_UNEXPECTED_PDU = Status(0x050C)
"""An unexpected (out of sequence) provisioning PDU was received."""
BT_MESH_PROV_CONFIRMATION_FAILED = Status(0x050D)
"""The computed confirmation value did not match the expected value."""
BT_MESH_PROV_OUT_OF_RESOURCES = Status(0x050E)
"""Provisioning could not be continued due to insufficient resources."""
BT_MESH_PROV_DECRYPTION_FAILED = Status(0x050F)
"""The provisioning data block could not be decrypted."""
BT_MESH_PROV_UNEXPECTED_ERROR = Status(0x0510)
"""An unexpected error happened during provisioning."""
BT_MESH_PROV_CANNOT_ASSIGN_ADDR = Status(0x0511)
"""Device could not assign unicast addresses to all of its elements."""
BT_MESH_ADDRESS_TEMPORARILY_UNAVAILABLE = Status(0x0512)
"""Returned when trying to reuse an address of a previously deleted device before an IV Index Update has been executed."""
BT_MESH_ADDRESS_ALREADY_USED = Status(0x0513)
"""Returned when trying to assign an address that is used by one of the devices in the Device Database, or by the Provisioner itself."""
BT_MESH_PUBLISH_NOT_CONFIGURED = Status(0x0514)
"""Application key or publish address are not set"""
BT_MESH_APP_KEY_NOT_BOUND = Status(0x0515)
"""Application key is not bound to a model"""

# Bluetooth Mesh foundation status codes
BT_MESH_FOUNDATION_INVALID_ADDRESS = Status(0x1301)
"""Returned when address in request was not valid"""
BT_MESH_FOUNDATION_INVALID_MODEL = Status(0x1302)
"""Returned when model identified is not found for a given element"""
BT_MESH_FOUNDATION_INVALID_APP_KEY = Status(0x1303)
"""Returned when the key identified by AppKeyIndex is not stored in the node"""
BT_MESH_FOUNDATION_INVALID_NET_KEY = Status(0x1304)
"""Returned when the key identified by NetKeyIndex is not stored in the node"""
BT_MESH_FOUNDATION_INSUFFICIENT_RESOURCES = Status(0x1305)
"""Returned when The node cannot serve the request due to insufficient resources"""
BT_MESH_FOUNDATION_KEY_INDEX_EXISTS = Status(0x1306)
"""Returned when the key identified is already stored in the node and the new NetKey value is different"""
BT_MESH_FOUNDATION_INVALID_PUBLISH_PARAMS = Status(0x1307)
"""Returned when the model does not support the publish mechanism"""
BT_MESH_FOUNDATION_NOT_SUBSCRIBE_MODEL = Status(0x1308)
"""Returned when  the model does not support the subscribe mechanism"""
BT_MESH_FOUNDATION_STORAGE_FAILURE = Status(0x1309)
"""Returned when storing of the requested parameters failed"""
BT_MESH_FOUNDATION_NOT_SUPPORTED = Status(0x130A)
"""Returned when requested setting is not supported"""
BT_MESH_FOUNDATION_CANNOT_UPDATE = Status(0x130B)
"""Returned when the requested update operation cannot be performed due to general constraints"""
BT_MESH_FOUNDATION_CANNOT_REMOVE = Status(0x130C)
"""Returned when the requested delete operation cannot be performed due to general constraints"""
BT_MESH_FOUNDATION_CANNOT_BIND = Status(0x130D)
"""Returned when the requested bind operation cannot be performed due to general constraints"""
BT_MESH_FOUNDATION_TEMPORARILY_UNABLE = Status(0x130E)
"""Returned when The node cannot start advertising with Node Identity or Proxy since the maximum number of parallel advertising is reached"""
BT_MESH_FOUNDATION_CANNOT_SET = Status(0x130F)
"""Returned when the requested state cannot be set"""
BT_MESH_FOUNDATION_UNSPECIFIED = Status(0x1310)
"""Returned when an unspecified error took place"""
BT_MESH_FOUNDATION_INVALID_BINDING = Status(0x1311)
"""Returned when the NetKeyIndex and AppKeyIndex combination is not valid for a Config AppKey Update"""

# Wi-Fi Errors
WIFI_INVALID_KEY = Status(0x0B01)
"""Invalid firmware keyset"""
WIFI_FIRMWARE_DOWNLOAD_TIMEOUT = Status(0x0B02)
"""The firmware download took too long"""
WIFI_UNSUPPORTED_MESSAGE_ID = Status(0x0B03)
"""Unknown request ID or wrong interface ID used"""
WIFI_WARNING = Status(0x0B04)
"""The request is successful but some parameters have been ignored"""
WIFI_NO_PACKET_TO_RECEIVE = Status(0x0B05)
"""No Packets waiting to be received"""
WIFI_SLEEP_GRANTED = Status(0x0B08)
"""The sleep mode is granted"""
WIFI_SLEEP_NOT_GRANTED = Status(0x0B09)
"""The WFx does not go back to sleep"""
WIFI_SECURE_LINK_MAC_KEY_ERROR = Status(0x0B10)
"""The SecureLink MAC key was not found"""
WIFI_SECURE_LINK_MAC_KEY_ALREADY_BURNED = Status(0x0B11)
"""The SecureLink MAC key is already installed in OTP"""
WIFI_SECURE_LINK_RAM_MODE_NOT_ALLOWED = Status(0x0B12)
"""The SecureLink MAC key cannot be installed in RAM"""
WIFI_SECURE_LINK_FAILED_UNKNOWN_MODE = Status(0x0B13)
"""The SecureLink MAC key installation failed"""
WIFI_SECURE_LINK_EXCHANGE_FAILED = Status(0x0B14)
"""SecureLink key (re)negotiation failed"""
WIFI_WRONG_STATE = Status(0x0B18)
"""The device is in an inappropriate state to perform the request"""
WIFI_CHANNEL_NOT_ALLOWED = Status(0x0B19)
"""The request failed due to regulatory limitations"""
WIFI_NO_MATCHING_AP = Status(0x0B1A)
"""The connection request failed because no suitable AP was found"""
WIFI_CONNECTION_ABORTED = Status(0x0B1B)
"""The connection request was aborted by host"""
WIFI_CONNECTION_TIMEOUT = Status(0x0B1C)
"""The connection request failed because of a timeout"""
WIFI_CONNECTION_REJECTED_BY_AP = Status(0x0B1D)
"""The connection request failed because the AP rejected the device"""
WIFI_CONNECTION_AUTH_FAILURE = Status(0x0B1E)
"""The connection request failed because the WPA handshake did not complete successfully"""
WIFI_RETRY_EXCEEDED = Status(0x0B1F)
"""The request failed because the retry limit was exceeded"""
WIFI_TX_LIFETIME_EXCEEDED = Status(0x0B20)
"""The request failed because the MSDU life time was exceeded"""

# MVP Driver and MVP Math status codes
COMPUTE_DRIVER_FAULT = Status(0x1501)
"""Critical fault"""
COMPUTE_DRIVER_ALU_NAN = Status(0x1502)
"""ALU operation output NaN"""
COMPUTE_DRIVER_ALU_OVERFLOW = Status(0x1503)
"""ALU numeric overflow"""
COMPUTE_DRIVER_ALU_UNDERFLOW = Status(0x1504)
"""ALU numeric underflow"""
COMPUTE_DRIVER_STORE_CONVERSION_OVERFLOW = Status(0x1505)
"""Overflow during array store"""
COMPUTE_DRIVER_STORE_CONVERSION_UNDERFLOW = Status(0x1506)
"""Underflow during array store conversion"""
COMPUTE_DRIVER_STORE_CONVERSION_INFINITY = Status(0x1507)
"""Infinity encountered during array store conversion"""
COMPUTE_DRIVER_STORE_CONVERSION_NAN = Status(0x1508)
"""NaN encountered during array store conversion"""
COMPUTE_MATH_NAN = Status(0x1512)
"""MATH NaN encountered"""
COMPUTE_MATH_INFINITY = Status(0x1513)
"""MATH Infinity encountered"""
COMPUTE_MATH_OVERFLOW = Status(0x1514)
"""MATH numeric overflow"""
COMPUTE_MATH_UNDERFLOW = Status(0x1515)
"""MATH numeric underflow"""

# Zigbee status codes
ZIGBEE_PACKET_HANDOFF_DROPPED = Status(0x0C01)
"""Packet is dropped by packet-handoff callbacks"""
ZIGBEE_DELIVERY_FAILED = Status(0x0C02)
"""The APS layer attempted to send or deliver a message and failed"""
ZIGBEE_MAX_MESSAGE_LIMIT_REACHED = Status(0x0C03)
"""The maximum number of in-flight messages ::EMBER_APS_UNICAST_MESSAGE_COUNT has been reached"""
ZIGBEE_BINDING_IS_ACTIVE = Status(0x0C04)
"""The application is trying to delete or overwrite a binding that is in use"""
ZIGBEE_ADDRESS_TABLE_ENTRY_IS_ACTIVE = Status(0x0C05)
"""The application is trying to overwrite an address table entry that is in use"""
ZIGBEE_MOVE_FAILED = Status(0x0C06)
"""After moving, a mobile node's attempt to re-establish contact with the network failed"""
ZIGBEE_NODE_ID_CHANGED = Status(0x0C07)
"""The local node ID has changed. The application can get the new node ID by calling ::sl_zigbee_get_node_id()"""
ZIGBEE_INVALID_SECURITY_LEVEL = Status(0x0C08)
"""The chosen security level is not supported by the stack"""
ZIGBEE_IEEE_ADDRESS_DISCOVERY_IN_PROGRESS = Status(0x0C09)
"""An error occurred when trying to encrypt at the APS Level"""
ZIGBEE_APS_ENCRYPTION_ERROR = Status(0x0C0A)
"""An error occurred when trying to encrypt at the APS Level"""
ZIGBEE_SECURITY_STATE_NOT_SET = Status(0x0C0B)
"""There was an attempt to form or join a network with security without calling ::sl_zigbee_set_initial_security_state() first"""
ZIGBEE_TOO_SOON_FOR_SWITCH_KEY = Status(0x0C0C)
"""There was an attempt to broadcast a key switch too quickly after broadcasting the next network key. The Trust Center must wait at least a period equal to the broadcast timeout so that all routers have a chance to receive the broadcast of the new network key"""
ZIGBEE_SIGNATURE_VERIFY_FAILURE = Status(0x0C0D)
"""The received signature corresponding to the message that was passed to the CBKE Library failed verification and is not valid"""
ZIGBEE_KEY_NOT_AUTHORIZED = Status(0x0C0E)
"""The message could not be sent because the link key corresponding to the destination is not authorized for use in APS data messages"""
ZIGBEE_BINDING_HAS_CHANGED = Status(0x0C0F)
"""The application tried to use a binding that has been remotely modified and the change has not yet been reported to the application"""
ZIGBEE_TRUST_CENTER_SWAP_EUI_HAS_CHANGED = Status(0x0C10)
"""The EUI of the Trust center has changed due to a successful rejoin after TC Swapout"""
ZIGBEE_TRUST_CENTER_SWAP_EUI_HAS_NOT_CHANGED = Status(0x0C11)
"""A Trust Center Swapout Rejoin has occurred without the EUI of the TC changing"""
ZIGBEE_INSUFFICIENT_RANDOM_DATA = Status(0x0C12)
"""An attempt to generate random bytes failed because of insufficient random data from the radio"""
ZIGBEE_SOURCE_ROUTE_FAILURE = Status(0x0C13)
"""A Zigbee route error command frame was received indicating that a source routed message from this node failed en route"""
ZIGBEE_MANY_TO_ONE_ROUTE_FAILURE = Status(0x0C14)
"""A Zigbee route error command frame was received indicating that a message sent to this node along a many-to-one route failed en route"""
ZIGBEE_STACK_AND_HARDWARE_MISMATCH = Status(0x0C15)
"""A critical and fatal error indicating that the version of the stack trying to run does not match with the chip it's running on"""
ZIGBEE_PAN_ID_CHANGED = Status(0x0C16)
"""The local PAN ID has changed. The application can get the new PAN ID by calling ::emberGetPanId()"""
ZIGBEE_CHANNEL_CHANGED = Status(0x0C17)
"""The channel has changed."""
ZIGBEE_NETWORK_OPENED = Status(0x0C18)
"""The network has been opened for joining."""
ZIGBEE_NETWORK_CLOSED = Status(0x0C19)
"""The network has been closed for joining."""
ZIGBEE_RECEIVED_KEY_IN_THE_CLEAR = Status(0x0C1A)
"""An attempt was made to join a Secured Network using a pre-configured key, but the Trust Center sent back a Network Key in-the-clear when an encrypted Network Key was required. (::EMBER_REQUIRE_ENCRYPTED_KEY)"""
ZIGBEE_NO_NETWORK_KEY_RECEIVED = Status(0x0C1B)
"""An attempt was made to join a Secured Network, but the device did not receive a Network Key."""
ZIGBEE_NO_LINK_KEY_RECEIVED = Status(0x0C1C)
"""After a device joined a Secured Network, a Link Key was requested (::EMBER_GET_LINK_KEY_WHEN_JOINING) but no response was ever received."""
ZIGBEE_PRECONFIGURED_KEY_REQUIRED = Status(0x0C1D)
"""An attempt was made to join a Secured Network without a pre-configured key, but the Trust Center sent encrypted data using a pre-configured key."""
ZIGBEE_EZSP_ERROR = Status(0x0C1E)
"""A Zigbee EZSP error has occured. Track the origin and corresponding EzspStatus for more info."""

_doc = {
    # Generic Errors
    0x0000: "No error.",
    0x0001: "Generic error.",

    # State Errors
    0x0002: "Generic invalid state error.",
    0x0003: "Module is not ready for requested operation.",
    0x0004: "Module is busy and cannot carry out requested operation.",
    0x0005: "Operation is in progress and not yet complete (pass or fail).",
    0x0006: "Operation aborted.",
    0x0007: "Operation timed out.",
    0x0008: "Operation not allowed per permissions.",
    0x0009: "Non-blocking operation would block.",
    0x000A: "Operation/module is Idle, cannot carry requested operation.",
    0x000B: "Operation cannot be done while construct is waiting.",
    0x000C: "No task/construct waiting/pending for that action/event.",
    0x000D: "Operation cannot be done while construct is suspended.",
    0x000E: "Feature not available due to software configuration.",
    0x000F: "Feature not supported.",
    0x0010: "Initialization failed.",
    0x0011: "Module has not been initialized.",
    0x0012: "Module has already been initialized.",
    0x0013: "Object/construct has been deleted.",
    0x0014: "Illegal call from ISR.",
    0x0015: "Illegal call because network is up.",
    0x0016: "Illegal call because network is down.",
    0x0017: "Failure due to not being joined in a network.",
    0x0018: "Invalid operation as there are no beacons.",

    # Allocation/ownership Errors
    0x0019: "Generic allocation error.",
    0x001A: "No more resource available to perform the operation.",
    0x001B: "Item/list/queue is empty.",
    0x001C: "Item/list/queue is full.",
    0x001D: "Item would overflow.",
    0x001E: "Item/list/queue has been overflowed.",
    0x001F: "Generic ownership error.",
    0x0020: "Already/still owning resource.",

    # Invalid Parameters Errors
    0x0021: "Generic invalid argument or consequence of invalid argument.",
    0x0022: "Invalid null pointer received as argument.",
    0x0023: "Invalid configuration provided.",
    0x0024: "Invalid mode.",
    0x0025: "Invalid handle.",
    0x0026: "Invalid type for operation.",
    0x0027: "Invalid index.",
    0x0028: "Invalid range.",
    0x0029: "Invalid key.",
    0x002A: "Invalid credentials.",
    0x002B: "Invalid count.",
    0x002C: "Invalid signature / verification failed.",
    0x002D: "Item could not be found.",
    0x002E: "Item already exists.",

    # IO/Communication Errors
    0x002F: "Generic I/O failure.",
    0x0030: "I/O failure due to timeout.",
    0x0031: "Generic transmission error.",
    0x0032: "Transmit underflowed.",
    0x0033: "Transmit is incomplete.",
    0x0034: "Transmit is busy.",
    0x0035: "Generic reception error.",
    0x0036: "Failed to read on/via given object.",
    0x0037: "Failed to write on/via given object.",
    0x0038: "Message is too long.",

    # EEPROM/Flash Errors
    0x0039: "EEPROM MFG version mismatch.",
    0x003A: "EEPROM Stack version mismatch.",
    0x003B: "Flash write is inhibited.",
    0x003C: "Flash verification failed.",
    0x003D: "Flash programming failed.",
    0x003E: "Flash erase failed.",

    # MAC Errors
    0x003F: "MAC no data.",
    0x0040: "MAC no ACK received.",
    0x0041: "MAC indirect timeout.",
    0x0042: "MAC unknown header type.",
    0x0043: "MAC ACK unknown header type.",
    0x0044: "MAC command transmit failure.",

    # CLI_STORAGE Errors
    0x0045: "Error in open NVM",

    # Security status codes
    0x0046: "Image checksum is not valid.",
    0x0047: "Decryption failed",

    # Command status codes
    0x0048: "Command was not recognized",
    0x0049: "Command or parameter maximum length exceeded",
    0x004A: "Data received does not form a complete command",

    # Misc Errors
    0x004B: "Bus error, e.g. invalid DMA address",

    # Unified MAC Errors
    0x004C: "CCA failure.",

    # Scan errors
    0x004D: "MAC scanning.",
    0x004E: "MAC incorrect scan type.",
    0x004F: "Invalid channel mask.",
    0x0050: "Bad scan duration.",

    # MAC transmit related status
    0x0053: "The MAC transmit queue is full",
    0x0054: "The transmit attempt failed because the radio scheduler could not find a slot to transmit this packet in or a higher priority event interrupted it",
    0x0055: "An unsupported channel setting was specified",
    0x0056: "An unsupported power setting was specified",
    0x0057: "The expected ACK was received after the last transmission",
    0x0058: "The transmit attempt was blocked from going over the air. Typically this is due to the Radio Hold Off (RHO) or Coexistence plugins as they can prevent transmits based on external signals.",

    # NVM3 specific errors
    0x0059: "The initialization was aborted as the NVM3 instance is not aligned properly in memory",
    0x005A: "The initialization was aborted as the size of the NVM3 instance is too small",
    0x005B: "The initialization was aborted as the NVM3 page size is not supported",
    0x005C: "The application that there was an error initializing some of the tokens",
    0x005D: "The initialization was aborted as the NVM3 instance was already opened with other parameters",
    0x005E: "Initialization aborted, no valid page found",
    0x005F: "The object size is not supported",
    0x0060: "Trying to access a data object which is currently a counter object",
    0x0061: "Trying to access a counter object which is currently a data object",
    0x0062: "The object is too large",
    0x0063: "Trying to read with a length different from actual object size",
    0x0064: "The module was opened with a full NVM",
    0x0065: "Illegal parameter",
    0x0066: "Not enough NVM to complete resize",
    0x0067: "Erase counts are not valid",
    0x0068: "A NVM function call was failing",
    0x0069: "Crypto initialization failed",
    0x006A: "Error in obtaining encryption key",
    0x006B: "Error in obtaining random number",
    0x006C: "Encryption failed",
    0x006D: "Write to memory that is not erased",
    0x006E: "Invalid NVM address",
    0x006F: "Key validation failure",
    0x0070: "Size mismatch error",
    0x0071: "Emulator error",
    0x0072: "Security init failed",
    0x0073: "Get data region location failed",

    # Bluetooth status codes
    0x0402: "Bonding procedure can't be started because device has no space left for bond.",
    0x0403: "Unspecified error",
    0x0404: "Hardware failure",
    0x0406: "The bonding does not exist.",
    0x0407: "Error using crypto functions",
    0x0408: "Data was corrupted.",
    0x040A: "Invalid periodic advertising sync handle",
    0x040B: "Bluetooth cannot be used on this hardware",
    0x040C: "Error received from radio",
    0x040D: "Returned when remote disconnects the connection-oriented channel by sending disconnection request.",
    0x040E: "Returned when local host disconnect the connection-oriented channel by sending disconnection request.",
    0x040F: "Returned when local host did not find a connection-oriented channel with given destination CID.",
    0x0410: "Returned when connection-oriented channel disconnected due to LE connection is dropped.",
    0x0412: "Returned when connection-oriented channel disconnected due to remote end send data even without credit.",
    0x0413: "Returned when connection-oriented channel disconnected due to remote end send flow control credits exceed 65535.",
    0x0414: "Returned when connection-oriented channel has run out of flow control credit and local application still trying to send data.",
    0x0415: "Returned when connection-oriented channel has not received connection response message within maximum timeout.",
    0x0416: "Returned when local host received a connection-oriented channel connection response with an invalid destination CID.",
    0x0417: "Returned when local host application tries to send a command which is not suitable for L2CAP channel's current state.",
    0x041B: "Flash reserved for PS store is full",
    0x041C: "PS key not found",
    0x041D: "Mismatched or insufficient security level",
    0x041E: "Encryption/decryption operation failed.",

    # Bluetooth controller status codes
    0x1002: "Connection does not exist, or connection open request was cancelled.",
    0x1005: "Pairing or authentication failed due to incorrect results in the pairing or authentication procedure. This could be due to an incorrect PIN or Link Key",
    0x1006: "Pairing failed because of missing PIN, or authentication failed because of missing Key",
    0x1007: "Controller is out of memory.",
    0x1008: "Link supervision timeout has expired.",
    0x1009: "Controller is at limit of connections it can support.",
    0x100A: "The Synchronous Connection Limit to a Device Exceeded error code indicates that the Controller has reached the limit to the number of synchronous connections that can be achieved to a device.",
    0x100B: "The ACL Connection Already Exists error code indicates that an attempt to create a new ACL Connection to a device when there is already a connection to this device.",
    0x100C: "Command requested cannot be executed because the Controller is in a state where it cannot process this command at this time.",
    0x100D: "The Connection Rejected Due To Limited Resources error code indicates that an incoming connection was rejected due to limited resources.",
    0x100E: "The Connection Rejected Due To Security Reasons error code indicates that a connection was rejected due to security requirements not being fulfilled, like authentication or pairing.",
    0x100F: "The Connection was rejected because this device does not accept the BD_ADDR. This may be because the device will only accept connections from specific BD_ADDRs.",
    0x1010: "The Connection Accept Timeout has been exceeded for this connection attempt.",
    0x1011: "A feature or parameter value in the HCI command is not supported.",
    0x1012: "Command contained invalid parameters.",
    0x1013: "User on the remote device terminated the connection.",
    0x1014: "The remote device terminated the connection because of low resources",
    0x1015: "Remote Device Terminated Connection due to Power Off",
    0x1016: "Local device terminated the connection.",
    0x1017: "The Controller is disallowing an authentication or pairing procedure because too little time has elapsed since the last authentication or pairing attempt failed.",
    0x1018: "The device does not allow pairing. This can be for example, when a device only allows pairing during a certain time window after some user input allows pairing",
    0x101A: "The remote device does not support the feature associated with the issued command.",
    0x101E: "Indicates that some LMP PDU / LL Control PDU parameters were invalid",
    0x101F: "No other error code specified is appropriate to use.",
    0x1022: "Connection terminated due to link-layer procedure timeout.",
    0x1023: "LL procedure has collided with the same transaction or procedure that is already in progress.",
    0x1025: "The requested encryption mode is not acceptable at this time.",
    0x1026: "Link key cannot be changed because a fixed unit key is being used.",
    0x1028: "LMP PDU or LL PDU that includes an instant cannot be performed because the instant when this would have occurred has passed.",
    0x1029: "It was not possible to pair as a unit key was requested and it is not supported.",
    0x102A: "LMP transaction was started that collides with an ongoing transaction.",
    0x102E: "The Controller cannot perform channel assessment because it is not supported.",
    0x102F: "The HCI command or LMP PDU sent is only possible on an encrypted link.",
    0x1030: "A parameter value requested is outside the mandatory range of parameters for the given HCI command or LMP PDU.",
    0x1037: "The IO capabilities request or response was rejected because the sending Host does not support Secure Simple Pairing even though the receiving Link Manager does.",
    0x1038: "The Host is busy with another pairing operation and unable to support the requested pairing. The receiving device should retry pairing again later.",
    0x1039: "The Controller could not calculate an appropriate value for the Channel selection operation.",
    0x103A: "Operation was rejected because the controller is busy and unable to process the request.",
    0x103B: "Remote device terminated the connection because of an unacceptable connection interval.",
    0x103C: "Advertising for a fixed duration completed or, for directed advertising, that advertising completed without a connection being created.",
    0x103D: "Connection was terminated because the Message Integrity Check (MIC) failed on a received packet.",
    0x103E: "LL initiated a connection but the connection has failed to be established. Controller did not receive any packets from remote end.",
    0x103F: "The MAC of the 802.11 AMP was requested to connect to a peer, but the connection failed.",
    0x1040: "The master, at this time, is unable to make a coarse adjustment to the piconet clock, using the supplied parameters. Instead the master will attempt to move the clock using clock dragging.",
    0x1042: "A command was sent from the Host that should identify an Advertising or Sync handle, but the Advertising or Sync handle does not exist.",
    0x1043: "Number of operations requested has been reached and has indicated the completion of the activity (e.g., advertising or scanning).",
    0x1044: "A request to the Controller issued by the Host and still pending was successfully canceled.",
    0x1045: "An attempt was made to send or receive a packet that exceeds the maximum allowed packet length.",
    0x1046: "Information was provided too late to the controller.",
    0x1047: "Information was provided too early to the controller.",
    0x1048: "Indicates that the result of the requested operation would yield too few physical channels.",

    # Bluetooth attribute status codes
    0x1101: "The attribute handle given was not valid on this server",
    0x1102: "The attribute cannot be read",
    0x1103: "The attribute cannot be written",
    0x1104: "The attribute PDU was invalid",
    0x1105: "The attribute requires authentication before it can be read or written.",
    0x1106: "Attribute Server does not support the request received from the client.",
    0x1107: "Offset specified was past the end of the attribute",
    0x1108: "The attribute requires authorization before it can be read or written.",
    0x1109: "Too many prepare writes have been queued",
    0x110A: "No attribute found within the given attribute handle range.",
    0x110B: "The attribute cannot be read or written using the Read Blob Request",
    0x110C: "The Encryption Key Size used for encrypting this link is insufficient.",
    0x110D: "The attribute value length is invalid for the operation",
    0x110E: "The attribute request that was requested has encountered an error that was unlikely, and therefore could not be completed as requested.",
    0x110F: "The attribute requires encryption before it can be read or written.",
    0x1110: "The attribute type is not a supported grouping attribute as defined by a higher layer specification.",
    0x1111: "Insufficient Resources to complete the request",
    0x1112: "The server requests the client to rediscover the database.",
    0x1113: "The attribute parameter value was not allowed.",
    0x1180: "When this is returned in a BGAPI response, the application tried to read or write the value of a user attribute from the GATT database.",
    0x11FC: "The requested write operation cannot be fulfilled for reasons other than permissions.",
    0x11FD: "The Client Characteristic Configuration descriptor is not configured according to the requirements of the profile or service.",
    0x11FE: "The profile or service request cannot be serviced because an operation that has been previously triggered is still in progress.",
    0x11FF: "The attribute value is out of range as defined by a profile or service specification.",

    # Bluetooth Security Manager Protocol status codes
    0x1201: "The user input of passkey failed, for example, the user cancelled the operation",
    0x1202: "Out of Band data is not available for authentication",
    0x1203: "The pairing procedure cannot be performed as authentication requirements cannot be met due to IO capabilities of one or both devices",
    0x1204: "The confirm value does not match the calculated compare value",
    0x1205: "Pairing is not supported by the device",
    0x1206: "The resultant encryption key size is insufficient for the security requirements of this device",
    0x1207: "The SMP command received is not supported on this device",
    0x1208: "Pairing failed due to an unspecified reason",
    0x1209: "Pairing or authentication procedure is disallowed because too little time has elapsed since last pairing request or security request",
    0x120A: "The Invalid Parameters error code indicates: the command length is invalid or a parameter is outside of the specified range.",
    0x120B: "Indicates to the remote device that the DHKey Check value received doesn't match the one calculated by the local device.",
    0x120C: "Indicates that the confirm values in the numeric comparison protocol do not match.",
    0x120D: "Indicates that the pairing over the LE transport failed due to a Pairing Request sent over the BR/EDR transport in process.",
    0x120E: "Indicates that the BR/EDR Link Key generated on the BR/EDR transport cannot be used to derive and distribute keys for the LE transport.",
    0x120F: "Indicates that the device chose not to accept a distributed key.",

    # Bluetooth Mesh status codes
    0x0501: "Returned when trying to add a key or some other unique resource with an ID which already exists",
    0x0502: "Returned when trying to manipulate a key or some other resource with an ID which does not exist",
    0x0503: "Returned when an operation cannot be executed because a pre-configured limit for keys, key bindings, elements, models, virtual addresses, provisioned devices, or provisioning sessions is reached",
    0x0504: 'Returned when trying to use a reserved address or add a "pre-provisioned" device using an address already used by some other device',
    0x0505: "In a BGAPI response, the user supplied malformed data; in a BGAPI event, the remote end responded with malformed or unrecognized data",
    0x0506: "An attempt was made to initialize a subsystem that was already initialized.",
    0x0507: "An attempt was made to use a subsystem that wasn't initialized yet. Call the subsystem's init function first.",
    0x0508: "Returned when trying to establish a friendship as a Low Power Node, but no acceptable friend offer message was received.",
    0x0509: "Provisioning link was unexpectedly closed before provisioning was complete.",
    0x050A: "An unrecognized provisioning PDU was received.",
    0x050B: "A provisioning PDU with wrong length or containing field values that are out of bounds was received.",
    0x050C: "An unexpected (out of sequence) provisioning PDU was received.",
    0x050D: "The computed confirmation value did not match the expected value.",
    0x050E: "Provisioning could not be continued due to insufficient resources.",
    0x050F: "The provisioning data block could not be decrypted.",
    0x0510: "An unexpected error happened during provisioning.",
    0x0511: "Device could not assign unicast addresses to all of its elements.",
    0x0512: "Returned when trying to reuse an address of a previously deleted device before an IV Index Update has been executed.",
    0x0513: "Returned when trying to assign an address that is used by one of the devices in the Device Database, or by the Provisioner itself.",
    0x0514: "Application key or publish address are not set",
    0x0515: "Application key is not bound to a model",

    # Bluetooth Mesh foundation status codes
    0x1301: "Returned when address in request was not valid",
    0x1302: "Returned when model identified is not found for a given element",
    0x1303: "Returned when the key identified by AppKeyIndex is not stored in the node",
    0x1304: "Returned when the key identified by NetKeyIndex is not stored in the node",
    0x1305: "Returned when The node cannot serve the request due to insufficient resources",
    0x1306: "Returned when the key identified is already stored in the node and the new NetKey value is different",
    0x1307: "Returned when the model does not support the publish mechanism",
    0x1308: "Returned when  the model does not support the subscribe mechanism",
    0x1309: "Returned when storing of the requested parameters failed",
    0x130A: "Returned when requested setting is not supported",
    0x130B: "Returned when the requested update operation cannot be performed due to general constraints",
    0x130C: "Returned when the requested delete operation cannot be performed due to general constraints",
    0x130D: "Returned when the requested bind operation cannot be performed due to general constraints",
    0x130E: "Returned when The node cannot start advertising with Node Identity or Proxy since the maximum number of parallel advertising is reached",
    0x130F: "Returned when the requested state cannot be set",
    0x1310: "Returned when an unspecified error took place",
    0x1311: "Returned when the NetKeyIndex and AppKeyIndex combination is not valid for a Config AppKey Update",

    # Wi-Fi Errors
    0x0B01: "Invalid firmware keyset",
    0x0B02: "The firmware download took too long",
    0x0B03: "Unknown request ID or wrong interface ID used",
    0x0B04: "The request is successful but some parameters have been ignored",
    0x0B05: "No Packets waiting to be received",
    0x0B08: "The sleep mode is granted",
    0x0B09: "The WFx does not go back to sleep",
    0x0B10: "The SecureLink MAC key was not found",
    0x0B11: "The SecureLink MAC key is already installed in OTP",
    0x0B12: "The SecureLink MAC key cannot be installed in RAM",
    0x0B13: "The SecureLink MAC key installation failed",
    0x0B14: "SecureLink key (re)negotiation failed",
    0x0B18: "The device is in an inappropriate state to perform the request",
    0x0B19: "The request failed due to regulatory limitations",
    0x0B1A: "The connection request failed because no suitable AP was found",
    0x0B1B: "The connection request was aborted by host",
    0x0B1C: "The connection request failed because of a timeout",
    0x0B1D: "The connection request failed because the AP rejected the device",
    0x0B1E: "The connection request failed because the WPA handshake did not complete successfully",
    0x0B1F: "The request failed because the retry limit was exceeded",
    0x0B20: "The request failed because the MSDU life time was exceeded",

    # MVP Driver and MVP Math status codes
    0x1501: "Critical fault",
    0x1502: "ALU operation output NaN",
    0x1503: "ALU numeric overflow",
    0x1504: "ALU numeric underflow",
    0x1505: "Overflow during array store",
    0x1506: "Underflow during array store conversion",
    0x1507: "Infinity encountered during array store conversion",
    0x1508: "NaN encountered during array store conversion",
    0x1512: "MATH NaN encountered",
    0x1513: "MATH Infinity encountered",
    0x1514: "MATH numeric overflow",
    0x1515: "MATH numeric underflow",

    # Zigbee status codes
    0x0C01: "Packet is dropped by packet-handoff callbacks",
    0x0C02: "The APS layer attempted to send or deliver a message and failed",
    0x0C03: "The maximum number of in-flight messages ::EMBER_APS_UNICAST_MESSAGE_COUNT has been reached",
    0x0C04: "The application is trying to delete or overwrite a binding that is in use",
    0x0C05: "The application is trying to overwrite an address table entry that is in use",
    0x0C06: "After moving, a mobile node's attempt to re-establish contact with the network failed",
    0x0C07: "The local node ID has changed. The application can get the new node ID by calling ::sl_zigbee_get_node_id()",
    0x0C08: "The chosen security level is not supported by the stack",
    0x0C09: "An error occurred when trying to encrypt at the APS Level",
    0x0C0A: "An error occurred when trying to encrypt at the APS Level",
    0x0C0B: "There was an attempt to form or join a network with security without calling ::sl_zigbee_set_initial_security_state() first",
    0x0C0C: "There was an attempt to broadcast a key switch too quickly after broadcasting the next network key. The Trust Center must wait at least a period equal to the broadcast timeout so that all routers have a chance to receive the broadcast of the new network key",
    0x0C0D: "The received signature corresponding to the message that was passed to the CBKE Library failed verification and is not valid",
    0x0C0E: "The message could not be sent because the link key corresponding to the destination is not authorized for use in APS data messages",
    0x0C0F: "The application tried to use a binding that has been remotely modified and the change has not yet been reported to the application",
    0x0C10: "The EUI of the Trust center has changed due to a successful rejoin after TC Swapout",
    0x0C11: "A Trust Center Swapout Rejoin has occurred without the EUI of the TC changing",
    0x0C12: "An attempt to generate random bytes failed because of insufficient random data from the radio",
    0x0C13: "A Zigbee route error command frame was received indicating that a source routed message from this node failed en route",
    0x0C14: "A Zigbee route error command frame was received indicating that a message sent to this node along a many-to-one route failed en route",
    0x0C15: "A critical and fatal error indicating that the version of the stack trying to run does not match with the chip it's running on",
    0x0C16: "The local PAN ID has changed. The application can get the new PAN ID by calling ::emberGetPanId()",
    0x0C17: "The channel has changed.",
    0x0C18: "The network has been opened for joining.",
    0x0C19: "The network has been closed for joining.",
    0x0C1A: "An attempt was made to join a Secured Network using a pre-configured key, but the Trust Center sent back a Network Key in-the-clear when an encrypted Network Key was required. (::EMBER_REQUIRE_ENCRYPTED_KEY)",
    0x0C1B: "An attempt was made to join a Secured Network, but the device did not receive a Network Key.",
    0x0C1C: "After a device joined a Secured Network, a Link Key was requested (::EMBER_GET_LINK_KEY_WHEN_JOINING) but no response was ever received.",
    0x0C1D: "An attempt was made to join a Secured Network without a pre-configured key, but the Trust Center sent encrypted data using a pre-configured key.",
    0x0C1E: "A Zigbee EZSP error has occured. Track the origin and corresponding EzspStatus for more info."
}

if __name__ == "__main__":
    main()
```
