## Supplementary material for "VesperFlow: An automated pipeline for decoding multisensor data from biologging tags": util.py

```
######################################################
NOTE: This code has been modified by Lily Broos (2026)
######################################################

"""
Application utility module
"""

# Copyright 2023 Silicon Laboratories Inc. www.silabs.com
#
# SPDX-License-Identifier: Zlib
#
# The licensor of this software is Silicon Laboratories Inc.
#
# This software is provided 'as-is', without any express or implied
# warranty. In no event will the authors be held liable for any damages
# arising from the use of this software.
#
# Permission is granted to anyone to use this software for any purpose,
# including commercial applications, and to alter it and redistribute it
# freely, subject to the following restrictions:
#
# 1. The origin of this software must not be misrepresented; you must not
#    claim that you wrote the original software. If you use this software
#    in a product, an acknowledgment in the product documentation would be
#    appreciated but is not required.
# 2. Altered source versions must be plainly marked as such, and must not be
#    misrepresented as being the original software.
# 3. This notice may not be removed or altered from any source distribution.

import argparse
import itertools
import logging
import os.path
import socket
import sys
import threading
import traceback
import struct
from datetime import datetime
import bgapi
from bgapi.connector import ConnectorException
import serial.tools.list_ports
from .status import Status

LOG_FORMAT_SINGLE = "%(asctime)s: %(levelname)s - %(message)s"
LOG_FORMAT = "%(asctime)s: %(name)s %(levelname)s - %(message)s"
BT_XAPI = os.path.join(os.path.dirname(__file__), "../api/sl_bt.xapi")
BTMESH_XAPI = os.path.join(os.path.dirname(__file__), "../api/sl_btmesh.xapi")


GAPP_CMD_GETNFILES = 	    1
GAPP_CMD_FINDFILE =		    2
GAPP_CMD_UPDATERECIPE =	    3
GAPP_CMD_UPDATERCOMMIT =	4
GAPP_CMD_STOPDNLD =	        6
GAPP_CMD_STARTDNLD =	    7
GAPP_CMD_RELEASE =		    8
GAPP_CMD_GETTIME =		    9
GAPP_CMD_SETTIME =		    0x0A
GAPP_CMD_GETSTATUS =		0x0F
GAPP_CMD_SETCFG =		    0x10
CONFIG_REGS_TXPWR = 0x03											
GAPP_CMD_GETCFG =			0x11											
GAPP_CMD_GETPAGE =			0x12


# Patch bgapi package to use a modified version of CommandFailedError
class CommandFailedError(bgapi.bglib.CommandError):
    """ Convert errorcode into Status object. """
    def __init__(self, response, command=None):
        self.errorcode = Status(response._errorcode)
        self.response = response
        self.command = command
        msg = f"Command failed with result {self.errorcode:#06x}: '{self.errorcode}'"
        if command:
            msg += f"\n> {command}\n< {response}"
        super().__init__(msg)

bgapi.bglib.CommandFailedError = CommandFailedError

class GenericApp(threading.Thread):
    """ Generic application class. """
    _id = itertools.count(0)
    def __init__(self, connector, apis):
        self.id = next(self._id)
        self.lib = bgapi.BGLib(connector, apis)
        self.log = logging.getLogger(f"{type(self).__name__}#{self.id}")
        # Set the ready event in the child classes to feed the watchdog
        self.ready = threading.Event()
        self._run = threading.Event()
        super().__init__()

    def event_handler(self, evt):
        """ Public event handler to perform user actions. Meant to be overridden by child classes. """

    def _event_handler(self, evt):
        """ Private event handler to perform internal actions. """
        print("evt =", evt)
        print("evt._str =", evt._str)
        print("evt == 'bt_evt_system_boot' ?", evt == "bt_evt_system_boot")

    def run(self):
        """ Main execution loop of the application. """
        self._run.set()
        exit_code = 0

        self.log.info("Open device")
        try:
            self.lib.open()
        except ConnectorException as err:
            self.log.error("%s", err)
            sys.exit(-1)

        # Start watchdog in the background
        threading.Thread(target=self.watchdog, daemon=True).start()

        # Enter main program loop.
        while self._run.is_set():
            try:
                # The timeout is needed to get the KeyboardInterrupt.
                # On Windows hosts, timeout is needed in both threaded and non-threaded modes.
                # On POSIX hosts, timeout is needed only in threaded mode.
                # The timeout value is a tradeoff between CPU load and KeyboardInterrupt response time.
                # timeout=None: minimal CPU usage, KeyboardInterrupt not recognized until the next event.
                # timeout=0: maximal CPU usage, KeyboardInterrupt recognized immediately.
                # See the documentation of Queue.get method for details.
                evt = self.lib.get_event(timeout=0.1)
                if evt is None:
                    continue
                # Convert event parameters with errorcode datatype into Status objects
                for param in evt._apinode.params:
                    if param.datatype.name == "errorcode":
                        value = getattr(evt, param.name)
                        setattr(evt, param.name, Status(value))
                self._event_handler(evt)
                if not self.ready.is_set():
                    # Unexpected events may happen if the previous host execution aborted and the
                    # target device continues emitting events. Therefore, calling application event
                    # handlers should be prevented before the device is ready.
                    continue
                self.event_handler(evt)
                # Call dedicated event callback if available.
                event_callback = getattr(self, evt._str, None)
                if event_callback is not None:
                    event_callback(evt)
            except bgapi.bglib.CommandFailedError as err:
                # Get additional info from trace.
                trace = traceback.extract_tb(sys.exc_info()[-1])[-3]
                self.log.error("%s", err)
                self.log.error("  File '%s', line %d, in %s", trace.filename, trace.lineno, trace.name)
                self.log.error("    %s", trace.line)
                self._run.clear()
                exit_code = -1
            except KeyboardInterrupt:
                self.log.info("User interrupt")
                self._run.clear()

        self.log.info("Close device")
        self.lib.close()
        sys.exit(exit_code)

    def stop(self):
        """ Terminate main execution loop. """
        self._run.clear()

    def reset(self):
        """ Reset device, meant to be overridden by child classes. """

    def watchdog(self):
        """ Device supervisor task. """
        # Wait 1 second for the device to be ready before sending the first reset command
        if self.ready.wait(1):
            return
        retry = 0
        while retry < 3:
            self.log.info("Resetting device (%d)...", retry)
            retry += 1
            self.reset()
            # Wait 10 seconds for the device to be ready before sending the next reset command
            if self.ready.wait(10):
                return
        self.log.error("Device unreachable.")
        self.stop()

class BluetoothApp(GenericApp):
    """ Application class for Bluetooth devices. """
    def __init__(self, connector, apis=BT_XAPI):
        self.address = None
        self.address_type = None
        super().__init__(connector=connector, apis=apis)

    def _event_handler(self, evt):
        """ Internal Bluetooth event handler. """
        if evt == "bt_evt_system_boot":
            # Feed the watchdog
            self.ready.set()
            # Check Bluetooth stack version
            version = "{major}.{minor}.{patch}".format(**vars(evt))
            self.log.info("Bluetooth stack booted: v%s-b%s", version, evt.build)
            if version != self.lib.bt.__version__:
                self.log.warning("BGAPI version mismatch: %s (target) != %s (host)", version, self.lib.bt.__version__)
            # Get Bluetooth address
            _, self.address, self.address_type = self.lib.bt.system.get_identity_address()
            self.log.info("Bluetooth %s address: %s",
                "static random" if self.address_type else "public device",
                self.address)

    def reset(self):
        """ Reset Bluetooth device. """
        self.lib.bt.system.reboot()

class BtMeshApp(GenericApp):
    """ Application class for Bluetooth mesh devices """
    def __init__(self, connector, apis=[BT_XAPI, BTMESH_XAPI]):
        super().__init__(connector=connector, apis=apis)

    def _event_handler(self, evt):
        """ Internal Bluetooth event handler. """
        if evt == "bt_evt_system_boot":
            # Feed the watchdog
            self.ready.set()
           
            # Check Bluetooth stack version
            version = "{major}.{minor}.{patch}".format(**vars(evt))
            self.log.info("Bluetooth stack booted: v%s-b%s", version, evt.build)
            if version != self.lib.bt.__version__:
                self.log.warning("BGAPI version mismatch: %s (target) != %s (host)", version, self.lib.bt.__version__)
            # Initialize Bluetooth Mesh device
            self.lib.btmesh.node.init()

    def reset(self):
        """ Reset for Bluetooth mesh device """
        self.lib.bt.system.reboot()

class CustomHelpFormatter(argparse.ArgumentDefaultsHelpFormatter,
                          argparse.RawDescriptionHelpFormatter):
    """ Combination of help formatters. """

class ArgumentParser(argparse.ArgumentParser):
    """ Custom argument parser for GenericApp and its derivatives """
    def __init__(self, *args, single_mode=True, epilog=None, formatter_class=CustomHelpFormatter, **kwargs):
        self.single_mode = single_mode
        self.cpc_options = hasattr(bgapi, 'CpcConnector')
        if self.single_mode:
            nargs = "?"
            cpc_const = []
            examples = (
                "examples:"
                "\n  %(prog)s                 Try to autodetect serial port"
                "\n  %(prog)s COM4            Open serial port on Windows"
                "\n  %(prog)s /dev/ttyACM0    Open serial port on POSIX"
                "\n  %(prog)s 192.168.1.10    Open TCP port")
            if self.cpc_options:
                examples += "\n  %(prog)s -c              Open default CPC daemon instance"
                examples += "\n  %(prog)s -c cpcd_1       Open CPC daemon instance"
        else:
            nargs = "*"
            cpc_const = None
            examples = (
                "examples:"
                "\n  %(prog)s                                        Try to autodetect all serial ports"
                "\n  %(prog)s COM4 COM5 COM6 COM7 COM8               Open serial ports on Windows"
                "\n  %(prog)s /dev/ttyACM0 /dev/ttyACM1              Open serial ports on POSIX"
                "\n  %(prog)s 192.168.1.10 192.168.1.11              Open TCP ports"
                "\n  %(prog)s /dev/ttyACM0 192.168.1.10              Open serial port and TCP port")
            if self.cpc_options:
                examples += "\n  %(prog)s -c cpcd_0 cpcd_1 cpcd_2                Open CPC daemon instances"
                examples += "\n  %(prog)s /dev/ttyACM0 192.168.1.10 -c cpcd_1    Open serial port, TCP port and CPC daemon instance"
        if epilog is None:
            epilog = examples
        else:
            epilog = examples + "\n" + epilog

        super().__init__(*args, epilog=epilog, formatter_class=formatter_class, **kwargs)

        self.add_argument(
            "conn",
            nargs=nargs,
            help="Serial or TCP connection parameter. See the examples for details.")
        if self.cpc_options:
            self.add_argument(
                "-c", "--cpc",
                nargs=nargs,
                help="CPC instance",
                const=cpc_const)
            self.add_argument(
                "--cpc_lib_path",
                help="CPC shared library path",
                default="/usr/local/lib/libcpc.so")
            self.add_argument(
                "--cpc_tracing",
                help="Enable CPC tracing",
                action="store_true")
        self.add_argument(
            "-l", "--log",
            type=str.upper,
            choices=logging._nameToLevel.keys(),
            help="Log level",
            default="INFO")
        self.add_argument(
            "--robust",
            help="Enable robust communication",
            action="store_true")
        self.add_argument(
            "--no_crc",
            dest="robust_crc",
            help="Disable CRC checking for robust communication. Ignored if robust communication is disabled.",
            action="store_false")

    def parse_args(self, *args, **kwargs):
        """ Implement special argument parsing rules """
        args = super().parse_args(*args, **kwargs)
        # Propagate single_mode to the arguments
        args.single_mode = self.single_mode
        # Configure logging
        if args.single_mode:
            log_format = LOG_FORMAT_SINGLE
        else:
            log_format = LOG_FORMAT
        logging.basicConfig(level=args.log, format=log_format)
        # Check connection parameters
        if not self.cpc_options:
            # cpc attribute is always granted
            args.cpc = None
        if args.cpc and args.single_mode:
            # Use list representation in single mode too
            args.cpc = [args.cpc]
        if not args.cpc and args.cpc is not None:
            # Use default CPC instance if no argument provided
            args.cpc = ["cpcd_0"]
        if args.cpc and not os.path.exists(args.cpc_lib_path):
            self.print_usage()
            print(f"{self.prog}: error: CPC library doesn't exist at {args.cpc_lib_path}")
            sys.exit(-1)
        if args.cpc and args.conn and args.single_mode:
            self.print_usage()
            print(f"{self.prog}: error: Too many connections specified:"
                f" -c {args.cpc[0]}, {args.conn}")
            sys.exit(-1)
        if args.conn and args.single_mode:
            # Use list representation in single mode too
            args.conn = [args.conn]
        if not args.conn and not args.cpc:
            # Try to autodetect serial device if no input is provided
            args.conn = get_device_list()
            if not args.conn:
                self.print_usage()
                print(f"{self.prog}: error: No serial device found."
                    " Please specify connection explicitly.")
                sys.exit(-1)
            elif args.single_mode and len(args.conn) > 1:
                self.print_usage()
                print(f"{self.prog}: error: {len(args.conn)} serial devices found:"
                    f"{', '.join(args.conn)}. Please specify connection explicitly.")
                sys.exit(-1)
        return args

def get_connector(args=None):
    """ Return CPC, serial or socket connector instance from arguments. """
    if args is None:
        args = ArgumentParser().parse_args()
    connector = []
    if args.cpc:
        for cpc in args.cpc:
            try:
                cpc_conn = bgapi.CpcConnector(
                    lib_path=args.cpc_lib_path,
                    cpc_instance=cpc,
                    tracing=args.cpc_tracing)
            except ConnectorException as err:
                logging.error("%s", err)
                logging.error("Is CPC daemon instance '%s' running?", cpc)
                sys.exit(-1)
            connector.append(cpc_conn)
    if args.conn:
        connector += [connector_from_str(conn) for conn in args.conn]
    if args.robust:
        # Enable robust layer on all connectors
        connector = [bgapi.RobustConnector(conn, args.robust_crc) for conn in connector]
    if args.single_mode:
        return connector[0]
    return connector

def get_device_list():
    """ Find Segger J-Link devices based on USB vendor ID. """
    return [com.device for com in serial.tools.list_ports.comports() if com.vid == 0x1366]

def connector_from_str(param):
    """ Return a serial or socket connector instance from a string parameter.

    This function is optimized for Silicon Labs development boards with default parameters.
    For non-default settings use the SerialConnector and SocketConnector constructors directly.
    """
    connector_type = bgapi.SocketConnector
    try:
        # Check for a valid IPv4 address.
        socket.inet_aton(param)
        # Append WSTK serial port number.
        param = (param, 4901)
    except OSError:
        # Assume serial port.
        connector_type = bgapi.SerialConnector
    return connector_type(param)

def find_service_in_advertisement(adv_data, uuid):
    """ Find service with the given UUID in the advertising data. """
    if len(uuid) != 2 and len(uuid) != 16:
        raise ValueError("Invalid UUID length.")
    # Incomplete List of 16 or 128-bit  Service Class UUIDs.
    incomplete_list = 0x02 if len(uuid) == 2 else 0x06
    # Complete List of 16 or 128-bit  Service Class UUIDs.
    complete_list = 0x03 if len(uuid) == 2 else 0x07

    # Parse advertisement packet.
    i = 0
    while i < len(adv_data):
        try:
            ad_field_length = adv_data[i]
            ad_field_type = adv_data[i + 1]
            # Find AD types of interest.
            if ad_field_type in (incomplete_list, complete_list):
                ad_uuid_count = int((ad_field_length - 1) / len(uuid))
                # Compare each UUID to the service UUID to be found.
                for j in range(ad_uuid_count):
                    start_idx = i + 2 + j*len(uuid)
                    # Get UUID from AD data.
                    ad_uuid = adv_data[start_idx: start_idx + len(uuid)]
                    if ad_uuid == uuid:
                        return True
            # Advance to the next AD structure.
            i += ad_field_length + 1
        except IndexError:
            # Malformed advertising data
            return False
    # UUID not found.
    return False

from datetime import datetime, timezone

def unix_timestamp_to_datetime(unix_timestamp):
    #Convert a Unix timestamp (seconds since epoch) into a timezone-aware UTC datetime.
    dt = datetime.fromtimestamp(unix_timestamp, tz=timezone.utc)
    return dt

def find_name_in_advertisement(adv_data, name):
    """ Find service with the given UUID in the advertising data. """
    if len(name) != 3 :
        raise ValueError("Invalid Name length.")
    
    complete_name = 0x09
    # Parse advertisement packet.
    i = 0
    while i < len(adv_data):
        try:
            ad_field_length = adv_data[i]
            ad_field_type = adv_data[i + 1]
            # Find AD types of interest.
            if ad_field_type == complete_name:
                ad_uuid_count = int((ad_field_length - 1) / len(name))
                # Compare each name to the service name to be found.
                for j in range(ad_uuid_count):
                    start_idx = i + 2 + j*len(name)
                    # Get name from AD data.
                    ad_name = adv_data[start_idx: start_idx + len(name)].decode(encoding="utf-8")
                    if ad_name == name:
                        return True
            # Advance to the next AD structure.
            i += ad_field_length + 1
        except IndexError:
            # Malformed advertising data
            return False
    # name not found.
    return False

def find_logs_in_advertisement(adv_data):  # [bool, addr, datetime, mode, type, txpwr, batcap, logs]
    mfr_data = 0xff

    # Parse advertisement packet.
    i = 0
    while i < len(adv_data):
        try:
            ad_field_length = adv_data[i]
            ad_field_type = adv_data[i + 1]
            # Find AD types of interest.
            if ad_field_type == mfr_data and ad_field_length == 21:
                start_idx = i + 2 + 2

                rtxper = int.from_bytes(adv_data[start_idx: start_idx+1], byteorder='little', signed=True)
                start_idx += 1
                rbatcap = int.from_bytes(adv_data[start_idx: start_idx+1], byteorder='little', signed=False)
                start_idx += 1

                unixtd = struct.unpack('<I', adv_data[start_idx: start_idx+4])[0] 

                rdt = unix_timestamp_to_datetime(unixtd)
                start_idx += 4
                revaddr = reversed(adv_data[start_idx: start_idx+6])
                addr = ''.join('{:02x}:'.format(x) for x in revaddr) 
                addr = addr[:-1]
                start_idx += 6
                mode = int.from_bytes(adv_data[start_idx: start_idx+1], byteorder='little', signed=False)
                start_idx += 1
                rtype = int.from_bytes(adv_data[start_idx: start_idx+1], byteorder='little', signed=False)
                start_idx += 1
                rlogs = struct.unpack('<I', adv_data[start_idx: start_idx+4])[0]
                start_idx += 4
                return (True, addr, rdt, mode, rtype, rtxper, rbatcap, rlogs)
            # Advance to the next AD structure.
            i += ad_field_length + 1
        except IndexError:
            # Malformed advertising data
            return (False, '', datetime.now(timezone.utc), 0, 0, 0, 0, 0)
    # Not found.
    return (False, '', datetime.now(timezone.utc), 0, 0, 0, 0, 0)


def bytes_to_datetime(byte_array):
    # Unpack the bytes into the corresponding fields
    # Define the C struct format
    struct_format = 'BBBBBBH'
    unpacked_data = struct.unpack(struct_format, byte_array)

    # Extract the individual fields
    year, month, day, hour, minute, second, millisecond = unpacked_data

    # Convert the year to a full year (assuming base year is 2000)
    year += 1900
    month += 1

    # Create and return a datetime object
    return datetime(year, month, day, hour, minute, second, millisecond * 1000)

def datetime_to_bytes(dt):
    # Define the C struct format
    struct_format = 'BBBBBBH'
    # Extract the individual fields from the datetime object
    year = dt.year - 1900  # Assuming base year is 1900
    month = dt.month - 1
    day = dt.day
    hour = dt.hour
    minute = dt.minute
    second = dt.second
    millisecond = int(dt.microsecond / 1000)  # Convert microseconds to milliseconds

    # Pack the fields into a byte array
    byte_array = struct.pack(struct_format, year, month, day, hour, minute, second, millisecond)

    return byte_array


def curdatetime_to_bytes():
    return datetime_to_bytes(datetime.utcnow()) #Sets time to UTC


def encode_set_rtc_cmd():
    return bytes([GAPP_CMD_SETTIME]) + (curdatetime_to_bytes())

def encode_set_tx_power_cmd(tx_dbm):
    """
    Encode command to change TX power.
    tx_dbm: TX power in dBm (e.g. -3, 0, 6)
    """

    #GAPP_CMD_SETCFG = 0x10
    #CONFIG_REGS_TXPWR = 0x03

    value = int(tx_dbm * 10) # convert dBm → dBm*10
    value_bytes = value.to_bytes(2, byteorder="little", signed=True) # convert to signed 16-bit little endian

    cmd = bytearray(28)
    cmd[0] = GAPP_CMD_SETCFG

    # register ID
    cmd[1] = CONFIG_REGS_TXPWR & 0xFF
    cmd[2] = (CONFIG_REGS_TXPWR >> 8) & 0xFF

    # register value
    cmd[3] = value_bytes[0]
    cmd[4] = value_bytes[1]

    return bytes(cmd)

def encode_get_rtc_cmd():
    existing_bytes = b'\x00\x00\x00\x00\x00\x00\x00\x00\x00\x00\x00\x00\x00\x00\x00\x00\x00\x00\x00\x00\x00\x00\x00\x00\x00\x00\x00\x00'
    return bytes([GAPP_CMD_GETTIME]) + (existing_bytes)

def decode_log(addr, array_of_bytes):
    r = False

    start_time = datetime.now(timezone.utc)
    seconds_duration = 0

    unit1 = addr
    unit2 = '00:00:00:00:00:00'
    rssi = -999
    tx_power = -999

    if(len(array_of_bytes) >= 16):
        i = 0
        unixtd = struct.unpack('<I', array_of_bytes[i: i+4])[0] 
        start_time = unix_timestamp_to_datetime(unixtd)
        i+=4
        seconds_duration = struct.unpack('<I', array_of_bytes[i: i+4])[0] 
        i+=4
        revaddr = reversed(array_of_bytes[i: i+6])
        unit2 = ''.join('{:02x}:'.format(x) for x in revaddr) 
        unit2 = unit2[:-1]
        i += 6
        rssi = int.from_bytes(array_of_bytes[i: i+1], byteorder='little', signed=True)
        i += 1
        tx_power = int.from_bytes(array_of_bytes[i: i+1], byteorder='little', signed=True)
        i += 1
        r = True
    
    return (r, unit1, unit2, rssi, tx_power, start_time, seconds_duration)
```
