## Supplementary material for "VesperFlow: An automated pipeline for decoding multisensor data from biologging tags": app.py

```
######################################################
NOTE: This code has been modified by Lily Broos (2026)
######################################################

#!/usr/bin/env python3
"""
Thermometer Client NCP-host Example Application.
"""

# Copyright 2021 Silicon Laboratories Inc. www.silabs.com
#
# SPDX-License-Identifier: Zlib
#
# The licensor of this software is Silicon Laboratories Inc.
#
# This software is provided 'as-is', without any express or implied
# warranty. In no event will the authors be held liable for any damages
# arising from the use of this software.
#
# Permission is granted to anyone to use this software for any purpose,
# including commercial applications, and to alter it and redistribute it
# freely, subject to the following restrictions:
#
# 1. The origin of this software must not be misrepresented; you must not
#    claim that you wrote the original software. If you use this software
#    in a product, an acknowledgment in the product documentation would be
#    appreciated but is not required.
# 2. Altered source versions must be plainly marked as such, and must not be
#    misrepresented as being the original software.
# 3. This notice may not be removed or altered from any source distribution.

from dataclasses import dataclass
import os.path
import sys
import time
import csv
from datetime import datetime

BASE_DIR = r"C:\yourpathway"
os.makedirs(BASE_DIR, exist_ok=True)

def is_file_locked(file_path):
    """Check if file is locked (e.g., opened in Excel)."""
    if not os.path.exists(file_path):
        return False
    try:
        with open(file_path, mode='a'):
            return False
    except PermissionError:
        return True

def get_log_file():
    """Return the daily log file path YYYYMMDD_logs.csv in BASE_DIR."""
    from datetime import datetime, timezone
    today = datetime.now(timezone.utc).strftime("%Y%m%d")
    return os.path.join(BASE_DIR, f"{today}_logs.csv")


sys.path.append(os.path.join(os.path.dirname(__file__), ".."))

from common.util import ArgumentParser, BluetoothApp, get_connector, find_service_in_advertisement, find_name_in_advertisement, find_logs_in_advertisement, curdatetime_to_bytes, encode_set_rtc_cmd, encode_set_tx_power_cmd, decode_log
import common.status as status


# Constants
DEVINFO_SERVICE = b"\x0A\x18"
MANUF_NAME_CHAR = b"\x29\x2a"
SYSTEM_ID_CHAR  = b"\x23\x2a"
SERIAL_NUM_CHAR = b"\x25\x2a"
HW_ID_CHAR      = b"\x27\x2a"
FW_ID_CHAR      = b"\x26\x2a"
MODEL_CHAR      = b"\x24\x2a"


PRX_SERVICE     = b"\x74\xc8\x5c\x56\x4a\x35\x29\x9c\xda\x47\x13\xbe\x76\x95\xe5\x01"
PRX_DATACP_CHAR = b"\x74\x0b\x4c\xb6\xda\xf7\xb3\x9f\xac\x4f\xce\x82\xa5\xf8\x93\x80"
PRX_LOG_CHAR    = b"\xac\x3e\x47\x46\x98\xa3\x6d\xa1\xa3\x47\x39\x9f\x24\x69\x31\x1d"

OTA_SERVICE = b"\x0A\x18"

CONN_INTERVAL_MIN = 80    # 100 ms
CONN_INTERVAL_MAX = 100   # 125 ms (original code = 80)
CONN_SLAVE_LATENCY = 1    # no latency
CONN_TIMEOUT = 1000       # 1000 ms
CONN_MIN_CE_LENGTH = 0
CONN_MAX_CE_LENGTH = 65535

SCAN_INTERVAL = 16        # 10 ms
SCAN_WINDOW = 16          # 10 ms
SCAN_PASSIVE = 0

# The maximum number of connections has to match with the configuration on the target side.
SL_BT_CONFIG_MAX_CONNECTIONS = 5

@dataclass
class Connection:
    """ Connection representation """
    address: str
    address_type: int
    service_prx: int=None
    service_devinfo: int=None
    characteristic_prxcp: int=None
    characteristic_log: int=None
    characteristic_manuf: int=None
    characteristic_sysid: int=None
    characteristic_sernum: int=None
    characteristic_hwid: int=None
    characteristic_fwid: int=None
    characteristic_model: int=None
    _stage: int=0
    _chars: int=0

    mfg_str: str=None
    sysid_str: str=None
    sernum_str: str=None
    hwid_str: str=None
    fwid_str: str=None
    model_str: str=None
    newlogs = []
    logs_available: int=0


class App(BluetoothApp):
    """ Application derived from generic BluetoothApp. """
    def bt_evt_system_boot(self, evt):
        """ Bluetooth event callback

        This event indicates that the device has started and the radio is ready.
        Do not call any stack command before receiving this boot event!
        """
        # Set the default connection parameters for subsequent connections
        self.lib.bt.connection.set_default_parameters(
            CONN_INTERVAL_MIN,
            CONN_INTERVAL_MAX,
            CONN_SLAVE_LATENCY,
            CONN_TIMEOUT,
            CONN_MIN_CE_LENGTH,
            CONN_MAX_CE_LENGTH)
        # Start scanning - looking for thermometer devices
        self.lib.bt.scanner.start(
            self.lib.bt.scanner.SCAN_PHY_SCAN_PHY_1M,
            self.lib.bt.scanner.DISCOVER_MODE_DISCOVER_GENERIC)
        self.log.info("Scanning started...")
        self.conn_state = "scanning"
        self.connections = dict[int, Connection]()

    def bt_evt_scanner_legacy_advertisement_report(self, evt):
        """ Bluetooth event callback """
        if self.conn_state not in ["scanning", "idle"]:
            # Already connecting or connected, skip this advertisement
            self.log.debug("Skipping advertisement: already connecting or connected")
            return
        # Parse advertisement packets
        if (evt.event_flags & self.lib.bt.scanner.EVENT_FLAG_EVENT_FLAG_CONNECTABLE and
            evt.event_flags & self.lib.bt.scanner.EVENT_FLAG_EVENT_FLAG_SCANNABLE):
            # If a thermometer advertisement is found...
            if find_name_in_advertisement(evt.data, "PRX"):
                rval = find_logs_in_advertisement(evt.data)
                # Clear and readable UTC-aware printout
                if rval[0]:
                    print(f"{rval[1]} | Tag time (UTC): {rval[2].strftime('%Y-%m-%d %H:%M:%S')} UTC "
                          f"| Mode: {rval[3]} | Logs: {rval[7]}")
                else:
                    print("Invalid or malformed advertisement packet received")

            # Save logs_available as before
                self.logs_available = rval[7]
                from datetime import datetime, timezone
                if (rval[0] and (self.logs_available > 0 or rval[2].year != datetime.now(timezone.utc).year)):
                    if len(self.connections) < SL_BT_CONFIG_MAX_CONNECTIONS:
                        # then stop scanning for a while
                        self.lib.bt.scanner.stop()
                        time.sleep(0.2)                # Wait 200 ms to ensure scanner fully stops
                        # and connect to that device
                        self.lib.bt.connection.open(
                            evt.address,
                            evt.address_type,
                            self.lib.bt.gap.PHY_PHY_1M)
                        self.conn_state = "opening"

    def bt_evt_connection_opened(self, evt):
        """ Bluetooth event callback """
        self.log.info(f"Connection opened to {evt.address}")
        self.connections[evt.connection] = Connection(evt.address, evt.address_type)
        # Discover PRX service on the slave device
        self._stage = 0
        self._chars = 0
        self.newlogs = []
        self.address = evt.address
        self.lib.bt.gatt.discover_primary_services_by_uuid(evt.connection, DEVINFO_SERVICE)
        #self.lib.bt.gatt.discover_primary_services(evt.connection)
        self.conn_state = "discover_services_1"

    def bt_evt_gatt_service(self, evt):
        """ Bluetooth event callback """
        if(self._stage == 0):
            self.connections[evt.connection].service_devinfo = evt.service               
        elif(self._stage == 1): 
            self.connections[evt.connection].service_prx = evt.service
            

    def bt_evt_gatt_characteristic(self, evt):
        """ Bluetooth event callback """
        if(self._chars == 0):
            self.connections[evt.connection].characteristic_manuf = evt.characteristic
        elif(self._chars == 1): 
            self.connections[evt.connection].characteristic_sysid = evt.characteristic
        elif(self._chars == 2): 
            self.connections[evt.connection].characteristic_sernum = evt.characteristic
        elif(self._chars == 3): 
            self.connections[evt.connection].characteristic_hwid = evt.characteristic
        elif(self._chars == 4): 
            self.connections[evt.connection].characteristic_fwid = evt.characteristic
        elif(self._chars == 5): 
            self.connections[evt.connection].characteristic_model = evt.characteristic
        elif(self._chars == 10): 
            self.connections[evt.connection].characteristic_prxcp = evt.characteristic
        elif(self._chars == 11): 
            self.connections[evt.connection].characteristic_log = evt.characteristic


    def bt_evt_gatt_procedure_completed(self, evt):
        """ Bluetooth event callback """
        if evt.result != status.OK:
            address = self.connections[evt.connection].address
            self.log.error(f"GATT procedure for {address} completed with status {evt.result:#x}")
    
        # If stack ran out of resources, reset connection gracefully
            try:
                self.conn_state = "conn_close"
                self.lib.bt.connection.close(evt.connection)
            except Exception as e:
                self.log.warning(f"Connection close failed: {e}")
            return

        # If service discovery finished
        if self.conn_state == "discover_services_1":
            self._stage = 1
            self._chars = 4
            self.lib.bt.gatt.discover_primary_services_by_uuid(evt.connection, PRX_SERVICE)
            self.conn_state = "discover_characteristics_4"

        elif self.conn_state == "discover_services_2":
            self._chars = 0
            self.lib.bt.gatt.discover_characteristics_by_uuid(evt.connection, self.connections[evt.connection].service_devinfo, MANUF_NAME_CHAR)
            self.conn_state = "discover_characteristics_1"

        elif self.conn_state == "discover_characteristics_1":
            self._chars = 1
            self.lib.bt.gatt.discover_characteristics_by_uuid(evt.connection, self.connections[evt.connection].service_devinfo, SYSTEM_ID_CHAR)
            self.conn_state = "discover_characteristics_2"

        elif self.conn_state == "discover_characteristics_2":
            self._chars = 2
            self.lib.bt.gatt.discover_characteristics_by_uuid(evt.connection, self.connections[evt.connection].service_devinfo, SERIAL_NUM_CHAR)
            self.conn_state = "discover_characteristics_3"

        elif self.conn_state == "discover_characteristics_3":
            self._chars = 3
            self.lib.bt.gatt.discover_characteristics_by_uuid(evt.connection, self.connections[evt.connection].service_devinfo, HW_ID_CHAR)
            self.conn_state = "discover_characteristics_4"

        elif self.conn_state == "discover_characteristics_4":
            self._chars = 4
            self.lib.bt.gatt.discover_characteristics_by_uuid(evt.connection, self.connections[evt.connection].service_devinfo, FW_ID_CHAR)
            self.conn_state = "discover_characteristics_6"


        elif self.conn_state == "discover_characteristics_5":
            self._chars = 5
            self.lib.bt.gatt.discover_characteristics_by_uuid(evt.connection, self.connections[evt.connection].service_devinfo, MODEL_CHAR)
            self.conn_state = "discover_characteristics_6"

        elif self.conn_state == "discover_characteristics_6":
            self._chars = 10
            self.lib.bt.gatt.discover_characteristics_by_uuid(evt.connection, self.connections[evt.connection].service_prx, PRX_DATACP_CHAR)
            self.conn_state = "discover_characteristics_7"

        elif self.conn_state == "discover_characteristics_7":
            self._chars = 11
            self.lib.bt.gatt.discover_characteristics_by_uuid(evt.connection, self.connections[evt.connection].service_prx, PRX_LOG_CHAR)
            self.conn_state = "discover_characteristics_8"

        elif self.conn_state == "discover_characteristics_8":
            self._chars = 12
            self.conn_state = "discover_characteristics"
            self.conn_state = "read_fwid"
            self.lib.bt.gatt.read_characteristic_value(evt.connection, self.connections[evt.connection].characteristic_fwid)

        # If characteristic discovery finished
        elif self.conn_state == "read_fwid":
            self.conn_state = "write_rtc"
            self.log.info("Write RTC")
            self.lib.bt.gatt.write_characteristic_value(
                evt.connection, 
                self.connections[evt.connection].characteristic_prxcp, 
                encode_set_rtc_cmd())

        elif self.conn_state == "write_rtc":
            self.conn_state = "set_tx_power"
            
            self.lib.bt.gatt.write_characteristic_value(
                evt.connection,
                self.connections[evt.connection].characteristic_prxcp,
                encode_set_tx_power_cmd(-3)						##SET TX POWER HERE
            )

        elif self.conn_state == "set_tx_power":
            self.log.info(f"TX power set to -3 dBm")
            if self.logs_available > 0:
                self.conn_state = "read_logs"
                self.log.info("Read Logs")
                self.logs_available -= 1
                try:
                    self.lib.bt.gatt.read_characteristic_value(
                        evt.connection,
                        self.connections[evt.connection].characteristic_log
                    )
                except Exception as e:
                    self.log.error(f"GATT read failed for {self.connections[evt.connection].address}: {e}")
                    self.conn_state = "conn_close"
                    try:
                        self.lib.bt.connection.close(evt.connection)
                    except Exception as close_err:
                        self.log.warning(f"Failed to close connection cleanly: {close_err}")
                    return
            else:
                self.conn_state = "conn_close"
                self.lib.bt.connection.close(evt.connection)


        # If indication enable process finished
        elif self.conn_state == "read_logs":
            if(self.logs_available > 0):
                self.conn_state = "read_logs"
                self.log.info("Read Logs")
                self.logs_available -= 1
                try:
                    # Try reading the characteristic
                    self.lib.bt.gatt.read_characteristic_value(
                        evt.connection,
                        self.connections[evt.connection].characteristic_log
                    )
                except Exception as e:
                    self.log.error(f"GATT read failed for {self.connections[evt.connection].address}: {e}")
                    # Likely the device is out of range — close gracefully
                    self.conn_state = "conn_close"
                    try:
                        self.lib.bt.connection.close(evt.connection)
                    except Exception as close_err:
                        self.log.warning(f"Failed to close connection cleanly: {close_err}")
                    return
            else:
                self.conn_state = "conn_close"
                self.lib.bt.connection.close(evt.connection)


    def bt_evt_gatt_characteristic_value(self, evt):
        """ Bluetooth event callback - handle log data as it arrives """
        if self.conn_state == "read_logs":
            newlog = decode_log(self.address, evt.value)
            if newlog[0] is True:
                self.newlogs.append(newlog)  # keep in memory only

                file_path = get_log_file()

                try:
                    # Always try writing to the main logs.csv
                    with open(file_path, mode='a', newline='', encoding='utf-8') as file:
                        writer = csv.writer(file)
                        # If file is empty, write header
                        if os.path.getsize(file_path) == 0:
                            writer.writerow(['Status', 'Unit1', 'Unit2', 'RSSI', 'TX Power', 'Start Time', 'Seconds Duration'])
                        # Remove timezone info (+00:00) before writing
                        newlog = [
                            item.replace(tzinfo=None) if isinstance(item, datetime) else item
                            for item in newlog
                        ]
                        writer.writerow(newlog)
                        file.flush()
                        os.fsync(file.fileno())  # ensure it's really written to disk
                        self.log.info(f"Log written to {file_path}")

                except PermissionError:
                    # If logs.csv is locked (e.g., Excel is open), fall back to backup file
                    base, ext = os.path.splitext(file_path)
                    alt_file = f"{base}_backup_{int(time.time())}{ext}"
                    self.log.warning(f"{file_path} locked. Writing log to {alt_file}")

                    with open(alt_file, mode='a', newline='', encoding='utf-8') as file:
                        writer = csv.writer(file)
                        if os.path.getsize(alt_file) == 0:
                            writer.writerow(['Status', 'Unit1', 'Unit2', 'RSSI', 'TX Power', 'Start Time', 'Seconds Duration'])
                        # Remove timezone info (+00:00) before writing
                        newlog = [
                            item.replace(tzinfo=None) if isinstance(item, datetime) else item
                            for item in newlog
                        ]
                        writer.writerow(newlog)
                        file.flush()
                        os.fsync(file.fileno())
                        self.log.info(f"Log written to backup for {self.address}: {newlog}")


    def bt_evt_connection_closed(self, evt):
        """ Bluetooth event callback """
        address = self.connections[evt.connection].address
        self.log.info(f"Connection to {address} closed with reason {evt.reason:#x}: '{evt.reason}'")
    
        # Just clean up state (no log writing here anymore)
        del self.connections[evt.connection]
        self._stage = 0
        self._chars = 0
        self.logs_available = 0
        self.newlogs = []
        self.address = 0

        # Add a short cooldown before scanning again
        time.sleep(0.6)

        if self.conn_state != "scanning":
            # start scanning again to find new devices
            self.lib.bt.scanner.start(
                self.lib.bt.scanner.SCAN_PHY_SCAN_PHY_1M,
                self.lib.bt.scanner.DISCOVER_MODE_DISCOVER_GENERIC)
            self.conn_state = "scanning"

# Script entry point.
if __name__ == "__main__":
    parser = ArgumentParser(description=__doc__)
    args = parser.parse_args()
    connector = get_connector(args)
    # Instantiate the application.
    app = App(connector)
    # Running the application blocks execution until it terminates.
    app.run()
```
