## Supplementary material for "VesperFlow: An automated pipeline for decoding multisensor data from biologging tags": app.py

```
#!/usr/bin/env python3
"""
Empty NCP-host Example Application.
"""

# Copyright 2021 Silicon Laboratories Inc. www.silabs.com
#
# SPDX-License-Identifier: Zlib
#
# The licensor of this software is Silicon Laboratories Inc.
#
# This software is provided 'as-is', without any express or implied
# warranty. In no event will the authors be held liable for any damages
# arising from the use of this software.
#
# Permission is granted to anyone to use this software for any purpose,
# including commercial applications, and to alter it and redistribute it
# freely, subject to the following restrictions:
#
# 1. The origin of this software must not be misrepresented; you must not
#    claim that you wrote the original software. If you use this software
#    in a product, an acknowledgment in the product documentation would be
#    appreciated but is not required.
# 2. Altered source versions must be plainly marked as such, and must not be
#    misrepresented as being the original software.
# 3. This notice may not be removed or altered from any source distribution.

import os.path
import sys

sys.path.append(os.path.join(os.path.dirname(__file__), "../.."))
from common.util import ArgumentParser, BluetoothApp, get_connector

# Characteristic values
GATTDB_DEVICE_NAME = b"Empty Example"
GATTDB_MANUFACTURER_NAME_STRING = b"Silicon Labs"

class App(BluetoothApp):
    """ Application derived from generic BluetoothApp. """
    def bt_evt_system_boot(self, evt):
        """ Bluetooth event callback

        This event indicates that the device has started and the radio is ready.
        Do not call any stack command before receiving this boot event!
        """
        self.adv_handle = None
        self.gattdb_init()
        self.adv_start()

    def bt_evt_connection_opened(self, evt):
        """ Bluetooth event callback """
        self.log.info(f"Connection opened to {evt.address}")

    def bt_evt_connection_closed(self, evt):
        """ Bluetooth event callback """
        self.log.info(f"Connection closed with reason {evt.reason:#x}: '{evt.reason}'")
        self.adv_start()

    #####################################
    # Add further event callbacks here. #
    #####################################

    def gattdb_init(self):
        """ Initialize GATT database. """
        _, session = self.lib.bt.gattdb.new_session()

        # Generic Access
        _, service = self.lib.bt.gattdb.add_service(
            session,
            self.lib.bt.gattdb.SERVICE_TYPE_PRIMARY_SERVICE,
            0,
            b"\x00\x18"
        )
        # Device Name
        self.lib.bt.gattdb.add_uuid16_characteristic(
            session,
            service,
            self.lib.bt.gattdb.CHARACTERISTIC_PROPERTIES_CHARACTERISTIC_READ |
            self.lib.bt.gattdb.CHARACTERISTIC_PROPERTIES_CHARACTERISTIC_WRITE,
            0,
            0,
            b"\x00\x2A",
            self.lib.bt.gattdb.VALUE_TYPE_FIXED_LENGTH_VALUE,
            len(GATTDB_DEVICE_NAME),
            GATTDB_DEVICE_NAME
        )
        # Appearance
        self.lib.bt.gattdb.add_uuid16_characteristic(
            session,
            service,
            self.lib.bt.gattdb.CHARACTERISTIC_PROPERTIES_CHARACTERISTIC_READ,
            0,
            0,
            b"\x01\x2A",
            self.lib.bt.gattdb.VALUE_TYPE_FIXED_LENGTH_VALUE,
            2,
            b"\x00\x00"
        )
        self.lib.bt.gattdb.start_service(session, service)

        # Device Information
        _, service = self.lib.bt.gattdb.add_service(
            session,
            self.lib.bt.gattdb.SERVICE_TYPE_PRIMARY_SERVICE,
            0,
            b"\x0A\x18"
        )
        # Manufacturer Name String
        self.lib.bt.gattdb.add_uuid16_characteristic(
            session,
            service,
            self.lib.bt.gattdb.CHARACTERISTIC_PROPERTIES_CHARACTERISTIC_READ,
            0,
            0,
            b"\x29\x2A",
            self.lib.bt.gattdb.VALUE_TYPE_FIXED_LENGTH_VALUE,
            len(GATTDB_MANUFACTURER_NAME_STRING),
            GATTDB_MANUFACTURER_NAME_STRING
        )
        # System ID
        # Pad and reverse unique ID to get System ID.
        addr = self.address.split(":")
        system_id = bytes.fromhex("".join(addr[:3] + ["ff", "fe"] + addr[3:]))
        self.lib.bt.gattdb.add_uuid16_characteristic(
            session,
            service,
            self.lib.bt.gattdb.CHARACTERISTIC_PROPERTIES_CHARACTERISTIC_READ,
            0,
            0,
            b"\x23\x2A",
            self.lib.bt.gattdb.VALUE_TYPE_FIXED_LENGTH_VALUE,
            8,
            system_id
        )
        self.lib.bt.gattdb.start_service(session, service)

        ################################
        # Add further attributes here. #
        ################################

        self.lib.bt.gattdb.commit(session)

    def adv_start(self):
        """ Start advertising. """
        if self.adv_handle is None:
            # Create advertising set for the first call.
            _, self.adv_handle = self.lib.bt.advertiser.create_set()
            self.lib.bt.legacy_advertiser.generate_data(
                self.adv_handle,
                self.lib.bt.advertiser.DISCOVERY_MODE_GENERAL_DISCOVERABLE)
            # Set advertising interval to 100 ms.
            self.lib.bt.advertiser.set_timing(
                self.adv_handle,
                160,  # interval min
                160,  # interval max
                0,    # duration
                0)    # max events
        self.lib.bt.legacy_advertiser.start(
            self.adv_handle,
            self.lib.bt.legacy_advertiser.CONNECTION_MODE_CONNECTABLE)

# Script entry point.
if __name__ == "__main__":
    parser = ArgumentParser(description=__doc__)
    args = parser.parse_args()
    connector = get_connector(args)
    # Instantiate the application.
    app = App(connector)
    # Running the application blocks execution until it terminates.
    app.run()
```
