## Supplementary material for "VesperFlow: An automated pipeline for decoding multisensor data from biologging tags": README.md

# Bluetooth - Empty

Original C example: **Bluetooth - SoC Empty**

This example is ***almost*** empty. It implements a basic application to demonstrate how to handle
events and how to use the GATT database.

* A simple application is implemented in the event handler function that starts advertising on boot
    (and on *connection_closed* event). As a result, remote devices can find the device and connect
    to it.

* A simple GATT database is defined by adding Generic Access and Device Information services. As a
    result, remote devices can read basic information, such as the device name.
