## Supplementary material for "VesperFlow: An automated pipeline for decoding multisensor data from biologging tags": app.py

```
#!/usr/bin/env python3
"""
iBeacon NCP-host Example Application.
"""

# Copyright 2021 Silicon Laboratories Inc. www.silabs.com
#
# SPDX-License-Identifier: Zlib
#
# The licensor of this software is Silicon Laboratories Inc.
#
# This software is provided 'as-is', without any express or implied
# warranty. In no event will the authors be held liable for any damages
# arising from the use of this software.
#
# Permission is granted to anyone to use this software for any purpose,
# including commercial applications, and to alter it and redistribute it
# freely, subject to the following restrictions:
#
# 1. The origin of this software must not be misrepresented; you must not
#    claim that you wrote the original software. If you use this software
#    in a product, an acknowledgment in the product documentation would be
#    appreciated but is not required.
# 2. Altered source versions must be plainly marked as such, and must not be
#    misrepresented as being the original software.
# 3. This notice may not be removed or altered from any source distribution.

import os.path
import struct
import sys

sys.path.append(os.path.join(os.path.dirname(__file__), "../.."))
from common.util import ArgumentParser, BluetoothApp, get_connector

BEACON_ADV_DATA = struct.pack(">BBBBBHH16sHHb",    # Most of the unsigned short (H) values are big endian (>).
    # Flag bits - See Bluetooth Core Specification Supplement v10, Part A, Section 1.3 for more details on flags.
    2,            # Length of field.
    0x01,         # Type of field.
    0x04 | 0x02,  # Flags: LE General Discoverable Mode, BR/EDR is disabled.

    # Manufacturer specific data.
    26,   # Length of field.
    0xFF, # Type of field.

    # The first two data octets shall contain a company identifier code from
    # the Assigned Numbers - Company Identifiers document.
    # 0x004C = Apple
    # Little endian value, swap byte order.
    0x4C00,

    # Beacon type.
    # 0x0215 is iBeacon.
    0x0215,

    # 128 bit / 16 byte UUID
    b"\xE2\xC5\x6D\xB5\xDF\xFB\x48\xD2\xB0\x60\xD0\xF5\xA7\x10\x96\xE0",

    # Beacon major number.
    34987,

    # Beacon minor number.
    1025,

    # The Beacon's measured RSSI at 1 meter distance in dBm.
    -41
)

class App(BluetoothApp):
    """ Application derived from generic BluetoothApp. """
    def bt_evt_system_boot(self, evt):
        """ Bluetooth event callback

        This event indicates that the device has started and the radio is ready.
        Do not call any stack command before receiving this boot event!
        """
        # Create advertising set.
        _, adv_handle = self.lib.bt.advertiser.create_set()
        # Set custom advertising data.
        self.lib.bt.legacy_advertiser.set_data(
            adv_handle,
            0,               # packet type
            BEACON_ADV_DATA) # adv_data
        # Set advertising interval to 100 ms.
        self.lib.bt.advertiser.set_timing(
            adv_handle,
            160,  # interval min
            160,  # interval max
            0,    # duration
            0)    # max events
        # Start advertising in user mode and disable connections.
        self.lib.bt.legacy_advertiser.start(
            adv_handle,
            self.lib.bt.legacy_advertiser.CONNECTION_MODE_NON_CONNECTABLE)
        self.log.info("iBeacon started")

# Script entry point.
if __name__ == "__main__":
    parser = ArgumentParser(description=__doc__)
    args = parser.parse_args()
    connector = get_connector(args)
    # Instantiate the application.
    app = App(connector)
    # Running the application blocks execution until it terminates.
    app.run()
```
