## Supplementary material for "VesperFlow: An automated pipeline for decoding multisensor data from biologging tags": README.md

# Bluetooth - iBeacon

Original C example: **Bluetooth - SoC iBeacon**

An [iBeacon]((https://developer.apple.com/ibeacon/)) device is an implementation that sends
non-connectable advertisements in iBeacon format.
The iBeacon Service gives Bluetooth accessories a simple and convenient way to send iBeacon to
smartphones.
