## Supplementary material for "VesperFlow: An automated pipeline for decoding multisensor data from biologging tags": app.py

```
#!/usr/bin/env python3
"""
Network Co-Processor (NCP) tester host application.
Demonstrates the communication between an NCP host and NCP target using BGAPI
user messages, responses and events. Can be used as a starting point for
creating custom commands or for testing purposes.
"""

# Copyright 2023 Silicon Laboratories Inc. www.silabs.com
#
# SPDX-License-Identifier: Zlib
#
# The licensor of this software is Silicon Laboratories Inc.
#
# This software is provided 'as-is', without any express or implied
# warranty. In no event will the authors be held liable for any damages
# arising from the use of this software.
#
# Permission is granted to anyone to use this software for any purpose,
# including commercial applications, and to alter it and redistribute it
# freely, subject to the following restrictions:
#
# 1. The origin of this software must not be misrepresented; you must not
#    claim that you wrote the original software. If you use this software
#    in a product, an acknowledgment in the product documentation would be
#    appreciated but is not required.
# 2. Altered source versions must be plainly marked as such, and must not be
#    misrepresented as being the original software.
# 3. This notice may not be removed or altered from any source distribution.

import argparse
import enum
import os.path
import random
import sys

sys.path.append(os.path.join(os.path.dirname(__file__), "../.."))
from common.util import ArgumentParser, BluetoothApp, get_connector

DEFAULT_MESSAGE_LENGTH = 180
DEFAULT_INTERVAL_MS = 20

class UserCommandId(enum.IntEnum):
    """ IDs of the user commands """
    PERIODIC_ASYNC = 1
    PERIODIC_ASYNC_STOP = 2
    GET_BOARD_NAME = 3
    ECHO = 4

class App(BluetoothApp):
    """ Application derived from generic BluetoothApp. """
    def __init__(self, connector, args):
        super().__init__(connector)

        self.echo_mode = args.echo
        self.test_length = args.width
        self.test_interval = args.interval
        self.test_end = args.end
        self.test_iteration = 0

    def bt_evt_system_boot(self, evt):
        """ Bluetooth event callback """
        board_name = self.get_board_name()
        self.log.info(f"Board name: {board_name}")

        if self.echo_mode:
            self.log.info("Start testing with echo commands")
            self.log.info(f"Parameters: Width={self.test_length}, End={self.test_end}")
            # Note that there is no event processing until the end of this iteration.
            for i in range(self.test_end):
                print(f"\rTest iteration {i} ", end="")
                self.echo(self.test_length)
            print("")
            self.log.info("Test passed.")
            self.stop()
        else:
            self.log.info("Start testing with periodic asynchronous events")
            self.log.info(f"Parameters: Width={self.test_length}, Interval={self.test_interval}, End={self.test_end}")
            self.test_iteration = 0
            self.start_periodic_async_test(self.test_interval, self.test_length)

    def bt_evt_user_message_to_host(self, evt):
        """ Bluetooth event callback """
        user_event_id = evt.message[0]
        user_event_data = evt.message[1:]

        if user_event_id == UserCommandId.PERIODIC_ASYNC:
            print(f"\rTest iteration {self.test_iteration} ", end="")

            if len(evt.message) != self.test_length:
                raise RuntimeError(f"Unexpected event length. Expected: {self.test_length}. "
                                   f"Received: {len(evt.message)}.")

            # The expected event payload is derived from the iteration counter.
            expected_data = bytes([self.test_iteration % 256]) * (self.test_length - 1)
            if user_event_data != expected_data:
                raise RuntimeError(f"Unexpected event data. Expected: 0x{expected_data.hex()}. "
                                   f"Received: 0x{user_event_data.hex()}.")

            self.test_iteration += 1
            if self.test_iteration >= self.test_end:
                print("")
                self.log.info("Test passed.")
                self.stop_periodic_async_test()
                self.stop()
        else:
            # In this example, only one event ID is expected, but any number of event types may be defined.
            self.log.error(f"Unexpected event ID. Expected: {UserCommandId.PERIODIC_ASYNC}. "
                           f"Received: {user_event_id}.")
            return

    def get_board_name(self):
        """ Send user command to get the board name as string. """
        command = bytes([UserCommandId.GET_BOARD_NAME])
        _, response = self.lib.bt.user.message_to_target(command)
        return response.decode("ascii")

    def start_periodic_async_test(self, interval, length):
        """ Send user command to start emitting periodic asynchronous events. """
        command = bytes([UserCommandId.PERIODIC_ASYNC, interval, length])
        expected_response = bytes([UserCommandId.PERIODIC_ASYNC])
        _, response = self.lib.bt.user.message_to_target(command)
        if response != expected_response:
            raise RuntimeError(f"Unexpected response. Expected: 0x{expected_response.hex()}. "
                               f"Received: 0x{response.hex()}.")

    def stop_periodic_async_test(self):
        """ Send user command to stop emitting periodic asynchronous events. """
        command = bytes([UserCommandId.PERIODIC_ASYNC_STOP])
        expected_response = command
        _, response = self.lib.bt.user.message_to_target(command)
        if response != expected_response:
            raise RuntimeError(f"Unexpected response. Expected: 0x{expected_response.hex()}. "
                               f"Received: 0x{response.hex()}.")

    def echo(self, length):
        """ Send user command with random data as payload and expect the same payload as response. """
        command = bytes([UserCommandId.ECHO]) + random.randbytes(length)
        expected_data = command
        _, response = self.lib.bt.user.message_to_target(command)
        if response != expected_data:
            raise RuntimeError(f"Unexpected response. Expected: 0x{expected_data.hex()}. "
                               f"Received: 0x{response.hex()}.")

def check_int(min_value=None, max_value=None):
    """ Helper for checking integer range. """
    def check_int_helper(arg: str):
        value = int(arg, 0)
        if min_value is not None and value < min_value:
            raise argparse.ArgumentTypeError(f"Argument must be at least {min_value}")
        if max_value is not None and value > max_value:
            raise argparse.ArgumentTypeError(f"Argument must be at most {max_value}")
        return value
    return check_int_helper

def main():
    parser = ArgumentParser(description=__doc__)
    parser.add_argument(
        "--echo",
        action="store_true",
        help="Echo (i.e. synchronous) mode instead of asynchronous test."
    )
    parser.add_argument(
        "-w", "--width",
        type=check_int(1, 255),
        metavar="[1, 255]",
        default=DEFAULT_MESSAGE_LENGTH,
        help="Size of the test messages in bytes. " \
             "In case a test fails experiment decreasing this parameter."
    )
    parser.add_argument(
        "-i", "--interval",
        type=check_int(1, 255),
        metavar="[1, 255]",
        default=DEFAULT_INTERVAL_MS,
        help="Time interval between user events in milliseconds. " \
             "In case a test fails experiment increasing this parameter."
    )
    parser.add_argument(
        "-e", "--end",
        type=check_int(1),
        default=1000,
        help="Number of test messages to wait for before the test terminates."
    )
    args = parser.parse_args()

    # The default test parameters imitate a real use case with a relatively high load.
    # It's also possible to adjust these parameters in a way that results in a data rate
    # that exceeds the capabilities of the transport layer below the BGAPI protocol
    # (e.g., UART baud rate), and therefore, lead to the test to fail.
    # Note that the real data rate is higher because of the overhead of the BGAPI header.
    if not args.echo and args.width / args.interval > DEFAULT_MESSAGE_LENGTH / DEFAULT_INTERVAL_MS:
        print("WARNING: Requested data rate exceeds default data rate!")

    connector = get_connector(args)
    # Instantiate the application.
    app = App(connector, args)
    # Running the application blocks execution until it terminates.
    app.run()

# Script entry point.
if __name__ == "__main__":
    main()
```
