## Supplementary material for "VesperFlow: An automated pipeline for decoding multisensor data from biologging tags": README.md

# Bluetooth - NCP test

There are some applications that require high NCP bandwidth, e.g. scanning for advertising devices
in a crowded environment. In some situations NCP bandwidth can be a limiting factor, and sometimes
it's very challenging to reproduce the test environment. This example can help in these situations
by generating high NCP traffic.

This example has 2 working modes. In the default (asynchronous) mode, the host asks the target to
emit events with the given length and frequency. The event length is given in bytes, the frequency
is given as the interval between the events in milliseconds. The default test parameters imitate a
real use case with a relatively high load. It's also possible to adjust these parameters in a way
that results in a data rate that exceeds the capabilities of the transport layer below the BGAPI
protocol (e.g., UART baud rate), and therefore, lead to the test to fail.

When echo (synchronous) mode is requested, the host sends an echo command to the target and
receives the response with the same content.

In both working modes, the test will terminate if the number of the requested iterations is reached.

The default NCP target firmware implements these custom test commands, so it works out of the box.

This example can also be used as a reference to handle custom BGAPI commands and events. Please
note that implementing custom commands and events requires changes in the NCP target firmware too.

## Test evaluation

If the test passes, it means that the test messages arrived in the expected order with the expected
content.

If the test fails, further debugging is needed to find the root cause. Possible root causes might be
- insufficient NCP bandwidth
- insufficient host system resources (especially on low power single board computers)

In case of NCP bandwidth shortage, you can try increasing the NCP UART baud rate as described in
[this article](https://community.silabs.com/s/article/wstk-virtual-com-port-baudrate-setting?language=en_US).
