## Supplementary material for "VesperFlow: An automated pipeline for decoding multisensor data from biologging tags": central.py

```
#!/usr/bin/env python3
""" Roaming NCP-host Example Application for central devices.
"""

# Copyright 2023 Silicon Laboratories Inc. www.silabs.com
#
# SPDX-License-Identifier: Zlib
#
# The licensor of this software is Silicon Laboratories Inc.
#
# This software is provided 'as-is', without any express or implied
# warranty. In no event will the authors be held liable for any damages
# arising from the use of this software.
#
# Permission is granted to anyone to use this software for any purpose,
# including commercial applications, and to alter it and redistribute it
# freely, subject to the following restrictions:
#
# 1. The origin of this software must not be misrepresented; you must not
#    claim that you wrote the original software. If you use this software
#    in a product, an acknowledgment in the product documentation would be
#    appreciated but is not required.
# 2. Altered source versions must be plainly marked as such, and must not be
#    misrepresented as being the original software.
# 3. This notice may not be removed or altered from any source distribution.

from dataclasses import dataclass
import json
import logging
import os
import pathlib
import queue
import shutil
import subprocess
import sys
import time
import threading
import webbrowser
import bgapi

sys.path.append(os.path.join(os.path.dirname(__file__), "../.."))
from common.util import ArgumentParser, BluetoothApp, get_connector, find_service_in_advertisement
import common.status as status

# Virtual identity address of the access points
IDENTITY_ADDRESS = "DE:AD:BE:EF:12:34"
# Service UUID of interest: heart rate
GATT_SERVICE_UUID = b"\x0d\x18"
# Characteristic UUID of interest: heart rate measurement
GATT_CHARACTERISTIC_UUID = b"\x37\x2a"
# The maximum number of connections has to match with the configuration on the target side.
SL_BT_CONFIG_MAX_CONNECTIONS = 4
# Wait for boot event of an access point for this amount of time in seconds
BOOT_TIMEOUT = 2
# BLE connection attempts are aborted after this amount of time in seconds
CONNECTION_TIMEOUT = 3
# Duration of the scanning for advertising devices in seconds
SCANNING_TIMEOUT = 3
# Duration of the connection analysis in seconds
CONNECTION_ANALYSIS_TIMEOUT = 3
# Periodic scanning is performed with this time interval in seconds
SCANNING_PERIOD = 30
# Periodic RSSI measurements on connected devices are performed with this time interval in seconds
RSSI_MEASUREMENT_PERIOD = 10
# Minimum value for the connection event interval in units of 1.25 ms
CONNECTION_INTERVAL_MIN = 80
# Maximum value for the connection event interval in units of 1.25 ms
CONNECTION_INTERVAL_MAX = 100
# Path of the bonding database file
BONDING_DB_PATH = "bonding_db.json"
# Path of the network graph file
NETWORK_GRAPH_PATH = "graph.svg"

@dataclass
class Connection:
    """ Connection representation """
    address: str
    address_type: int
    bonding_data: dict
    opened: threading.Event
    rssi: int=None
    service: int=None
    characteristic: int=None
    notification: bool=False

class ApEvent:
    """ Generic event reported by the access point """

class ApEventConnectionOpened(ApEvent):
    """ Connection has been opened """

class ApEventConnectionClosed(ApEvent):
    """ Connection has been closed intentionally """

class ApEventConnectionLost(ApEvent):
    """ Connection has been closed accidentally """

class ApEventBondingDbChanged(ApEvent):
    """ Bonding database entry has changed """

@dataclass
class ApEventConnectionRssi(ApEvent):
    """ RSSI value measured on connected devices """
    ap_id: int
    connection: int
    rssi: int

@dataclass
class ApEventScanRssi(ApEvent):
    """ RSSI value measured on advertising devices """
    ap_id: int
    address: tuple
    rssi: int

@dataclass
class ApEventAnalyzerRssi(ApEvent):
    """ RSSI value measured by connection analyzer """
    ap_id: int
    rssi: int

class AccessPoint(BluetoothApp):
    """ Roaming Access Point """
    def __init__(self,
                 connector,
                 event_queue: queue.Queue,
                 identity_address=None,
                 identity_type=None):
        self._event_queue = event_queue
        self._identity_address = identity_address
        self._identity_type = identity_type
        self.ready = threading.Event()
        self.connections = dict[int, Connection]()
        self.analyzer = None
        self._rssi_thread = threading.Thread(target=self._rssi_task, daemon=True)
        self._rssi_thread.start()
        super().__init__(connector)

    def start_scan(self):
        """ Start scanning """
        self.lib.bt.scanner.start(
            self.lib.bt.scanner.SCAN_PHY_SCAN_PHY_1M,
            self.lib.bt.scanner.DISCOVER_MODE_DISCOVER_GENERIC)

    def stop_scan(self):
        """ Stop scanning """
        self.lib.bt.scanner.stop()

    def get_conn_params(self, conn_handle):
        """ Get connection parameters for connection analysis """
        return self.lib.bt.connection.get_scheduling_details(conn_handle)

    def start_analyzer(self, conn_params):
        """ Start connection analysis """
        if self.analyzer is not None:
            self.log.error("Analyzer is already running")
            return
        # Ignore the start_time_us parameter because of the delays caused by the NCP operation
        _, self.analyzer = self.lib.bt.connection_analyzer.start(
            conn_params.access_address,
            conn_params.crc_init,
            conn_params.interval,
            conn_params.supervision_timeout,
            conn_params.central_clock_accuracy,
            conn_params.central_phy,
            conn_params.peripheral_phy,
            conn_params.channel_selection_algorithm,
            conn_params.hop,
            conn_params.channel_map,
            conn_params.channel,
            conn_params.event_counter,
            0,
            self.lib.bt.connection_analyzer.FLAGS_RELATIVE_TIME)

    def stop_analyzer(self):
        """ Stop connection analysis """
        if self.analyzer is None:
            self.log.error("Analyzer is not running")
            return
        self.lib.bt.connection_analyzer.stop(self.analyzer)
        self.analyzer = None

    @property
    def connectable(self):
        """ Check if new connections can be established """
        return len(self.connections) < SL_BT_CONFIG_MAX_CONNECTIONS

    def connect(self, address, address_type=None, bonding_data=None, rssi=None):
        """ Connect to a device """
        if not self.connectable:
            raise RuntimeError("no more connection possible")
        if address_type is None:
            address_type = self.lib.bt.gap.ADDRESS_TYPE_PUBLIC_ADDRESS
        if bonding_data is None:
            bonding_data = {}
        _, connection = self.lib.bt.connection.open(
            address,
            address_type,
            self.lib.bt.gap.PHY_PHY_1M)
        self.connections[connection] = Connection(address,
                                                  address_type,
                                                  bonding_data,
                                                  threading.Event(),
                                                  rssi)
        if not self.connections[connection].opened.wait(timeout=CONNECTION_TIMEOUT):
            self.log.warning(f"failed to open connection to {address}")
            self.disconnect(connection)

    def disconnect(self, conn_handle):
        """ Close connection """
        try:
            self.lib.bt.connection.close(conn_handle)
        except bgapi.bglib.CommandFailedError as err:
            # Connection may already be closed at this point.
            if err.errorcode == status.INVALID_HANDLE:
                self.log.warning("connection '%d' already closed", conn_handle)
            else:
                self.log.error("failed to close connection '%d' with status %#x: '%s'",
                               conn_handle, err.errorcode, err.errorcode)

    def _rssi_task(self):
        """ Monitor connection RSSI """
        while True:
            # Avoid RuntimeError if a connection is closed during iteration
            conn_list = list(self.connections.keys())
            for conn in conn_list:
                try:
                    address = self.connections[conn].address
                    _, rssi = self.lib.bt.connection.get_median_rssi(conn)
                    self.log.info(f"RSSI [{address}]: {rssi} dBm")
                    self.connections[conn].rssi = rssi
                    self._event_queue.put(ApEventConnectionRssi(self.id, conn, rssi))
                except bgapi.bglib.CommandFailedError:
                    pass
            time.sleep(RSSI_MEASUREMENT_PERIOD)

    # Common event callbacks
    def bt_evt_system_boot(self, evt):
        """ Bluetooth event callback """
        # Check if external bonding database feature is available on the target
        try:
            self.lib.bt.sm.get_bonding_handles(0)
            self.log.error("External bonding database feature missing from the target firmware.")
            self.stop()
            return
        except bgapi.bglib.CommandFailedError as err:
            # "feature not available" error is expected
            if err.errorcode != status.NOT_AVAILABLE:
                raise
        if self._identity_address is not None:
            if self._identity_type is None:
                self._identity_type = self.lib.bt.gap.ADDRESS_TYPE_PUBLIC_ADDRESS
            self.lib.bt.gap.set_identity_address(self._identity_address, self._identity_type)
        self.lib.bt.sm.set_bondable_mode(1)
        # Connection timing parameters are critical for the connection analyzer feature to work properly
        self.lib.bt.connection.set_default_parameters(
            CONNECTION_INTERVAL_MIN,
            CONNECTION_INTERVAL_MAX,
            0,     # latency
            100,   # timeout
            0,     # min_ce_length
            0xffff # max_ce_length
        )
        self.ready.set()

    def bt_evt_scanner_legacy_advertisement_report(self, evt):
        """ Bluetooth event callback """
        # Filter irrelevant events
        if not scan_filter(evt):
            return
        connectable = evt.event_flags & self.lib.bt.scanner.EVENT_FLAG_EVENT_FLAG_CONNECTABLE
        if not connectable:
            return
        self._event_queue.put(ApEventScanRssi(self.id,
                                              (evt.address, evt.address_type),
                                              evt.rssi))

    def bt_evt_connection_analyzer_report(self, evt):
        """ Bluetooth event callback """
        if evt.analyzer != self.analyzer:
            self.log.warning("Report event from unexpected analyzer: %d", evt.analyzer)
            return
        if evt.peripheral_rssi == self.lib.bt.connection.RSSI_CONST_RSSI_UNAVAILABLE:
            # No valid RSSI value available, drop this event
            return
        self._event_queue.put(ApEventAnalyzerRssi(self.id, evt.peripheral_rssi))

    def bt_evt_connection_analyzer_completed(self, evt):
        """ Bluetooth event callback """
        if evt.analyzer != self.analyzer:
            self.log.warning("Completed event from unexpected analyzer: %d", evt.analyzer)
            return
        self.log.warning("Analyzer stopped with reason %#x: '%s'", evt.reason, evt.reason)
        self.analyzer = None

    def bt_evt_connection_opened(self, evt):
        """ Bluetooth event callback """
        self.log.info(f"Connection opened: {evt.address}")
        self.connections[evt.connection].opened.set()
        self._event_queue.put(ApEventConnectionOpened())

    def bt_evt_connection_closed(self, evt):
        """ Bluetooth event callback """
        address = self.connections[evt.connection].address
        self.log.info(f"Connection to {address} closed with reason {evt.reason:#x}: '{evt.reason}'")
        del self.connections[evt.connection]
        if evt.reason == status.BT_CTRL_CONNECTION_TERMINATED_BY_LOCAL_HOST:
            self._event_queue.put(ApEventConnectionClosed())
        else:
            self._event_queue.put(ApEventConnectionLost())

    def bt_evt_connection_parameters(self, evt):
        """ Bluetooth event callback """
        if evt.security_mode != 0:
            # Successfully bonded
            # Start GATT discovery with finding the service handle
            self.lib.bt.gatt.discover_primary_services_by_uuid(
                evt.connection,
                GATT_SERVICE_UUID)

    def bt_evt_sm_bonding_failed(self, evt):
        """ Bluetooth event callback """
        address = self.connections[evt.connection].address
        self.log.error(f"Bonding with {address} failed with reason {evt.reason:#x}: '{evt.reason}'")

    # External bonding database event callbacks
    def bt_evt_external_bondingdb_data_request(self, evt):
        """ Bluetooth event callback """
        # 0 length data means that the data is not available in the external bonding DB
        data = self.connections[evt.connection].bonding_data.get(evt.type, b"")
        self.lib.bt.external_bondingdb.set_data(evt.connection, evt.type, data)

    def bt_evt_external_bondingdb_data(self, evt):
        """ Bluetooth event callback """
        self.connections[evt.connection].bonding_data[evt.type] = evt.data
        self._event_queue.put(ApEventBondingDbChanged())

    def bt_evt_external_bondingdb_data_ready(self, evt):
        """ Bluetooth event callback """
        # Initiate bonding
        self.lib.bt.sm.increase_security(evt.connection)

    # GATT event callbacks
    def bt_evt_gatt_service(self, evt):
        """ Bluetooth event callback """
        # Store service handle for the connection
        self.connections[evt.connection].service = evt.service

    def bt_evt_gatt_characteristic(self, evt):
        """ Bluetooth event callback """
        # Store characteristic handle for the connection
        self.connections[evt.connection].characteristic = evt.characteristic

    def bt_evt_gatt_procedure_completed(self, evt):
        """ Bluetooth event callback """
        if evt.result != status.OK:
            self.log.error(f"GATT procedure completed with status {evt.result:#x}: {evt.result}")
            return
        if self.connections[evt.connection].characteristic is None:
            # Continue GATT discovery with finding the characteristic handle
            self.lib.bt.gatt.discover_characteristics_by_uuid(
                evt.connection,
                self.connections[evt.connection].service,
                GATT_CHARACTERISTIC_UUID)
        elif not self.connections[evt.connection].notification:
            # Finally, request notification for the characteristic
            self.lib.bt.gatt.set_characteristic_notification(
                evt.connection,
                self.connections[evt.connection].characteristic,
                self.lib.bt.gatt.CLIENT_CONFIG_FLAG_NOTIFICATION)
            self.connections[evt.connection].notification = True

    def bt_evt_gatt_characteristic_value(self, evt):
        """ Bluetooth event callback """
        process_characteristic_value(self, evt)

class NetworkCoordinator:
    """ Roaming network coordinator managing multiple access points """
    def __init__(self, connectors, rssi_th=None, graph=False):
        if rssi_th is None:
            # Setting the lowest possible RSSI threshold is equivalent with disabling it
            self.rssi_threshold = -127
        else:
            self.rssi_threshold = rssi_th
        self._event_queue: "queue.Queue[ApEvent]" = queue.Queue()
        self._analyzer_queue: "queue.Queue[ApEventConnectionRssi]" = queue.Queue()
        self._ap_lock = threading.Lock() # prohibit parallel scanning and connection analysis
        self._start_scan = threading.Event()
        self._scan_result = dict[tuple, dict]()
        self._analyzer_result = {}
        self._bonding_db = load_bonding_db()
        self._bonding_db_dirty = threading.Event()
        self._graph_dirty = threading.Event()
        self.ap_dict = dict[int, AccessPoint]()
        for connector in connectors:
            ap = AccessPoint(connector,
                             self._event_queue,
                             identity_address=IDENTITY_ADDRESS)
            self.ap_dict[ap.id] = ap
        self.log = logging.getLogger(type(self).__name__)
        self._threads = list[threading.Thread]()
        self._threads.append(threading.Thread(target=self.event_handler, daemon=True))
        self._threads.append(threading.Thread(target=self.scan_scheduler, daemon=True))
        self._threads.append(threading.Thread(target=self.scan_task, daemon=True))
        self._threads.append(threading.Thread(target=self.analyzer_task, daemon=True))
        self._threads.append(threading.Thread(target=self.bonding_db_task, daemon=True))
        if graph:
            if shutil.which("dot") is None:
                self.log.error("Graphviz dot tool not found, continue without network graph viewer")
            else:
                self._graph_thread = threading.Thread(target=self.graph_task, daemon=True)

    def start(self):
        """ Start access points and wait for boot event, start worker threads """
        for ap in self.ap_dict.values():
            ap.start()
        for ap in self.ap_dict.values():
            if not ap.ready.wait(timeout=BOOT_TIMEOUT):
                raise RuntimeError(f"AP#{ap.id} failed to boot")
        self.log.info("all access points booted")
        for thread in self._threads:
            thread.start()
        if hasattr(self, "_graph_thread"):
            self._graph_thread.start()

    def stop(self):
        """ Stop all access points. """
        for ap in self.ap_dict.values():
            ap.stop()

    def scan(self):
        """ Scan for and connect to devices. """
        # clear scan result
        self._scan_result = dict[tuple, dict]()
        self.log.info("scan for devices...")
        # select only connectable access points for scanning
        scan_ap_list = [ap for ap in self.ap_dict.values() if ap.connectable]
        for ap in scan_ap_list:
            ap.start_scan()
        time.sleep(SCANNING_TIMEOUT)
        for ap in scan_ap_list:
            ap.stop_scan()
        if len(self._scan_result) == 0:
            self.log.info("no devices found")
            return
        self.log.info("device(s) found:")
        self.log.info(self._scan_result)
        # The following iteration walks through the discovered devices and picks the access point
        # with the best available RSSI to connect to. This algorithm disregards the fact that APs
        # have only a limited number of connections, and is therefore suboptimal under certain
        # conditions.
        for (address, address_type), result in self._scan_result.items():
            # remove non-connectable APs from the results
            result = {ap_id: rssi for ap_id, rssi in result.items() if self.ap_dict[ap_id].connectable}
            if len(result) == 0:
                self.log.warning("No AP available to connect to %s", address)
                continue
            # find access point with the largest RSSI value
            ap_id = max(result, key=result.get)
            self.ap_dict[ap_id].connect(
                address=address,
                address_type=address_type,
                bonding_data=self.get_bonding_data(address),
                rssi=result[ap_id])

    def update_scan_result(self, evt: ApEventScanRssi):
        """ Collect scan report events from various access points. """
        if evt.address not in self._scan_result.keys():
            self._scan_result[evt.address] = {}
        # New RSSI value overwrites old one (i.e. no averaging)
        self._scan_result[evt.address][evt.ap_id] = evt.rssi

    def analyze_connection(self, ap_id, conn_handle, rssi):
        """ Check if better RSSI is available for a connection """
        self._analyzer_result = {}
        conn_params = self.ap_dict[ap_id].get_conn_params(conn_handle)
        address = self.ap_dict[ap_id].connections[conn_handle].address
        address_type = self.ap_dict[ap_id].connections[conn_handle].address_type
        self.log.info("Start connection analysis for %s", address)
        ap_list = [ap for ap in self.ap_dict.values() if ap.connectable and ap.id != ap_id]
        if len(ap_list) == 0:
            self.log.info("No APs available for connection analysis")
            return
        for ap in ap_list:
            ap.start_analyzer(conn_params)
        time.sleep(CONNECTION_ANALYSIS_TIMEOUT)
        for ap in ap_list:
            ap.stop_analyzer()
        # Check if result dict is empty
        if not self._analyzer_result:
            self.log.info("No RSSI values available from connection analysis for %s", address)
            return
        # Actual connection RSSI is considered the best initially
        best_rssi = rssi
        best_ap = ap_id
        for _ap_id, _rssi in self._analyzer_result.items():
            if _rssi > best_rssi and _rssi > self.rssi_threshold:
                best_ap = _ap_id
                best_rssi = _rssi
        if best_ap == ap_id:
            self.log.info("%s - stay at AP#%d (%d dBm), no better AP available",
                          address, ap_id, rssi)
            return
        # Better AP available, close connection
        self.log.info("%s - switch from AP#%d (%d dBm) to AP#%d (%d dBm)",
                        address, ap_id, rssi, best_ap, best_rssi)
        self.ap_dict[ap_id].disconnect(conn_handle)
        # Connect to the best AP available
        self.ap_dict[best_ap].connect(
            address=address,
            address_type=address_type,
            bonding_data=self.get_bonding_data(address),
            # best_rssi might be a float value due to averaging, store it as integer
            rssi=int(best_rssi))

    def update_analyzer_result(self, evt: ApEventAnalyzerRssi):
        """ Collect RSSI measurements from various access points. """
        # RSSI values are expected only from one address
        if evt.ap_id not in self._analyzer_result.keys():
            # Store first RSSI value from AP
            self._analyzer_result[evt.ap_id] = evt.rssi
        else:
            # Upcoming RSSI values are averaged with the previous value
            self._analyzer_result[evt.ap_id] += evt.rssi
            self._analyzer_result[evt.ap_id] /= 2

    def get_bonding_data(self, address):
        """ Provide bonding data for connection """
        if address not in self._bonding_db.keys():
            # Create bonding data entry if not available yet
            self._bonding_db[address] = {}
        return self._bonding_db[address]

    def event_handler(self):
        """ Handle events from all access points """
        while True:
            try:
                # Timeout is needed for Windows hosts to get the KeyboardInterrupt in the main thread.
                evt = self._event_queue.get(timeout=0.5)
            except queue.Empty:
                continue
            if isinstance(evt, (ApEventConnectionRssi,
                                ApEventConnectionOpened,
                                ApEventConnectionClosed,
                                ApEventConnectionLost)):
                self._graph_dirty.set()

            if isinstance(evt, ApEventConnectionRssi):
                if evt.rssi < self.rssi_threshold:
                    # Trigger connection analysis
                    self._analyzer_queue.put(evt)
            elif isinstance(evt, ApEventAnalyzerRssi):
                self.update_analyzer_result(evt)
            elif isinstance(evt, ApEventScanRssi):
                self.update_scan_result(evt)
            elif isinstance(evt, ApEventConnectionLost):
                self._start_scan.set()
            elif isinstance(evt, ApEventBondingDbChanged):
                self._bonding_db_dirty.set()

    @property
    def graph(self):
        """ Graph representation of the network in dot language """
        dot = 'graph G {\n'
        # list access point nodes
        for ap_id in self.ap_dict.keys():
            dot += f'{ap_id} [label="AP {ap_id}"]\n'
        # represent device nodes with box shape
        dot += 'node [shape=box]\n'
        # list device nodes and edges
        for ap_id, ap in self.ap_dict.items():
            for conn_id, conn in ap.connections.items():
                # derive node name from access point ID and connection handle
                node = f"{ap_id}_{conn_id}"
                # unstable connections are marked with red color
                color = ', color="red"' if conn.rssi < self.rssi_threshold else ''
                dot += f'"{node}" [label="{conn.address}"{color}]\n'
                dot += f'{ap_id} -- "{node}" [label="{conn.rssi} dBm"{color}]\n'
        dot += '}\n'
        return dot

    def scan_scheduler(self):
        """ Schedule scan procedure periodically """
        while True:
            self._start_scan.set()
            time.sleep(SCANNING_PERIOD)

    def scan_task(self):
        """ Check for scan request """
        while True:
            self._start_scan.wait()
            with self._ap_lock:
                self.scan()
            self._start_scan.clear()

    def analyzer_task(self):
        """ Check for connection analysis request """
        while True:
            try:
                # Timeout is needed for Windows hosts to get the KeyboardInterrupt in the main thread.
                evt = self._analyzer_queue.get(timeout=0.5)
            except queue.Empty:
                continue
            # Check if connection still exists
            if evt.connection in self.ap_dict[evt.ap_id].connections.keys():
                with self._ap_lock:
                    self.analyze_connection(evt.ap_id, evt.connection, evt.rssi)

    def bonding_db_task(self):
        """ Check if bonding database needs to be saved """
        while True:
            self._bonding_db_dirty.wait()
            # Delay writing to file to avoid too frequent file access
            time.sleep(1)
            self._bonding_db_dirty.clear()
            save_bonding_db(self._bonding_db)

    def graph_task(self):
        """ Check if graph viewer needs to be updated """
        while True:
            self._graph_dirty.wait()
            # Limit rendering frequency by adding delay
            time.sleep(1)
            self._graph_dirty.clear()
            view(self.graph)

def scan_filter(evt):
    """ Filter for selecting devices of interest """
    return find_service_in_advertisement(evt.data, GATT_SERVICE_UUID)

def process_characteristic_value(ap: AccessPoint, evt):
    """ Process characteristic notification events """
    value = int(evt.value[1])
    address = ap.connections[evt.connection].address
    ap.log.info(f"heart rate [{address}]: {value} BPM")

def load_bonding_db():
    """ Load bonding database from file """
    if not os.path.exists(BONDING_DB_PATH):
        return {}
    with open(BONDING_DB_PATH, "r", encoding="utf-8") as bonding_db:
        db_in = json.load(bonding_db)
        db_out = {}
        for address, data in db_in.items():
            db_out[address] = {int(key): bytes.fromhex(value) for key, value in data.items()}
        return db_out

def save_bonding_db(db_in: dict):
    """ Save bonding database to file """
    db_out = {}
    for address, data in db_in.items():
        db_out[address] = {key: value.hex() for key, value in data.items()}
    with open(BONDING_DB_PATH, "w", encoding="utf-8") as bonding_db:
        json.dump(db_out, bonding_db)

def view(dot: str):
    """ View network graph """
    result = subprocess.run(["dot", "-Tsvg"], text=True, input=dot, stdout=subprocess.PIPE, check=True)
    with open(NETWORK_GRAPH_PATH, "w", encoding="utf-8") as svg:
        svg.write(result.stdout)
    webbrowser.open(pathlib.Path(NETWORK_GRAPH_PATH).absolute().as_uri())

def main():
    """ Main function. """
    parser = ArgumentParser(description=__doc__, single_mode=False)
    parser.add_argument(
        "-d",
        action="store_true",
        help="Delete bondings")
    parser.add_argument(
        "-g",
        action="store_true",
        help="View network graph")
    parser.add_argument(
        "-r", "--rssi",
        type=int,
        help="Connection is closed if connection RSSI value drops below this value in dBm and if an AP with potentially better RSSI is available.")
    args = parser.parse_args()
    connectors = get_connector(args)
    if args.d:
        # Delete bonding database file if exists
        try:
            os.remove(BONDING_DB_PATH)
        except FileNotFoundError:
            pass
    app = NetworkCoordinator(connectors, rssi_th=args.rssi, graph=args.g)
    app.start()
    # Catch KeyboardInterrupt
    try:
        while True:
            time.sleep(60)
    except KeyboardInterrupt:
        app.stop()

# Script entry point.
if __name__ == "__main__":
    main()
```
