## Supplementary material for "VesperFlow: An automated pipeline for decoding multisensor data from biologging tags": peripheral.py

```
#!/usr/bin/env python3
""" Roaming NCP-host Example Application for peripheral devices.
"""

# Copyright 2023 Silicon Laboratories Inc. www.silabs.com
#
# SPDX-License-Identifier: Zlib
#
# The licensor of this software is Silicon Laboratories Inc.
#
# This software is provided 'as-is', without any express or implied
# warranty. In no event will the authors be held liable for any damages
# arising from the use of this software.
#
# Permission is granted to anyone to use this software for any purpose,
# including commercial applications, and to alter it and redistribute it
# freely, subject to the following restrictions:
#
# 1. The origin of this software must not be misrepresented; you must not
#    claim that you wrote the original software. If you use this software
#    in a product, an acknowledgment in the product documentation would be
#    appreciated but is not required.
# 2. Altered source versions must be plainly marked as such, and must not be
#    misrepresented as being the original software.
# 3. This notice may not be removed or altered from any source distribution.

import os.path
import sys
import threading
import time
import bgapi

sys.path.append(os.path.join(os.path.dirname(__file__), "../.."))
from common.util import ArgumentParser, BluetoothApp, get_connector
import common.status as status

# Heart rate measurement characteristic notification period in seconds
NOTIFICATION_PERIOD = 10.0
# Characteristic values
GATTDB_DEVICE_NAME = b"Heart Rate Sensor"
GATTDB_MANUFACTURER_NAME_STRING = b"Silicon Labs"

class HeartRateSensor(BluetoothApp):
    """ Heart Rate Sensor derived from generic BluetoothApp. """
    def __init__(self, connector, delete_bondings=False, **kwargs):
        self.delete_bondings = delete_bondings
        self.adv_handle = None
        self.gattdb_heart_rate_measurement = None
        self.notification_event = threading.Event()
        self.notification_thread = threading.Thread(target=self.notification_task, daemon=True)
        self.notification_thread.start()
        super().__init__(connector, **kwargs)
        # Dummy heart rate data derived from instance ID
        self.heart_rate = self.id + 100

    def bt_evt_system_boot(self, evt):
        """ Bluetooth event callback """
        # Check if external bonding database feature is available on the target
        try:
            self.lib.bt.sm.get_bonding_handles(0)
        except bgapi.bglib.CommandFailedError as err:
            if err.errorcode == status.NOT_AVAILABLE:
                self.log.error("External bonding database feature present in the target firmware.")
            raise
        self.notification_event.clear()
        self.adv_handle = None
        self.gattdb_init()
        if self.delete_bondings:
            # Delete bondings on first boot only.
            self.delete_bondings = False
            self.lib.bt.sm.delete_bondings()
        self.lib.bt.sm.set_bondable_mode(1)
        self.adv_start()

    def bt_evt_connection_opened(self, evt):
        """ Bluetooth event callback """
        self.log.info("Connection opened: %s", evt.address)

    def bt_evt_connection_closed(self, evt):
        """ Bluetooth event callback """
        self.notification_event.clear()
        self.log.info(f"Connection closed with reason {evt.reason:#x}: '{evt.reason}'")
        self.adv_start()

    def bt_evt_connection_parameters(self, evt):
        """ Bluetooth event callback """
        if evt.security_mode != 0:
            self.log.info("Bonded")

    def bt_evt_sm_bonding_failed(self, evt):
        """ Bluetooth event callback """
        self.log.error(f"Bonding failed with reason {evt.reason:#x}: '{evt.reason}'")

    def bt_evt_gatt_server_characteristic_status(self, evt):
        """ Bluetooth event callback """
        if evt.characteristic == self.gattdb_heart_rate_measurement:
            if evt.status_flags == self.lib.bt.gatt_server.CHARACTERISTIC_STATUS_FLAG_CLIENT_CONFIG:
                # The remote client requested the status change.
                if evt.client_config_flags == self.lib.bt.gatt_server.CLIENT_CONFIGURATION_DISABLE:
                    self.log.info("Notification disabled.")
                    self.notification_event.clear()
                elif not self.notification_event.is_set():
                    self.log.info("Notification enabled.")
                    self.notification_event.set()

    def gattdb_init(self):
        """ Initialize GATT database. """
        _, session = self.lib.bt.gattdb.new_session()

        # Generic Access
        _, service = self.lib.bt.gattdb.add_service(
            session,
            self.lib.bt.gattdb.SERVICE_TYPE_PRIMARY_SERVICE,
            0,
            b"\x00\x18"
        )
        # Device Name
        self.lib.bt.gattdb.add_uuid16_characteristic(
            session,
            service,
            self.lib.bt.gattdb.CHARACTERISTIC_PROPERTIES_CHARACTERISTIC_READ |
            self.lib.bt.gattdb.CHARACTERISTIC_PROPERTIES_CHARACTERISTIC_WRITE,
            0,
            0,
            b"\x00\x2A",
            self.lib.bt.gattdb.VALUE_TYPE_FIXED_LENGTH_VALUE,
            len(GATTDB_DEVICE_NAME),
            GATTDB_DEVICE_NAME
        )
        # Appearance
        self.lib.bt.gattdb.add_uuid16_characteristic(
            session,
            service,
            self.lib.bt.gattdb.CHARACTERISTIC_PROPERTIES_CHARACTERISTIC_READ,
            0,
            0,
            b"\x01\x2A",
            self.lib.bt.gattdb.VALUE_TYPE_FIXED_LENGTH_VALUE,
            2,
            b"\x00\x00"
        )
        self.lib.bt.gattdb.start_service(session, service)

        # Device Information
        _, service = self.lib.bt.gattdb.add_service(
            session,
            self.lib.bt.gattdb.SERVICE_TYPE_PRIMARY_SERVICE,
            0,
            b"\x0A\x18"
        )
        # Manufacturer Name String
        self.lib.bt.gattdb.add_uuid16_characteristic(
            session,
            service,
            self.lib.bt.gattdb.CHARACTERISTIC_PROPERTIES_CHARACTERISTIC_READ,
            0,
            0,
            b"\x29\x2A",
            self.lib.bt.gattdb.VALUE_TYPE_FIXED_LENGTH_VALUE,
            len(GATTDB_MANUFACTURER_NAME_STRING),
            GATTDB_MANUFACTURER_NAME_STRING
        )
        # System ID
        # Pad and reverse unique ID to get System ID.
        addr = self.address.split(":")
        system_id = bytes.fromhex("".join(addr[:3] + ["ff", "fe"] + addr[3:]))
        self.lib.bt.gattdb.add_uuid16_characteristic(
            session,
            service,
            self.lib.bt.gattdb.CHARACTERISTIC_PROPERTIES_CHARACTERISTIC_READ,
            0,
            0,
            b"\x23\x2A",
            self.lib.bt.gattdb.VALUE_TYPE_FIXED_LENGTH_VALUE,
            8,
            system_id
        )
        self.lib.bt.gattdb.start_service(session, service)

        # Heart Rate Service
        _, service = self.lib.bt.gattdb.add_service(
            session,
            self.lib.bt.gattdb.SERVICE_TYPE_PRIMARY_SERVICE,
            self.lib.bt.gattdb.SERVICE_PROPERTY_FLAGS_ADVERTISED_SERVICE,
            b"\x0D\x18"
        )
        # Heart Rate Measurement
        _, self.gattdb_heart_rate_measurement = self.lib.bt.gattdb.add_uuid16_characteristic(
            session,
            service,
            self.lib.bt.gattdb.CHARACTERISTIC_PROPERTIES_CHARACTERISTIC_NOTIFY,
            self.lib.bt.gattdb.SECURITY_REQUIREMENTS_BONDED_NOTIFY,
            0,
            b"\x37\x2A",
            self.lib.bt.gattdb.VALUE_TYPE_FIXED_LENGTH_VALUE,
            2,
            b"\x00"*2
        )
        self.lib.bt.gattdb.start_service(session, service)

        self.lib.bt.gattdb.commit(session)

    def adv_start(self):
        """ Start advertising. """
        if self.adv_handle is None:
            # Create advertising set for the first call.
            _, self.adv_handle = self.lib.bt.advertiser.create_set()
            self.lib.bt.legacy_advertiser.generate_data(
                self.adv_handle,
                self.lib.bt.advertiser.DISCOVERY_MODE_GENERAL_DISCOVERABLE)
            # Set advertising interval to 100 ms.
            self.lib.bt.advertiser.set_timing(
                self.adv_handle,
                160,  # interval min
                160,  # interval max
                0,    # duration
                0)    # max events
        self.lib.bt.legacy_advertiser.start(
            self.adv_handle,
            self.lib.bt.legacy_advertiser.CONNECTION_MODE_CONNECTABLE)

    def send_notification(self):
        """ Send notification with dummy data. """
        self.log.info("Sending %d BPM", self.heart_rate)
        # The characteristic value consists of 1 byte Flags Field and 1 byte Measurement Value Field
        value = bytes([0, self.heart_rate])
        self.lib.bt.gatt_server.notify_all(self.gattdb_heart_rate_measurement, value)

    def notification_task(self):
        """ Indication task executed in its own thread. """
        while True:
            self.notification_event.wait()
            try:
                self.send_notification()
            except bgapi.bglib.CommandError as err:
                # Tolerate command error, e.g. if the device resets
                self.log.error(err)
                self.notification_event.clear()
            time.sleep(NOTIFICATION_PERIOD)

def main():
    """ Main function. """
    parser = ArgumentParser(description=__doc__, single_mode=False)
    parser.add_argument(
        "-d",
        action="store_true",
        help="Delete bondings")
    args = parser.parse_args()
    connector = get_connector(args)
    # Instantiate an application for every connector.
    apps = [HeartRateSensor(conn, args.d) for conn in connector]
    for app in apps:
        app.start()
    # Catch KeyboardInterrupt
    try:
        while True:
            time.sleep(60)
    except KeyboardInterrupt:
        for app in apps:
            app.stop()

# Script entry point.
if __name__ == "__main__":
    main()
```
