## Supplementary material for "VesperFlow: An automated pipeline for decoding multisensor data from biologging tags": README.md

# Bluetooth - Roaming

This pair of examples models a medical application. The goal is to gather data from several BLE
heart rate (HR) sensors in a large facility using multiple access points (AP). Whenever a person
with the HR sensor moves out from the radio range of an AP and loses connection, a nearby AP shall
find it and continue the HR data collection. The access points are supervised by a network
coordinator.

![Roaming Network](image/roaming_network.png)

From BLE connection point of vew, the APs are the central devices, the HR sensors are the peripheral
devices. The example is scalable, i.e., both `central.py` and `peripheral.py` can handle any number
of NCP radio boards.

## central.py

Each radio board controlled by the `central.py` forms a physical access point. The APs are
identified by a sequence number starting from 0. The access points are managed by a single network
coordinator. Scanning for heart rate sensors is performed for 3 seconds (`SCANNING_TIMEOUT`) in
every 30 seconds (`SCANNING_PERIOD`) or immediately when a device disconnects. When a heart rate
sensor is found, the nearest AP (i.e. the one with the largest RSSI) connects to it. Every AP uses
the same identity (`IDENTITY_ADDRESS`). I.e., all heart rate sensors seem to be connected to the
same virtual access point.

### RSSI measurement

The RSSI of the active connections is measured every 10 seconds (`RSSI_MEASUREMENT_PERIOD`). If the
RSSI drops below a threshold value, the other APs start to analyze this connection, i.e., they
measure the RSSI of the connection for 3 seconds (`CONNECTION_ANALYSIS_TIMEOUT`). If there is an AP
(e.g. AP4) that measures the peripheral side of the connection with a better RSSI than the AP in the
active connection (e.g. AP1), then the connection is closed and AP4 initiates a new connection to
the HR sensor. The RSSI threshold that triggers the connection analysis procedure depends on the
actual network, and is therefore a matter of experimenting. The RSSI threshold is adjustable using
the `-r` command line switch. When this switch is omitted, the connection analysis feature is
disabled.

### Network visualization

Following the network topology can be difficult already at low number of nodes. A simple graphical
tool has been added to help visualize the network. When enabled with the `-g` command line switch, a
`graph.svg` image is drawn whenever there is a change in the network (e.g. new HR sensor found, RSSI
value updated) and the image is opened in the web browser. The feature requires the
[Graphviz](https://graphviz.org/download/) package to be installed on your environment. Make sure
that the `dot` executable (which is part of the Graphviz package) is available on the system path.

<img src="image/graphviz_installer.png" alt="Graphviz Installer" width="500"/>

### Bonding

Reading the HR characteristic of the peripheral devices requires bonding. Bonding is performed by
the access points using external bonding database feature. This means that the user application is
responsible for storing the bonding keys provided by the Bluetooth stack and provide those keys when
requested by the stack. In this example, bonding data is stored in a json file between sessions.

During development, bonding keys can go out of sync. The bonding keys can be deleted either by using
the `-d` command line switch or by removing the `bonding_db.json` file. Bonding keys must be deleted
in the peripheral devices too.

Bonding keys are sensitive data therefore both the bonding data storage file and the NCP
communication between the host SW and the target FW shall be encrypted when developing products that
use the external bonding database feature.

### NCP target features

Neither the external bonding database nor the connection analyzer Bluetooth feature is part of the
**Bluetooth - NCP** example by default, these have to be added manually when creating the project
for the access points. Their SLC component IDs are `bluetooth_feature_external_bonding_database` and
`bluetooth_feature_connection_analyzer`. Heart rate sensor devices shall be used with the default
**Bluetooth - NCP** example without the additional features.


## peripheral.py

Each radio board controlled by the `peripheral.py` forms a HR sensor with limited functionalities
according to the [Heart Rate Service 1.0 specification](https://www.bluetooth.org/docman/handlers/downloaddoc.ashx?doc_id=239866).
The HR sensors are identified by a sequence number starting from 0. On boot event, the HR sensors
start advertising. The advertising stops when connected to an AP. The HR measurement value is a
unique constant dummy value, derived from the ID of the HR sensor.

### Bonding

Accessing the HR characteristic requires bonding. The bonding keys are stored by the BLE stack in
the NVM of the target device. During development, these bonding keys can go out of sync. The bonding
keys can be deleted by using the `-d` command line switch. Bonding keys must be deleted in the
central devices too.
