## Supplementary material for "VesperFlow: An automated pipeline for decoding multisensor data from biologging tags": README.md

# Bluetooth - Thermometer

Original C example: **Bluetooth - SoC Thermometer**

This example implements the [Health Thermometer Service](https://www.bluetooth.com/specifications/specs/health-thermometer-service-1-0/).
It enables a peer device to connect and receive temperature values via Bluetooth.

The NCP Host has no access to the board peripherals, only to the Bluetooth stack via BGAPI.
Therefore, neither the button nor the temperature sensor is available in this example. The
temperature measurement values are just dummy values for demo purposes.
