## Supplementary material for "VesperFlow: An automated pipeline for decoding multisensor data from biologging tags": app.py

```
#!/usr/bin/env python3
"""
Thermometer Client NCP-host Example Application.
"""

# Copyright 2021 Silicon Laboratories Inc. www.silabs.com
#
# SPDX-License-Identifier: Zlib
#
# The licensor of this software is Silicon Laboratories Inc.
#
# This software is provided 'as-is', without any express or implied
# warranty. In no event will the authors be held liable for any damages
# arising from the use of this software.
#
# Permission is granted to anyone to use this software for any purpose,
# including commercial applications, and to alter it and redistribute it
# freely, subject to the following restrictions:
#
# 1. The origin of this software must not be misrepresented; you must not
#    claim that you wrote the original software. If you use this software
#    in a product, an acknowledgment in the product documentation would be
#    appreciated but is not required.
# 2. Altered source versions must be plainly marked as such, and must not be
#    misrepresented as being the original software.
# 3. This notice may not be removed or altered from any source distribution.

from dataclasses import dataclass
import os.path
import sys

sys.path.append(os.path.join(os.path.dirname(__file__), "../.."))
from common.conversion import Ieee11073Float
from common.util import ArgumentParser, BluetoothApp, get_connector, find_service_in_advertisement
import common.status as status

# Constants
HEALTH_THERMOMETER_SERVICE = b"\x09\x18"
TEMPERATURE_MEASUREMENT_CHAR = b"\x1c\x2a"

CONN_INTERVAL_MIN = 80   # 100 ms
CONN_INTERVAL_MAX = 80   # 100 ms
CONN_SLAVE_LATENCY = 0   # no latency
CONN_TIMEOUT = 100       # 1000 ms
CONN_MIN_CE_LENGTH = 0
CONN_MAX_CE_LENGTH = 65535

SCAN_INTERVAL = 16       # 10 ms
SCAN_WINDOW = 16         # 10 ms
SCAN_PASSIVE = 0

# The maximum number of connections has to match with the configuration on the target side.
SL_BT_CONFIG_MAX_CONNECTIONS = 4

@dataclass
class Connection:
    """ Connection representation """
    address: str
    address_type: int
    service: int=None
    characteristic: int=None

class App(BluetoothApp):
    """ Application derived from generic BluetoothApp. """
    def bt_evt_system_boot(self, evt):
        """ Bluetooth event callback

        This event indicates that the device has started and the radio is ready.
        Do not call any stack command before receiving this boot event!
        """
        # Set the default connection parameters for subsequent connections
        self.lib.bt.connection.set_default_parameters(
            CONN_INTERVAL_MIN,
            CONN_INTERVAL_MAX,
            CONN_SLAVE_LATENCY,
            CONN_TIMEOUT,
            CONN_MIN_CE_LENGTH,
            CONN_MAX_CE_LENGTH)
        # Start scanning - looking for thermometer devices
        self.lib.bt.scanner.start(
            self.lib.bt.scanner.SCAN_PHY_SCAN_PHY_1M,
            self.lib.bt.scanner.DISCOVER_MODE_DISCOVER_GENERIC)
        self.log.info("Scanning started...")
        self.conn_state = "scanning"
        self.connections = dict[int, Connection]()

    def bt_evt_scanner_legacy_advertisement_report(self, evt):
        """ Bluetooth event callback """
        # Parse advertisement packets
        if (evt.event_flags & self.lib.bt.scanner.EVENT_FLAG_EVENT_FLAG_CONNECTABLE and
            evt.event_flags & self.lib.bt.scanner.EVENT_FLAG_EVENT_FLAG_SCANNABLE):
            # If a thermometer advertisement is found...
            if find_service_in_advertisement(evt.data, HEALTH_THERMOMETER_SERVICE):
                # then stop scanning for a while
                self.lib.bt.scanner.stop()
                # and connect to that device
                if len(self.connections) < SL_BT_CONFIG_MAX_CONNECTIONS:
                    self.lib.bt.connection.open(
                        evt.address,
                        evt.address_type,
                        self.lib.bt.gap.PHY_PHY_1M)
                    self.conn_state = "opening"

    def bt_evt_connection_opened(self, evt):
        """ Bluetooth event callback """
        self.log.info(f"Connection opened to {evt.address}")
        self.connections[evt.connection] = Connection(evt.address, evt.address_type)
        # Discover Health Thermometer service on the slave device
        self.lib.bt.gatt.discover_primary_services_by_uuid(
            evt.connection,
            HEALTH_THERMOMETER_SERVICE)
        self.conn_state = "discover_services"

    def bt_evt_gatt_service(self, evt):
        """ Bluetooth event callback """
        self.connections[evt.connection].service = evt.service

    def bt_evt_gatt_characteristic(self, evt):
        """ Bluetooth event callback """
        self.connections[evt.connection].characteristic = evt.characteristic

    def bt_evt_gatt_procedure_completed(self, evt):
        """ Bluetooth event callback """
        if evt.result != status.OK:
            address = self.connections[evt.connection].address
            self.log.error(f"GATT procedure for {address} completed with status {evt.result:#x}: {evt.result}")
            return
        # If service discovery finished
        if self.conn_state == "discover_services":
            # Discover thermometer characteristic on the slave device
            self.lib.bt.gatt.discover_characteristics_by_uuid(
                evt.connection,
                self.connections[evt.connection].service,
                TEMPERATURE_MEASUREMENT_CHAR)
            self.conn_state = "discover_characteristics"

        # If characteristic discovery finished
        elif self.conn_state == "discover_characteristics":
            # enable indications
            self.lib.bt.gatt.set_characteristic_notification(
                evt.connection,
                self.connections[evt.connection].characteristic,
                self.lib.bt.gatt.CLIENT_CONFIG_FLAG_INDICATION)
            self.conn_state = "enable_indication"

        # If indication enable process finished
        elif self.conn_state == "enable_indication":
            # and further connections are possible
            if len(self.connections) < SL_BT_CONFIG_MAX_CONNECTIONS:
                # start scanning again to find new devices
                self.lib.bt.scanner.start(
                    self.lib.bt.scanner.SCAN_PHY_SCAN_PHY_1M,
                    self.lib.bt.scanner.DISCOVER_MODE_DISCOVER_GENERIC)
                self.conn_state = "scanning"
            else:
                self.conn_state = "running"

    def bt_evt_connection_closed(self, evt):
        """ Bluetooth event callback """
        address = self.connections[evt.connection].address
        self.log.info(f"Connection to {address} closed with reason {evt.reason:#x}: '{evt.reason}'")
        del self.connections[evt.connection]
        if self.conn_state != "scanning":
            # start scanning again to find new devices
            self.lib.bt.scanner.start(
                self.lib.bt.scanner.SCAN_PHY_SCAN_PHY_1M,
                self.lib.bt.scanner.DISCOVER_MODE_DISCOVER_GENERIC)
            self.conn_state = "scanning"

    def bt_evt_gatt_characteristic_value(self, evt):
        """ Bluetooth event callback """
        address = self.connections[evt.connection].address
        temperature = Ieee11073Float.from_bytes(evt.value[1:])
        # The first byte of the characteristic value is the flags field,
        # the first bit in the flags field encodes the temperature unit.
        if evt.value[0] & 1:
            unit = "F"
        else:
            unit = "C"
        # Send confirmation for the indication
        self.lib.bt.gatt.send_characteristic_confirmation(evt.connection)
        # Get the RSSI of the connection
        _, rssi = self.lib.bt.connection.get_median_rssi(evt.connection)
        # Print the values
        print(f"{address} [{rssi:4} dBm] {temperature:6.6} {unit}")

# Script entry point.
if __name__ == "__main__":
    parser = ArgumentParser(description=__doc__)
    args = parser.parse_args()
    connector = get_connector(args)
    # Instantiate the application.
    app = App(connector)
    # Running the application blocks execution until it terminates.
    app.run()
```
