## Supplementary material for "VesperFlow: An automated pipeline for decoding multisensor data from biologging tags": README.md

# Bluetooth - Thermometer Client

Original C example: **Bluetooth - SoC Thermometer Client**

This example application demonstrates the operation of a central device in a multi-peripheral BLE
topology. The Silicon Labs Bluetooth stack supports simultaneous connection for up to eight peripheral
devices at a time. This sample application illustrates how to handle simultaneous connection
for up to four thermometer peripheral devices.
