## Supplementary material for "VesperFlow: An automated pipeline for decoding multisensor data from biologging tags": app.py

```
#!/usr/bin/env python3
"""
BtMesh Empty NCP-host Example Application.
"""

# Copyright 2021 Silicon Laboratories Inc. www.silabs.com
#
# SPDX-License-Identifier: Zlib
#
# The licensor of this software is Silicon Laboratories Inc.
#
# This software is provided 'as-is', without any express or implied
# warranty. In no event will the authors be held liable for any damages
# arising from the use of this software.
#
# Permission is granted to anyone to use this software for any purpose,
# including commercial applications, and to alter it and redistribute it
# freely, subject to the following restrictions:
#
# 1. The origin of this software must not be misrepresented; you must not
#    claim that you wrote the original software. If you use this software
#    in a product, an acknowledgment in the product documentation would be
#    appreciated but is not required.
# 2. Altered source versions must be plainly marked as such, and must not be
#    misrepresented as being the original software.
# 3. This notice may not be removed or altered from any source distribution.

import os.path
import sys

sys.path.append(os.path.join(os.path.dirname(__file__), "../.."))
from common.util import ArgumentParser, BtMeshApp, get_connector

# Advertising options
PB_ADV = 0x1
PB_GATT = 0x2

class App(BtMeshApp):
    """ Application derived from BtMeshApp. """
        # This event indicates the device has started and the radio is ready.
        # Do not call any stack command before receiving this boot event!
    def btmesh_evt_node_initialized(self, evt):
        """ Bluetooth mesh event callback """
        if not evt.provisioned:
            self.lib.btmesh.node.start_unprov_beaconing(PB_ADV | PB_GATT)

        ####################################
        # Add further event handlers here. #
        ####################################

# Script entry point.
if __name__ == "__main__":
    parser = ArgumentParser(description=__doc__)
    args = parser.parse_args()
    connector = get_connector(args)
    # Instantiate and run application.
    app = App(connector)
    app.run()
```
