## Supplementary material for "VesperFlow: An automated pipeline for decoding multisensor data from biologging tags": README.md

# Bluetooth mesh - Empty

Original C example: **Bluetooth Mesh - SoC Empty**

This example demonstrates the bare minimum needed for a Bluetooth Mesh Network Co-Processor
application. The application establishes connection with the target device according to the
parameters provided at application launch, starts Unprovisioned Device Beaconing after boot
and can be provisioned to a Mesh Network.
This example can be used as a starting point for a python host project and can be
extended according to application requirements, utilizing the whole set of pyBGAPI events
and commands.
