## Supplementary material for "VesperFlow: An automated pipeline for decoding multisensor data from biologging tags": app.py

```
#!/usr/bin/env python3
"""
BtMesh NCP Light Server Example Application.
"""

# Copyright 2022 Silicon Laboratories Inc. www.silabs.com
#
# SPDX-License-Identifier: Zlib
#
# The licensor of this software is Silicon Laboratories Inc.
#
# This software is provided 'as-is', without any express or implied
# warranty. In no event will the authors be held liable for any damages
# arising from the use of this software.
#
# Permission is granted to anyone to use this software for any purpose,
# including commercial applications, and to alter it and redistribute it
# freely, subject to the following restrictions:
#
# 1. The origin of this software must not be misrepresented; you must not
#    claim that you wrote the original software. If you use this software
#    in a product, an acknowledgment in the product documentation would be
#    appreciated but is not required.
# 2. Altered source versions must be plainly marked as such, and must not be
#    misrepresented as being the original software.
# 3. This notice may not be removed or altered from any source distribution.

import os.path
import sys
import threading
import struct

from dataclasses import dataclass, fields
from lighting_server_gui import lighting_server_gui_start
from ctl_server import CTLServer
from onoff_server import OnOffServer
from lightness_server import LightnessServer
from factory_reset import FactoryReset
from bgapi.bglib import CommandFailedError
sys.path.append(os.path.join(os.path.dirname(__file__), "../.."))
from common.util import ArgumentParser, get_connector
import common.status as status
import common.btmesh_models as model

IMMEDIATE = 0

# Advertising options
PB_ADV = 0x1
PB_GATT = 0x2
class App(CTLServer, FactoryReset):
    """ Application derived from BtMeshApp. """

    @dataclass
    class NVMState:
        """ Dataclass for NVM states. """
        lighting_state_nvm = None
        ctl_state_nvm = None

    # Common event callbacks
    def btmesh_evt_node_initialized(self, evt):
        """ Bluetooth mesh event callback """
        if evt.provisioned:
            self.log.info("Node initialized and provisioned")

        else:
            self.lib.btmesh.node.start_unprov_beaconing(PB_ADV | PB_GATT)
            self.log.info("Node is ready to be provisioned")

        self.lib.btmesh.generic_server.init()
        self.log.info("All generic server models initialized")
        # Scene model set up and init
        self.lib.btmesh.scene_server.init(0)
        self.lib.btmesh.scene_setup_server.init(0)
        # Initialize Friend functionality
        self.lib.btmesh.friend.init()
        self.log.info("Friend feature activated")
        self.restore_nvm_last_sate()

    def btmesh_evt_friend_friendship_established(self, _evt):
        """ Bluetooth mesh event callback """
        self.log.info("BT mesh Friendship established with LPN")

    def btmesh_evt_friend_friendship_terminated(self, _evt):
        """ Bluetooth mesh event callback """
        self.log.info("BT mesh Friendship terminated with LPN")

    def btmesh_evt_scene_server_recall(self, evt):
        """ Bluetooth mesh event callback """
        self.log.info(f"----- Recall scene {evt.selected_scene} -----")

    def btmesh_evt_scene_setup_server_store(self, evt):
        """ Bluetooth mesh event callback """
        self.log.info(f"----- Store to scene {evt.scene_id} -----")

    def btmesh_evt_generic_server_client_request(self, evt):
        """ Bluetooth mesh event callback """
        if (evt.type == self.lib.btmesh.generic_client.SET_REQUEST_TYPE_REQUEST_ON_OFF
            and evt.elem_index == 0):
            self.log.info("----- OnOff request -----")
            OnOffServer.onoff_request(self,evt)
        elif (evt.type == self.lib.btmesh.generic_client.SET_REQUEST_TYPE_REQUEST_LIGHTNESS_ACTUAL
              and evt.elem_index == 0):
            self.log.info("----- Lightness request -----")
            LightnessServer.lightness_request(self, evt)
        elif (evt.type == self.lib.btmesh.generic_client.SET_REQUEST_TYPE_REQUEST_LIGHTNESS_LINEAR
            and evt.elem_index == 0):
            self.log.info("----- Lightness request -----")
            LightnessServer.lightness_request(self, evt)
        elif (evt.type == self.lib.btmesh.generic_client.SET_REQUEST_TYPE_REQUEST_CTL
            and evt.elem_index == 0):
            self.log.info("----- CTL request -----")
            CTLServer.ctl_request(self, evt)
        elif (evt.type== self.lib.btmesh.generic_client.SET_REQUEST_TYPE_REQUEST_LEVEL_MOVE
            and evt.elem_index == 0):
            self.log.info("----- Level move -----")
            self.log.debug(evt)
            LightnessServer.pri_level_request(self, evt)
        elif (evt.type == self.lib.btmesh.generic_client.SET_REQUEST_TYPE_REQUEST_LEVEL
            and evt.elem_index == 0):
            self.log.info("----- Generic level -----")
            LightnessServer.pri_level_request(self, evt)
        elif (evt.type == self.lib.btmesh.generic_client.SET_REQUEST_TYPE_REQUEST_LEVEL_DELTA
            and evt.elem_index == 0):
            self.log.info("----- Level delta -----")
            LightnessServer.pri_level_request(self, evt)
        elif (evt.type== self.lib.btmesh.generic_client.SET_REQUEST_TYPE_REQUEST_LEVEL_HALT
            and evt.elem_index == 0):
            self.log.info("----- Level halt -----")
            LightnessServer.pri_level_request(self, evt)

    def btmesh_evt_generic_server_state_recall(self, evt):
        """ Bluetooth mesh event callback """
        # Generic OnOff Server
        if (evt.model_id == model.BTMESH_GENERIC_ON_OFF_SERVER_MODEL_ID
            and evt.elem_index == 0):
            self.onoff_recall(evt)
        # Generic Level Server
        elif (evt.model_id == model.BTMESH_GENERIC_LEVEL_SERVER_MODEL_ID
            and evt.elem_index == 0):
            self.pri_level_recall(evt)
        # Light Lightness Server
        elif (evt.model_id == model.BTMESH_LIGHTING_LIGHTNESS_SERVER_MODEL_ID
            and evt.elem_index == 0):
            self.lightness_recall(evt)
        # Light CTL Server
        elif (evt.model_id == model.BTMESH_LIGHTING_CTL_SERVER_MODEL_ID
            and evt.elem_index == 0):
            self.ctl_recall(evt)

    def btmesh_evt_generic_server_state_changed(self, evt):
        """ Bluetooth mesh event callback """
        if (evt.model_id == model.BTMESH_GENERIC_ON_OFF_SERVER_MODEL_ID
            and evt.elem_index == 0):
            self.onoff_change(evt)
        elif (evt.model_id == model.BTMESH_GENERIC_LEVEL_SERVER_MODEL_ID
            and evt.elem_index == 0):
            self.pri_level_change(evt)
        elif (evt.model_id == model.BTMESH_LIGHTING_LIGHTNESS_SERVER_MODEL_ID
            and evt.elem_index == 0):
            self.lightness_change(evt)
        elif (evt.model_id == model.BTMESH_LIGHTING_CTL_SERVER_MODEL_ID
            and evt.elem_index == 0):
            self.ctl_change(evt)

        ####################################
        # Add further event handlers here.
        ####################################

    def load_to_dataclass(self,
                          lighting_state_to: dataclass,
                          lighting_state_from: tuple):
        """ Convert the data from nvm to the given dataclass structure. """
        count = 0
        for field in fields(lighting_state_to):
            if getattr(lighting_state_to, field.name) != lighting_state_from[count]:
                setattr(lighting_state_to, field.name, lighting_state_from[count])
            count += 1

    def restore_nvm_last_sate(self):
        """
        Get the last state of the Lighting server and CTL server.
        Check if the nvm have the necessary keys to save the data.
        """
        try:
            lighting_state_nvm_load = self.lib.bt.nvm.load(self.lighting_server_key)
            self.NVMState.lighting_state_nvm = struct.unpack("<BBHHHHHHHHhh",
                                                    lighting_state_nvm_load.value)
            
            self.log.info(f"From nvm {self.NVMState.lighting_state_nvm}")
            self.load_to_dataclass(self.CTLLightbulbLightness,
                                   self.NVMState.lighting_state_nvm)
            
            # Update GUI
            self.lighting_set_level(self.CTLLightbulbLightness.lightness_current,
                                    IMMEDIATE)
        except CommandFailedError as e:
            # Can not find data in nvm with the given key
            if e.errorcode != status.BT_PS_KEY_NOT_FOUND:
                raise
            self.log.warning("Lighting server nvm data was not found")

        try:
            ctl_state_nvm_load = self.lib.bt.nvm.load(self.ctl_server_key)
            self.NVMState.ctl_state_nvm = struct.unpack("<HHHHHhhhhh", ctl_state_nvm_load.value)
            self.log.info(f"{self.NVMState.ctl_state_nvm}")
            self.load_to_dataclass(self.CTLLightbulbTemperature,
                                   self.NVMState.ctl_state_nvm)
            # Update GUI
            self.set_temperature_deltauv_level(self.CTLLightbulbTemperature.temperature_current,
                                               self.CTLLightbulDeltaUV.deltauv_current)
            self.lighting_set_level(self.CTLLightbulbLightness.lightness_current,
                                    IMMEDIATE)
        except CommandFailedError as e:
            # Can not find data in nvm with the given key
            if e.errorcode != status.BT_PS_KEY_NOT_FOUND:
                raise
            self.log.warning("CTL server nvm data was not found")


# Script entry point.
if __name__ == "__main__":
    # Instantiate and run application.
    parser = ArgumentParser(description=__doc__)
    args = parser.parse_args()
    connector = get_connector(args)
    app = App(connector=connector)
    t_app = threading.Thread(target=app.run, args=[], daemon=True)
    t_app.name = "App_thread"
    t_app.start()
    lighting_server_gui_start(app)
```
