## Supplementary material for "VesperFlow: An automated pipeline for decoding multisensor data from biologging tags": ctl_server.py

```
#!/usr/bin/env python3
"""
BtMesh NCP Light Server, CTL Model implementation.
"""

# Copyright 2022 Silicon Laboratories Inc. www.silabs.com
#
# SPDX-License-Identifier: Zlib
#
# The licensor of this software is Silicon Laboratories Inc.
#
# This software is provided 'as-is', without any express or implied
# warranty. In no event will the authors be held liable for any damages
# arising from the use of this software.
#
# Permission is granted to anyone to use this software for any purpose,
# including commercial applications, and to alter it and redistribute it
# freely, subject to the following restrictions:
#
# 1. The origin of this software must not be misrepresented; you must not
#    claim that you wrote the original software. If you use this software
#    in a product, an acknowledgment in the product documentation would be
#    appreciated but is not required.
# 2. Altered source versions must be plainly marked as such, and must not be
#    misrepresented as being the original software.
# 3. This notice may not be removed or altered from any source distribution.

from dataclasses import dataclass
import os.path
import sys
import struct
import threading

sys.path.append(os.path.join(os.path.dirname(__file__), "../.."))
from lighting_server_gui import MainPage
from lightness_server import LightnessServer
from bgapi.bglib import CommandFailedError
import common.status as status
import common.btmesh_models as model

IMMEDIATE = 0

class CTLServer(LightnessServer):
    """ Implementation of CTL Server. """
    def __init__(self, connector, **kwargs):
        super().__init__(connector=connector, **kwargs)
        # Copy of transition delay parameter, needed for delayed ctl request
        self.delayed_ctl_trans = 0
        # CTL server key to store data to nvm
        self.ctl_server_key = 0x4005
        # CTL server nvm save timer
        self.ctl_nvm_save_timer = threading.Timer(self.nvm_save_time * 0.001,
                                                  self.ctl_lightbulb_state_changed)

    @dataclass
    class CTLLightbulbTemperature:
        """ Dataclass of CTL lightbulb Tempereature state. """
        # Current temperature value
        temperature_current = 0
        # Target temperature value
        temperature_target = 0
        # Default temperature value
        temperature_default = 6500
        # Minimum temperature value
        temperature_min = 800
        # Maximum temperature value
        temperature_max = 20000
    @dataclass
    class CTLLightbulDeltaUV:
        """ Dataclass of CTL lightbulb Delta UV state. """
        # Current delta UV value
        deltauv_current = 0
        # Target delta UV value
        deltauv_target = 0
        # Default delta UV value
        deltauv_default = 0
        # Current secondary generic level value
        sec_level_current = 0
        # Target secondary generic level value
        sec_level_target = 0

    def ctl_request(self, event):
        """ Process light CTL model requests. """
        param = event.parameters
        lightness_actual = (param[1] << 8) + param[0]
        temperature = (param[3] << 8) + param[2]
        deltauv = (((param[5] << 8) + param[4]).to_bytes(2, 'little'))
        deltauv = int.from_bytes(deltauv, byteorder='little', signed=True )

        self.log.info(f"ctl_request: lightness: {lightness_actual}, " +
                      f"color temperature: {temperature}, delta_uv: {deltauv}, " +
                      f"trans: {event.transition_ms}, delay: {event.delay_ms}")

        if (self.CTLLightbulbLightness.lightness_current == lightness_actual
            and self.CTLLightbulbTemperature.temperature_current == temperature
            and self.CTLLightbulDeltaUV.deltauv_current == deltauv):
            self.log.info("Request for current CTL state received.")
        else:
            # Set target value and check current state
            if self.CTLLightbulbLightness.lightness_current != lightness_actual:
                self.log.info(f"Setting lightness to {lightness_actual}")
                self.CTLLightbulbLightness.lightness_target = lightness_actual
            if self.CTLLightbulbTemperature.temperature_current != temperature:
                self.log.info(f"Setting temperature to {temperature}")
                self.CTLLightbulbTemperature.temperature_target = temperature
            if self.CTLLightbulDeltaUV.deltauv_current != deltauv:
                self.log.info(f"Setting delta UV to {deltauv}")
                self.CTLLightbulDeltaUV.deltauv_target = deltauv

            # Immediate change
            if event.transition_ms == 0 and event.delay_ms == 0:
                self.CTLLightbulbLightness.lightness_current = lightness_actual
                if lightness_actual != 0:
                    self.CTLLightbulbLightness.lightness_last = lightness_actual

                self.CTLLightbulbTemperature.temperature_current = temperature
                self.CTLLightbulDeltaUV.deltauv_current = deltauv
                # Update GUI according to the set target value.
                self.set_temperature_deltauv_level(
                    self.CTLLightbulbTemperature.temperature_current,
                    self.CTLLightbulDeltaUV.deltauv_current,
                )
                self.lighting_set_level(lightness_actual, IMMEDIATE)

            elif event.delay_ms > 0:
                # A delay has been specified for the light change. Start a timer
                # That will trigger the change after the given delay.
                # Current state remains as is for now.
                # Timer start
                threading.Timer(event.delay_ms * 0.001,
                                self.delayed_ctl_request).start()
                self.delayed_ctl_trans = event.transition_ms

            else:
                # No delay but transition time has been set.
                # Update GUI according to the set target value.
                self.set_temperature_deltauv_level(self.CTLLightbulbTemperature.temperature_target,
                                                   self.CTLLightbulDeltaUV.deltauv_target,)
                self.lighting_set_level(lightness_actual, event.transition_ms)
                threading.Timer(event.transition_ms * 0.001,
                                self.ctl_transition_complete).start()

            # Save the state in flash after a small delay
            self.ctl_nvm_save_timer_start()
            # State has changed, so the current scene number is reset
            self.lib.btmesh.scene_server.reset_register(event.elem_index)

        remaining_ms = event.delay_ms + event.transition_ms
        if event.flags & 2:
            # Response required. If non-zero, the client expects a response from the server.
            self.ctl_response(0, event.client_address, event.appkey_index, remaining_ms)

        self.ctl_update_and_publish(0, remaining_ms)


    def delayed_ctl_request(self):
        """ Handle delayed light CTL requests. """
        self.log.info("Delayed CTL request")

        self.log.info("Starting delayed CTL request: lightness: " +
                      f"{self.CTLLightbulbLightness.lightness_current} -> " +
                      f"{self.CTLLightbulbLightness.lightness_target}, " +
                      f"color temperature: {self.CTLLightbulbTemperature.temperature_current} -> " +
                      f"{self.CTLLightbulbTemperature.temperature_target}, " +
                      f"delta_uv: {self.CTLLightbulDeltaUV.deltauv_current} -> " +
                      f"{self.CTLLightbulDeltaUV.deltauv_target}, " +
                      f"delay: {self.delayed_ctl_trans}ms")

        # Update GUI according to the set target value
        self.set_temperature_deltauv_level(self.CTLLightbulbTemperature.temperature_target,
                                           self.CTLLightbulDeltaUV.deltauv_target,)
        self.lighting_set_level(self.CTLLightbulbLightness.lightness_target,
                                self.delayed_ctl_trans)

        if self.delayed_ctl_trans == 0:
            # No transition delay, update state immediately
            self.CTLLightbulbLightness.lightness_current = (
                self.CTLLightbulbLightness.lightness_target)
            self.CTLLightbulbTemperature.temperature_current = (
                self.CTLLightbulbTemperature.temperature_target)
            self.CTLLightbulDeltaUV.deltauv_current = (
                self.CTLLightbulDeltaUV.deltauv_target)
            # Save the state in flash after a small delay
            self.ctl_nvm_save_timer_start()
            self.ctl_update_and_publish(0, self.delayed_ctl_trans)

        else:
            # State is updated when transition is complete
            threading.Timer(self.delayed_ctl_trans * 0.001,
                            self.ctl_transition_complete).start()

    def ctl_update(self, elem_index, remaining_ms):
        """ Update light CTL state. """
        self.log.info("CTL update")
        ctl = struct.pack("<HHhHHh",
                          self.CTLLightbulbLightness.lightness_current,
                          self.CTLLightbulbTemperature.temperature_current,
                          self.CTLLightbulDeltaUV.deltauv_current,
                          self.CTLLightbulbLightness.lightness_target,
                          self.CTLLightbulbTemperature.temperature_target,
                          self.CTLLightbulDeltaUV.deltauv_target)

        self.lib.btmesh.generic_server.update(
            elem_index,
            model.BTMESH_LIGHTING_CTL_SERVER_MODEL_ID,
            remaining_ms,
            self.lib.btmesh.generic_client.GET_STATE_TYPE_STATE_CTL,
            ctl)

    def ctl_temperature_update(self, elem_index, remaining_ms):
        """ Update light CTL temperature state. Needed for scene recall. """
        self.log.info("CTL temperature update")
        ctl = struct.pack("<HhHh",
                          self.CTLLightbulbTemperature.temperature_current,
                          self.CTLLightbulDeltaUV.deltauv_current,
                          self.CTLLightbulbTemperature.temperature_target,
                          self.CTLLightbulDeltaUV.deltauv_target,)

        try:
            self.lib.btmesh.generic_server.update(
                elem_index,
                model.BTMESH_LIGHTING_CTL_TEMPERATURE_SERVER_MODEL_ID,
                remaining_ms,
                self.lib.btmesh.generic_client.GET_STATE_TYPE_STATE_CTL_TEMPERATURE,
                ctl)
        except CommandFailedError as e:
            if e.errorcode != status.BT_MESH_DOES_NOT_EXIST:
                raise

    def ctl_response(self, elem_index, client_addr, appkey_index, remaining_ms):
        """ Respond to light CTL request. """
        self.log.info("CTL_response")
        ctl = struct.pack("<HHhHHh",
                          self.CTLLightbulbLightness.lightness_current,
                          self.CTLLightbulbTemperature.temperature_current,
                          self.CTLLightbulDeltaUV.deltauv_current,
                          self.CTLLightbulbLightness.lightness_target,
                          self.CTLLightbulbTemperature.temperature_target,
                          self.CTLLightbulDeltaUV.deltauv_target)

        self.lib.btmesh.generic_server.respond(
            client_addr,
            elem_index,
            model.BTMESH_LIGHTING_CTL_SERVER_MODEL_ID,
            appkey_index,
            remaining_ms,
            IMMEDIATE,
            self.lib.btmesh.generic_client.GET_STATE_TYPE_STATE_CTL,
            ctl,
        )

    def ctl_update_and_publish(self, elem_index, remaining_ms):
        """ Update light CTL state and publish model state to the network. """
        self.log.info("CTL update and publish")
        try:
            self.ctl_update(elem_index, remaining_ms)

            self.lib.btmesh.generic_server.publish(
                elem_index,  # 0
                model.BTMESH_LIGHTING_CTL_SERVER_MODEL_ID,
                self.lib.btmesh.generic_client.GET_STATE_TYPE_STATE_CTL,
            )
        except CommandFailedError as err:
            self.log.error("%s", err.errorcode)

    def ctl_transition_complete(self):
        """ Callback to Light CTL request with non-zero transition time. """
        self.log.info("CTL transition complete")
        self.CTLLightbulbLightness.lightness_current = (
            self.CTLLightbulbLightness.lightness_target)
        self.CTLLightbulbTemperature.temperature_current = (
            self.CTLLightbulbTemperature.temperature_target)
        self.CTLLightbulDeltaUV.deltauv_current = (
            self.CTLLightbulDeltaUV.deltauv_target)

        self.log.info("Transition complete. New lightness is " +
                      f"{self.CTLLightbulbLightness.lightness_current} " +
                      f"new color temperature: {self.CTLLightbulbTemperature.temperature_current} " +
                      f"new delta_uv: {self.CTLLightbulDeltaUV.deltauv_current}")

        # Save the state in flash after a small delay
        self.ctl_nvm_save_timer_start()
        self.ctl_update_and_publish(0, IMMEDIATE)

    def ctl_recall(self, event):
        """ Handle light CTL change events. """
        self.log.info("CTL recall")
        param = event.parameters
        lightness_actual = (param[1] << 8) + param[0]
        temperature = (param[3] << 8) + param[2]
        deltauv = (((param[5] << 8) + param[4]).to_bytes(2, 'little'))
        deltauv = int.from_bytes(deltauv, byteorder='little', signed=True)

        if (self.CTLLightbulbLightness.lightness_current == lightness_actual
            and self.CTLLightbulbTemperature.temperature_current == temperature
            and self.CTLLightbulDeltaUV.deltauv_current == deltauv):
            self.log.info("Request for current CTL state received.")
        else:
            # Set target value and check current state
            if self.CTLLightbulbLightness.lightness_current != lightness_actual:
                self.log.info(f"Setting lightness to {lightness_actual}")
                self.CTLLightbulbLightness.lightness_target = lightness_actual
            if self.CTLLightbulbTemperature.temperature_current != temperature:
                self.log.info(f"Setting temperature to {temperature}")
                self.CTLLightbulbTemperature.temperature_target = temperature
            if self.CTLLightbulDeltaUV.deltauv_current != deltauv:
                self.log.info(f"Setting delta UV to {deltauv}")
                self.CTLLightbulDeltaUV.deltauv_target = deltauv

            if event.transition_time_ms == 0:
                self.CTLLightbulbLightness.lightness_current = lightness_actual
                self.CTLLightbulbTemperature.temperature_current = temperature
                self.CTLLightbulDeltaUV.deltauv_current = deltauv

                self.set_temperature_deltauv_level(self.CTLLightbulbTemperature.temperature_current,
                                                   self.CTLLightbulDeltaUV.deltauv_current)
                self.lighting_set_level(self.CTLLightbulbLightness.lightness_target,
                                        event.transition_time_ms)

            else:
                threading.Timer(event.delay_ms * 0.001,
                                self.ctl_transition_complete).start()

            # Save the state in flash after a small delay
            self.ctl_nvm_save_timer_start()

        self.ctl_temperature_update(event.elem_index, event.transition_time_ms)

        try:
            self.lib.btmesh.generic_server.publish(
                0,
                model.BTMESH_LIGHTING_CTL_SERVER_MODEL_ID,
                self.lib.btmesh.generic_client.GET_STATE_TYPE_STATE_CTL,
            )
        except CommandFailedError as e:
            if e.errorcode != status.BT_MESH_PUBLISH_NOT_CONFIGURED:
                raise

    def ctl_change(self, event):
        """ Handle light CTL change event. """
        if event.type != self.lib.btmesh.generic_client.GET_STATE_TYPE_STATE_CTL:
            return

        self.log.info("CTL change")
        self.log.debug(event)
        param = event.parameters
        lightness_actual = (param[1] << 8) + param[0]
        temperature = (param[3] << 8) + param[2]
        deltauv = (((param[5] << 8) + param[4]).to_bytes(2, 'little'))
        deltauv = int.from_bytes(deltauv, byteorder='little', signed=True)

        # Lightness check
        if self.CTLLightbulbLightness.lightness_current == lightness_actual:
            self.log.info(f"Lightness update same value, {lightness_actual}")
        else:
            self.log.info(
                f"Lightness value update: from {self.CTLLightbulbLightness.lightness_current} \
                to {lightness_actual}"
            )
            self.CTLLightbulbLightness.lightness_current = lightness_actual
            # Save the state in flash after a small delay
            self.ctl_nvm_save_timer_start()

        # Temperature check
        if self.CTLLightbulbTemperature.temperature_current != temperature:
            self.log.info("Color temperature update: " +
                          f"from {self.CTLLightbulbTemperature.temperature_current} " +
                          f"to {temperature}")
            self.CTLLightbulbTemperature.temperature_current = temperature
            # Save the state in flash after a small delay
            self.ctl_nvm_save_timer_start()
        else:
            self.log.info(f"Color temperature update same value, {temperature}")

        # Delta UV check
        if self.CTLLightbulDeltaUV.deltauv_current != deltauv:
            self.log.info("Delta UV value update: "+ 
                          f"from {self.CTLLightbulDeltaUV.deltauv_current} " +
                          f"to {deltauv}")
            self.CTLLightbulDeltaUV.deltauv_current = deltauv
            # Save the state in flash after a small delay
            self.ctl_nvm_save_timer_start()
        else:
            self.log.info(f"Delta UV update same value, {deltauv}")

    def ctl_lightbulb_state_changed(self):
        """ Save the state to the nvm after the ctl lightbulb state is changed."""
        self.lib.bt.nvm.save (
            self.ctl_server_key,
            self.ctl_state_serialize()
        )

    def set_temperature_deltauv_level(self, temperature, delta_uv):
        """ Set GUI temperature and delta UV in given transition time. """
        if (self.CTLLightbulbTemperature.temperature_current <
            self.CTLLightbulbTemperature.temperature_min):
            self.CTLLightbulbTemperature.temperature_current = \
            self.CTLLightbulbTemperature.temperature_min
        elif (self.CTLLightbulbTemperature.temperature_current >
              self.CTLLightbulbTemperature.temperature_max):
            self.CTLLightbulbTemperature.temperature_current = \
            self.CTLLightbulbTemperature.temperature_max

        MainPage.get_instance(self).set_temperature_value(temperature)
        MainPage.get_instance(self).set_delta_uv_value(self.display_delta_uv(delta_uv))

    ##### HELPER FUNCTIONS #####

    def ctl_nvm_save_timer_start(self):
        """ Start or restart CTL NVM save timer. """
        if self.ctl_nvm_save_timer.is_alive():
            self.ctl_nvm_save_timer.cancel()
        self.ctl_nvm_save_timer = threading.Timer(self.nvm_save_time * 0.001,
                                                  self.ctl_lightbulb_state_changed)
        self.ctl_nvm_save_timer.start()

    def display_delta_uv(self, delta_uv):
        """ Convert delta UV raw value to display. """
        self.log.info(delta_uv)
        signed_delta = delta_uv
        if 32786 <= delta_uv:
            signed_delta = delta_uv - 65536
        return signed_delta / 32767

    def ctl_state_serialize(self):
        """ Serialize CTL state. """
        ctl_state = struct.pack("<HHHHHhhhhh",
                                self.CTLLightbulbTemperature.temperature_current,
                                self.CTLLightbulbTemperature.temperature_target,
                                self.CTLLightbulbTemperature.temperature_default,
                                self.CTLLightbulbTemperature.temperature_min,
                                self.CTLLightbulbTemperature.temperature_max,
                                self.CTLLightbulDeltaUV.deltauv_current,
                                self.CTLLightbulDeltaUV.deltauv_target,
                                self.CTLLightbulDeltaUV.deltauv_default,
                                self.CTLLightbulDeltaUV.sec_level_current,
                                self.CTLLightbulDeltaUV.sec_level_target)
        return ctl_state
```
