## Supplementary material for "VesperFlow: An automated pipeline for decoding multisensor data from biologging tags": factory_reset.py

```
"""
BtMesh NCP Light Server General APIs implementation.
"""

# Copyright 2022 Silicon Laboratories Inc. www.silabs.com
#
# SPDX-License-Identifier: Zlib
#
# The licensor of this software is Silicon Laboratories Inc.
#
# This software is provided 'as-is', without any express or implied
# warranty. In no event will the authors be held liable for any damages
# arising from the use of this software.
#
# Permission is granted to anyone to use this software for any purpose,
# including commercial applications, and to alter it and redistribute it
# freely, subject to the following restrictions:
#
# 1. The origin of this software must not be misrepresented; you must not
#    claim that you wrote the original software. If you use this software
#    in a product, an acknowledgment in the product documentation would be
#    appreciated but is not required.
# 2. Altered source versions must be plainly marked as such, and must not be
#    misrepresented as being the original software.
# 3. This notice may not be removed or altered from any source distribution.

import time
import os.path
import sys

sys.path.append(os.path.join(os.path.dirname(__file__), "../.."))
from common.util import BtMeshApp

class FactoryReset(BtMeshApp):
    """ Implementation of the general APIs. """
    # Node reset
    def node_reset(self):
        """ Initiate node reset."""
        # Perform a factory reset of the node. This removes all the keys
        # and other settings that have been configured for this node
        self.lib.btmesh.node.reset()
        # 2 seconds delay is required to finalize node reset
        time.sleep(2)
        self.lib.bt.system.reboot()
        self.log.info("Node reset")

    # Full factory reset
    def factory_reset(self):
        """ Initiate full factory reset."""
        # Perform a factory reset of the node. This removes all the keys
        # and other settings that have been configured for this node and
        # erases the NVM
        self.lib.btmesh.node.reset()
        self.lib.bt.nvm.erase_all()
        # 2 seconds delay is required to finalize node reset
        time.sleep(2)
        self.lib.bt.system.reboot()
        self.log.info("Factory reset")
```
