## Supplementary material for "VesperFlow: An automated pipeline for decoding multisensor data from biologging tags": lighting_server_gui.py

```
"""
BtMesh NCP Light Server GUI.
"""

# Copyright 2022 Silicon Laboratories Inc. www.silabs.com
#
# SPDX-License-Identifier: Zlib
#
# The licensor of this software is Silicon Laboratories Inc.
#
# This software is provided 'as-is', without any express or implied
# warranty. In no event will the authors be held liable for any damages
# arising from the use of this software.
#
# Permission is granted to anyone to use this software for any purpose,
# including commercial applications, and to alter it and redistribute it
# freely, subject to the following restrictions:
#
# 1. The origin of this software must not be misrepresented; you must not
#    claim that you wrote the original software. If you use this software
#    in a product, an acknowledgment in the product documentation would be
#    appreciated but is not required.
# 2. Altered source versions must be plainly marked as such, and must not be
#    misrepresented as being the original software.
# 3. This notice may not be removed or altered from any source distribution.

from dataclasses import dataclass
import tkinter
import os.path
import logging
import colour
from PIL import ImageTk, Image

class ResizableWindow(tkinter.Tk):
    """ Root window for the frame. """
    def __init__(self, app, *args, **kwargs):
        tkinter.Tk.__init__(self, *args, **kwargs)
        self.protocol("WM_DELETE_WINDOW", lambda args=app: self.exit(args))
        self.resizable(1, 1)

        # Adding a title to the window
        self.wm_title("BT Mesh Light")

        # Specifying the region where the frame is grided
        self.geometry("350x160")
        self.rowconfigure(0, weight=1)
        self.columnconfigure(0, weight=1)

        frame = MainPage(self, app)
        frame.grid(row=0, column=0, sticky=tkinter.NSEW)

    def exit(self, app):
        """ Shuting down the app. """
        app.stop()
        self.destroy()

class MainPage(tkinter.Frame):
    """Implementation of the used frame."""
    __instance = None
    def get_instance(self):
        """ Getting GUI instance. """
        if MainPage.__instance is None:
            self.gui_log.info("No instance for GUI yet")
        return MainPage.__instance

    def __init__(self, parent, app):
        self.gui_log = logging.getLogger(str(type(self)))
        tkinter.Frame.__init__(self, parent)
        if MainPage.__instance is not None:
            self.gui_log.error("GUI instance is already present")
            return
        self.root = self
        self.app = app
        MainPage.__instance = self

        # Variables
        self.CTLLightbulbState.lightness_value = tkinter.IntVar()
        self.CTLLightbulbState.temperature_value = tkinter.IntVar()
        self.CTLLightbulbState.temperature_value.set(6500)
        self.CTLLightbulbState.delta_uv_value = tkinter.DoubleVar()
        # Image size
        pixels_x = 100
        pixels_y = 100

        # Lightness
        slider_label = tkinter.Label(
            self,
            text='Lightness:'
        )

        slider_label.grid(
            row=0,
            column=1,
            sticky=tkinter.EW
        )

        self.lightness_value_label = tkinter.Label(
            self,
            text=f"{self.CTLLightbulbState.lightness_value.get()}%"
        )

        self.lightness_value_label.grid(
            row=0,
            column=2,
            sticky=tkinter.EW,
            padx=5,
            pady=5
        )

        # Temperature
        slider_label = tkinter.Label(
            self,
            text='Temperature:'
        )

        slider_label.grid(
            column=1,
            row=1,
            sticky=tkinter.EW
        )

        self.temperature_value_label = tkinter.Label(
            self,
            text=f"{self.CTLLightbulbState.temperature_value.get()}K"
        )

        self.temperature_value_label.grid(
            row=1,
            column=2,
            sticky=tkinter.EW,
            padx=5,
            pady=5
        )

        # Delta UV
        slider_label = tkinter.Label(
            self,
            text='Delta UV:'
        )

        slider_label.grid(
            column=1,
            row=2,
            sticky=tkinter.EW
        )

        self.delta_uv_value_label = tkinter.Label(
            self,
            text=f"{self.CTLLightbulbState.delta_uv_value.get():.2f}"
        )
        self.delta_uv_value_label.grid(
            row=2,
            column=2,
            sticky=tkinter.EW,
            padx=5,
            pady=5
        )

        # Factory reset button
        factory_reset = tkinter.Button(
            self,
            text="Factory reset",
            command=self.factory_reset
        )

        factory_reset.grid(
            row=3,
            column=2,
            sticky=tkinter.EW,
            padx=5,
            pady=5
        )

        # Node reset button
        node_reset = tkinter.Button(
            self,
            text="Node reset",
            command=self.node_reset
        )

        node_reset.grid(
            row=3,
            column=0,
            sticky=tkinter.EW,
            padx=5,
            pady=5
        )

        project_directory = os.path.realpath(os.path.join(os.path.dirname(__file__), '..'))
        image_path = Image.open(os.path.join(project_directory, 'btmesh_light', 'images',
                                            'lightbulb.png'))
        self.load_image = ImageTk.PhotoImage(image_path.resize((pixels_x, pixels_y)))
        self.lightbulb_image = tkinter.Label(self, image=self.load_image, borderwidth=4, bg="black")

        self.lightbulb_image.grid(
            row=0,
            column=0,
            rowspan=3,
            padx=5,
            pady=5
        )

        self.columnconfigure((0,1,2), weight=1, uniform='column')
        self.rowconfigure((0,1,2,3), weight=1, uniform='row')

    @dataclass
    class CTLLightbulbState:
        """ Dataclass of CTL ligthbulb state. """
        # Value of lightness
        lightness_value = None
        # Value of temperature
        temperature_value = None
        # Value of deltauv
        delta_uv_value = None

    def set_lightness_value(self, value):
        """ Set the Lightness value and updates the GUI. """
        self.CTLLightbulbState.lightness_value.set(value)
        self.lightness_value_label.configure(text=
                                             f"{self.CTLLightbulbState.lightness_value.get()}%")

    def set_temperature_value(self, value):
        """ Set the Temperature value and updates the GUI. """
        self.CTLLightbulbState.temperature_value.set(value)
        self.temperature_value_label.configure(text=
                                               f"{self.CTLLightbulbState.temperature_value.get()}K")

    def set_delta_uv_value(self, value):
        """ Set the Delta UV value and updates the GUI. """
        self.CTLLightbulbState.delta_uv_value.set(value)
        self.delta_uv_value_label.configure(text=
                                            f"{self.CTLLightbulbState.delta_uv_value.get():.2f}")

    def factory_reset(self):
        """ Call the factory reset function from the General class. """
        self.set_lightness_value(0)
        self.set_temperature_value(6500)
        self.set_delta_uv_value(0)
        self.fade(self.lightbulb_image, "#000000", 0)
        self.app.factory_reset()

    def node_reset(self):
        """ Call the node reset function from the General class. """
        self.set_lightness_value(0)
        self.set_temperature_value(6500)
        self.set_delta_uv_value(0)
        self.fade(self.lightbulb_image, "#000000", 0)
        self.app.node_reset()

    def fade(self, widget, bg, trans_ms):
        """
        Show faded effect on widget's different color options.

        :param widget: image with no background
        :param bg: fade background color to the given color (str)
        :param trans_ms: transition time in ms
        """
        if trans_ms == 0:
            widget.config(bg=bg)
        else:
            if not getattr(widget, "_after_ids", None):
                widget._after_ids = {}
            widget.after_cancel(widget._after_ids.get("bg", " "))
            c1 = tuple(map(lambda a: a / (65535), widget.winfo_rgb(widget["bg"])))
            c2 = tuple(map(lambda a: a / (65535), widget.winfo_rgb(bg)))

            colors = tuple(
                colour.rgb2hex(c, force_long=True)
                for c in colour.color_scale(
                    c1, c2, max(1, int((trans_ms * 60) // 1000))
                )
            )

            def worker(count=0):
                if len(colors) - 1 <= count:
                    return
                widget.config({"bg": colors[count]})
                widget._after_ids.update(
                    {"bg": widget.after(1000 // 60, worker, count + 1)}
                )

            worker()

def lighting_server_gui_start(app):
    """ Starting the GUI. """
    gui = ResizableWindow(app=app)
    gui.mainloop()
```
