## Supplementary material for "VesperFlow: An automated pipeline for decoding multisensor data from biologging tags": lightness_server.py

```
#!/usr/bin/env python3
"""
BtMesh NCP Light Server, Lightness Model implementation.
"""

# Copyright 2022 Silicon Laboratories Inc. www.silabs.com
#
# SPDX-License-Identifier: Zlib
#
# The licensor of this software is Silicon Laboratories Inc.
#
# This software is provided 'as-is', without any express or implied
# warranty. In no event will the authors be held liable for any damages
# arising from the use of this software.
#
# Permission is granted to anyone to use this software for any purpose,
# including commercial applications, and to alter it and redistribute it
# freely, subject to the following restrictions:
#
# 1. The origin of this software must not be misrepresented; you must not
#    claim that you wrote the original software. If you use this software
#    in a product, an acknowledgment in the product documentation would be
#    appreciated but is not required.
# 2. Altered source versions must be plainly marked as such, and must not be
#    misrepresented as being the original software.
# 3. This notice may not be removed or altered from any source distribution.

from dataclasses import dataclass
import os.path
import sys
import threading
import math
import struct
from onoff_server import OnOffServer
from bgapi.bglib import CommandFailedError

sys.path.append(os.path.join(os.path.dirname(__file__), "../.."))
import common.status as status
import common.btmesh_models as model

IMMEDIATE = 0

class LightnessServer(OnOffServer):
    """ Implementation of lightness server. """
    def __init__(self, connector, **kwargs):
        super().__init__(connector=connector, **kwargs)

        # Copy of lightness request kind, needed for delayed lightness request
        self.lightness_kind = (
            self.lib.btmesh.generic_client.GET_STATE_TYPE_STATE_LIGHTNESS_ACTUAL)
        # Copy of generic request kind, needed for delayed primary generic request
        self.pri_level_request_kind = (
            self.lib.btmesh.generic_client.SET_REQUEST_TYPE_REQUEST_LEVEL)
        # Move delta parameter for primary generic request
        self.move_pri_level_delta = 0
        # Requested lightness level
        self.request_level = 0
        # Actual lightness value
        self.lightness = 0
    @dataclass
    class LightnessServerTiming:
        """ Dataclass of Lightness Server timing. """
        # Values greater than 37200000 are treated as unknown remaining time
        unknown_remaining_time = 40000000
        # Copy of delayed pri level timer
        delayed_pri_level_timer = None
        # Copy of level move timer
        level_move_timer = None
        # Move transition parameter for primary generic request
        move_pri_level_trans = 0
        # Copy of delayed pri level transition delay parameter
        delayed_pri_level_trans = 0
        # Delayed Lightness transition time
        delayed_lightness_trans = 0

    def lightness_request(self, event):
        """ Process light lightness model requests. """
        if event.type == self.lib.btmesh.generic_client.GET_STATE_TYPE_STATE_LIGHTNESS_ACTUAL:
            self.lightness_kind = \
            self.lib.btmesh.generic_client.GET_STATE_TYPE_STATE_LIGHTNESS_ACTUAL
            actual_request = int.from_bytes(event.parameters, byteorder='little')
        elif event.type == self.lib.btmesh.generic_client.SET_REQUEST_TYPE_REQUEST_LIGHTNESS_LINEAR:
            self.lightness_kind = \
            self.lib.btmesh.generic_client.SET_REQUEST_TYPE_REQUEST_LIGHTNESS_LINEAR
            actual_request = linear2actual(int.from_bytes(event.parameters, byteorder='little'))

        self.log.info(f"Lightness_request: level = {actual_request}, " +
                      f"trans = {event.transition_ms}, " +
                      f"delay = {event.delay_ms}, type = {self.lightness_kind}")

        if self.CTLLightbulbLightness.lightness_current == actual_request:
            self.log.info("Request for current Light Lightness state received.")

        else:
            self.log.info(f"Setting lightness to {actual_request}")
            if event.transition_ms == 0 and event.delay_ms == 0:
                # Immediate change
                self.CTLLightbulbLightness.lightness_current = actual_request
                self.CTLLightbulbLightness.lightness_target = actual_request

                if actual_request != 0:
                    self.CTLLightbulbLightness.lightness_last = actual_request

                self.lighting_set_level(self.CTLLightbulbLightness.lightness_current,
                                        IMMEDIATE)

            elif event.delay_ms > 0:
                # A delay has been specified for the light change.
                # Current state remains as is for now.
                self.CTLLightbulbLightness.lightness_target = actual_request
                self.LightnessServerTiming.delayed_lightness_trans = event.transition_ms
                # Timer start
                threading.Timer(event.delay_ms * 0.001,
                                self.delayed_lightness_request).start()

            else:
                # No delay but transition time has been set.
                self.CTLLightbulbLightness.lightness_target = actual_request
                self.lighting_set_level(self.CTLLightbulbLightness.lightness_target,
                                        event.transition_ms)
                threading.Timer(event.transition_ms * 0.001,
                                self.lighting_transition_complete).start()

            # Save the state in flash after a small delay
            self.lighting_nvm_save_timer_start()
            # State has changed, so the current scene number is reset
            self.lib.btmesh.scene_server.reset_register(event.elem_index)

        remaining_ms = event.delay_ms + event.transition_ms
        if event.flags & 2:
            # Response required. If non-zero, the client expects a response from the server.
            self.lightness_response(event.elem_index,
                                    event.client_address,
                                    event.appkey_index,
                                    remaining_ms)

        self.lightness_update_and_publish(event.elem_index, self.delayed_onoff_trans)

        # Publish to bound states
        if (self.lightness_kind ==
            self.lib.btmesh.generic_client.GET_STATE_TYPE_STATE_LIGHTNESS_ACTUAL):
            self.lib.btmesh.generic_server.publish(
                0,
                model.BTMESH_LIGHTING_LIGHTNESS_SERVER_MODEL_ID,
                self.lib.btmesh.generic_client.GET_STATE_TYPE_STATE_LIGHTNESS_LINEAR,
            )
        else:
            self.lib.btmesh.generic_server.publish(
                0,
                model.BTMESH_LIGHTING_LIGHTNESS_SERVER_MODEL_ID,
                self.lib.btmesh.generic_client.GET_STATE_TYPE_STATE_LIGHTNESS_ACTUAL,
            )

        self.lib.btmesh.generic_server.publish(
            0,
            model.BTMESH_GENERIC_ON_OFF_SERVER_MODEL_ID,
            self.lib.btmesh.generic_client.GET_STATE_TYPE_STATE_ON_OFF,
        )

        try:
            self.lib.btmesh.generic_server.publish(
                0,
                model.BTMESH_GENERIC_LEVEL_SERVER_MODEL_ID,
                self.lib.btmesh.generic_client.GET_STATE_TYPE_STATE_LEVEL,
            )
        except CommandFailedError as e:
            # Application key or publish address are not set
            if e.errorcode != status.BT_MESH_PUBLISH_NOT_CONFIGURED:
                raise

        try:
            self.lib.btmesh.generic_server.publish(
                0,
                model.BTMESH_LIGHTING_CTL_SERVER_MODEL_ID,
                self.lib.btmesh.generic_client.GET_STATE_TYPE_STATE_CTL,
            )
        except CommandFailedError as e:
            # Application key or publish address are not set
            if e.errorcode != status.BT_MESH_PUBLISH_NOT_CONFIGURED:
                raise

    def delayed_lightness_request(self):
        """Handle delayed light lightness requests."""
        self.log.info("Starting delayed lightness request: "+
                      f"level {self.CTLLightbulbLightness.lightness_current} -> " +
                      f"{self.CTLLightbulbLightness.lightness_target}, " +
                      f"{self.LightnessServerTiming.delayed_lightness_trans} ms")

        self.lighting_set_level(self.CTLLightbulbLightness.lightness_target,
                                self.LightnessServerTiming.delayed_lightness_trans)
        if self.LightnessServerTiming.delayed_lightness_trans == 0:
            # No transition delay, update state immediately
            self.CTLLightbulbLightness.lightness_current = (
                self.CTLLightbulbLightness.lightness_target)
            if self.CTLLightbulbLightness.lightness_target != 0:
                self.CTLLightbulbLightness.lightness_last = (
                    self.CTLLightbulbLightness.lightness_target)

            # Save the state in flash after a small delay
            self.lighting_nvm_save_timer_start()
            self.lightness_update_and_publish(
                0,
                self.LightnessServerTiming.delayed_lightness_trans)
        else:
            # State is updated when transition is complete
            threading.Timer(self.LightnessServerTiming.delayed_lightness_trans * 0.001,
                            self.lighting_transition_complete).start()

    def lightness_update(self, elem_index, remaining_ms):
        """ Update light lightness state. """
        lightness = struct.pack(
            "<HH",
            self.CTLLightbulbLightness.lightness_current,
            self.CTLLightbulbLightness.lightness_target,
        )

        self.lib.btmesh.generic_server.update(
            elem_index,
            model.BTMESH_LIGHTING_CTL_TEMPERATURE_SERVER_MODEL_ID,
            remaining_ms,
            self.lightness_kind,
            lightness,
        )

    def lightness_response(self, elem_index, client_addr, appkey_index, remaining_ms):
        """Response to light lightness request."""
        if (self.lightness_kind
            == self.lib.btmesh.generic_client.GET_STATE_TYPE_STATE_LIGHTNESS_ACTUAL):
            lightness = struct.pack(
                "<HH",
                self.CTLLightbulbLightness.lightness_current,
                self.CTLLightbulbLightness.lightness_target,
            )
        else:
            lightness = struct.pack(
                "<HH",
                actual2linear(self.CTLLightbulbLightness.lightness_current),
                actual2linear(self.CTLLightbulbLightness.lightness_target),
            )

        self.lib.btmesh.generic_server.respond(
            client_addr,
            elem_index,  # 0
            model.BTMESH_LIGHTING_LIGHTNESS_SERVER_MODEL_ID,
            appkey_index,
            remaining_ms,
            0x00,
            self.lightness_kind,
            lightness,
        )

    def lightness_update_and_publish(self, elem_index, remaining_ms):
        """ Update light lightness state and publish model state to the network. """
        self.lightness_update(elem_index, remaining_ms)

        try:
            self.lib.btmesh.generic_server.publish(
                elem_index,  # 0
                model.BTMESH_LIGHTING_LIGHTNESS_SERVER_MODEL_ID,
                self.lightness_kind,
            )
        except CommandFailedError as e:
            # Application key or publish address are not set
            if e.errorcode != status.BT_MESH_PUBLISH_NOT_CONFIGURED:
                raise

    def lighting_transition_complete(self):
        """ Callback to light lightness request with non-zero transition time. """
        self.CTLLightbulbLightness.lightness_current = (
            self.CTLLightbulbLightness.lightness_target
        )
        if self.CTLLightbulbLightness.lightness_target != 0:
            self.CTLLightbulbLightness.lightness_last = (
                self.CTLLightbulbLightness.lightness_target
            )
        self.log.info("Transition complete. New level is " +
                      f"{self.CTLLightbulbLightness.lightness_current}")
        # Save the state in flash after a small delay
        self.lighting_nvm_save_timer_start()
        self.lightness_update_and_publish(0, IMMEDIATE)

    def lightness_recall(self, event):
        """ Handle light lightness recall events. """
        if (event.type
            != self.lib.btmesh.generic_client.GET_STATE_TYPE_STATE_LIGHTNESS_ACTUAL):
            return

        self.lightness_kind = self.lib.btmesh.generic_client.GET_STATE_TYPE_STATE_LIGHTNESS_ACTUAL
        actual_request = int.from_bytes(event.parameters, byteorder="little")
        self.CTLLightbulbLightness.lightness_target = actual_request

        if (self.CTLLightbulbLightness.lightness_current
            == self.CTLLightbulbLightness.lightness_target):
            self.log.info("Request for current Light Lightness state received.")
        else:
            self.log.info(f"Recall lightness to {self.CTLLightbulbLightness.lightness_target} \
                with transition {event.transition_time_ms}")
            self.lighting_set_level(self.CTLLightbulbLightness.lightness_target,
                                    event.transition_time_ms)

            if event.transition_time_ms == IMMEDIATE:
                self.CTLLightbulbLightness.lightness_current = (
                    self.CTLLightbulbLightness.lightness_target)
            else:
                # Lightbulb current state will be updated when transition is complete
                threading.Timer(event.transition_time_ms * 0.001,
                                self.lighting_transition_complete).start()

            # Save the state in flash after a small delay
            self.lighting_nvm_save_timer_start()

        self.lightness_update_and_publish(event.elem_index, event.transition_time_ms)

    def lightness_change(self, event):
        """ Handle light lightness change events. """
        if (event.type
            != self.lib.btmesh.generic_client.GET_STATE_TYPE_STATE_LIGHTNESS_ACTUAL):
            return

        param = event.parameters
        length = len(event.parameters)
        if length > 2:
            current = (param[1] << 8) + param[0]
        else:
            current = self.CTLLightbulbLightness.lightness_last

        self.log.info(f"Lightness change to {current}")
        self.lightness_kind = (
            self.lib.btmesh.generic_client.GET_STATE_TYPE_STATE_LIGHTNESS_ACTUAL)
        self.CTLLightbulbLightness.lightness_target = current

        if (self.CTLLightbulbLightness.lightness_current
            == self.CTLLightbulbLightness.lightness_target):
            self.log.info("Request for current Light Lightness state received: " +
                f"{self.CTLLightbulbLightness.lightness_current}.")
        else:
            self.log.info("Lightness update from " +
                          f"{self.CTLLightbulbLightness.lightness_current} " +
                          f"to {self.CTLLightbulbLightness.lightness_target}")
            self.CTLLightbulbLightness.lightness_current = (
                self.CTLLightbulbLightness.lightness_target)
            self.lighting_nvm_save_timer_start()

    def pri_level_request(self, event):
        """Handle generic level move request on primary element."""
        lightness = 0
        remaining_ms = self.LightnessServerTiming.unknown_remaining_time
        self.request_level = int.from_bytes(event.parameters,
                                            byteorder='little',
                                            signed=True)

        if event.type == self.lib.btmesh.generic_client.SET_REQUEST_TYPE_REQUEST_LEVEL:
            self.pri_level_move_stop()
            if self.pri_level_current == self.request_level:
                self.pri_level_target = self.request_level
            else:
                self.lightness = self.request_level + 32768
                # Immediate change
                if event.transition_ms == 0 and event.delay_ms == 0:
                    self.pri_level_current = self.request_level
                    self.pri_level_target = self.request_level
                    self.CTLLightbulbLightness.lightness_current = self.lightness
                    self.CTLLightbulbLightness.lightness_target = self.lightness
                    if self.lightness != 0:
                        self.CTLLightbulbLightness.lightness_last = lightness

                    self.lighting_set_level(self.lightness, IMMEDIATE)
                elif event.delay_ms > 0:
                    self.pri_level_target = self.request_level
                    self.CTLLightbulbLightness.lightness_target = self.lightness
                    self.pri_level_request_kind = (
                        self.lib.btmesh.generic_client.SET_REQUEST_TYPE_REQUEST_LEVEL)

                    threading.Timer(event.delay_ms * 0.001,
                                    self.delayed_pri_level_request).start()
                    self.LightnessServerTiming.delayed_pri_level_trans = event.transition_ms
                else:
                    self.CTLLightbulbLightness.pri_level_target = self.request_level
                    self.CTLLightbulbLightness.lightness_target = self.lightness
                    self.lighting_set_level(self.lightness, event.transition_ms)
                    self.lighting_transition_complete()

                # State has changed, so the current scene number is reset
                self.lib.btmesh.scene_server.reset_register(event.elem_index)

            remaining_ms = event.delay_ms + event.transition_ms

        elif (event.type
            == self.lib.btmesh.generic_client.SET_REQUEST_TYPE_REQUEST_LEVEL_MOVE):
            self.log.info(f"Pri_level_request (move): delta = {self.request_level}, " +
                f"transition = {event.transition_ms}, delay = {event.delay_ms}")
            # Store move
            self.move_pri_level_delta = self.request_level
            self.LightnessServerTiming.move_pri_level_trans = event.transition_ms

            requested_level = 0
            if self.move_pri_level_delta > 0:
                requested_level = 32767  # Max level value 0x7FFF
            elif self.move_pri_level_delta < 0:
                requested_level = -32768  # Min level value 0x8000

            if self.pri_level_current == requested_level:
                self.log.info("Request for current Generic Level state recieved")
                self.pri_level_target = requested_level
                remaining_ms = IMMEDIATE
            else:
                self.log.info(f"Setting pri_level to {requested_level}")
                lightness = requested_level + 32768

                if event.delay_ms > 0:
                    self.pri_level_target = requested_level
                    self.CTLLightbulbLightness.lightness_target = lightness
                    self.pri_level_request_kind = (
                        self.lib.btmesh.generic_client.SET_REQUEST_TYPE_REQUEST_LEVEL_MOVE)
                    self.LightnessServerTiming.delayed_pri_level_timer = threading.Timer(
                        event.delay_ms * 0.001, self.delayed_pri_level_request)
                    self.LightnessServerTiming.delayed_pri_level_timer.start()
                else:
                    # No delay to start move
                    self.pri_level_target = requested_level
                    self.CTLLightbulbLightness.lightness_target = lightness

                    remaining_delta = (
                        self.pri_level_target - self.pri_level_current)
                    self.pri_level_move_schedule_next_request(remaining_delta)

                remaining_ms = self.LightnessServerTiming.unknown_remaining_time
                # State has changed, so the current scene number is reset
                self.lib.btmesh.scene_server.reset_register(event.elem_index)

        elif (event.type
            == self.lib.btmesh.generic_client.SET_REQUEST_TYPE_REQUEST_LEVEL_HALT):
            if event.delay_ms > 0:
                remaining_ms = event.delay_ms
                threading.Timer(event.delay_ms * 0.001,
                                self.delayed_pri_level_request).start()
            else:
                self.CTLLightbulbLightness.lightness_current = (
                    self.CTLLightbulbLightness.lightness_current)
                self.CTLLightbulbLightness.lightness_target = (
                    self.CTLLightbulbLightness.lightness_current)
                self.pri_level_current = (
                    self.pri_level_current)
                self.CTLLightbulbLightness.pri_level_target = (
                    self.pri_level_current)
                self.pri_level_move_stop()
                self.lighting_set_level(
                    self.CTLLightbulbLightness.lightness_current, IMMEDIATE)
                remaining_ms = IMMEDIATE

        # Save the state in flash after a small delay
        self.lighting_nvm_save_timer_start()

        if event.flags & 2:
            # Response required. If non-zero, the client expects a response from the server.
            self.pri_level_response(event.elem_index,
                                    event.client_address,
                                    event.appkey_index,
                                    remaining_ms)

        self.pri_level_update_and_publish(event.elem_index, remaining_ms)

    def delayed_pri_level_request(self):
        """ Handle delayed generic level requests on primary element. """

        self.log.info("Starting delayed request: " +
                      f"level{self.pri_level_current} -> " +
                      f"{self.CTLLightbulbLightness.pri_level_target}, " +
                      f"{self.LightnessServerTiming.delayed_pri_level_trans} ms")

        if (self.pri_level_request_kind
            == self.lib.btmesh.generic_client.SET_REQUEST_TYPE_REQUEST_LEVEL):
            self.lighting_set_level(self.CTLLightbulbLightness.lightness_target,
                                    self.LightnessServerTiming.delayed_pri_level_trans)
            if self.LightnessServerTiming.delayed_pri_level_trans == 0:
                self.pri_level_current = self.pri_level_target
                self.CTLLightbulbLightness.lightness_current = (
                    self.CTLLightbulbLightness.lightness_target)

                if self.CTLLightbulbLightness.lightness_target != 0:
                    self.CTLLightbulbLightness.lightness_last = (
                        self.CTLLightbulbLightness.lightness_target)

                # Save the state in flash after a small delay
                self.lighting_nvm_save_timer_start()
                self.pri_level_update_and_publish(
                    0,
                    self.LightnessServerTiming.delayed_pri_level_trans)
            else:
                threading.Timer(self.LightnessServerTiming.delayed_pri_level_trans * 0.001,
                                self.lighting_transition_complete).start()

        elif (self.pri_level_request_kind
            == self.lib.btmesh.generic_client.SET_REQUEST_TYPE_REQUEST_LEVEL_MOVE):
            self.pri_level_move_schedule_next_request(
                self.pri_level_target
                - self.pri_level_current)
            self.pri_level_update_and_publish(0, self.LightnessServerTiming.unknown_remaining_time)

        elif (self.pri_level_request_kind
            == self.lib.btmesh.generic_client.SET_REQUEST_TYPE_REQUEST_LEVEL_HALT):
            self.CTLLightbulbLightness.lightness_current = (
                self.CTLLightbulbLightness.lightness_current)
            self.CTLLightbulbLightness.lightness_target = (
                self.CTLLightbulbLightness.lightness_current)
            self.pri_level_target = (
                self.pri_level_current - 32768)
            self.pri_level_target = (
                self.pri_level_current)
            self.pri_level_move_stop()
            self.lighting_set_level(self.CTLLightbulbLightness.lightness_current,
                                    IMMEDIATE)
            self.pri_level_update_and_publish(0, IMMEDIATE)

    def pri_level_update(self, elem_index, remaining_ms):
        """ Update generic level state on primary element. """
        pri_level = struct.pack(
            "<hh",
            self.pri_level_current,
            self.pri_level_target,
        )

        self.lib.btmesh.generic_server.update(
            elem_index,  # 0
            model.BTMESH_GENERIC_LEVEL_SERVER_MODEL_ID,
            remaining_ms,
            self.pri_level_request_kind,
            pri_level,
        )

    def pri_level_response(self, elem_index, client_addr, appkey_index, remaining_ms):
        """ Respond to generic level request on primary element. """
        pri_level = struct.pack(
            "<hh",
            self.pri_level_current,
            self.pri_level_target,
        )

        self.lib.btmesh.generic_server.respond(
            client_addr,
            elem_index,  # 0
            model.BTMESH_GENERIC_LEVEL_SERVER_MODEL_ID,
            appkey_index,
            remaining_ms,
            0x00,
            self.lib.btmesh.generic_client.SET_REQUEST_TYPE_REQUEST_LEVEL,
            pri_level,
        )

    def pri_level_update_and_publish(self, elem_index, remaining_ms):
        """ 
        Update generic level state on primary element 
        and publish model state to the network. 
        """
        self.pri_level_update(elem_index, remaining_ms)

        try:
            self.lib.btmesh.generic_server.publish(
                elem_index,
                model.BTMESH_GENERIC_LEVEL_SERVER_MODEL_ID,
                self.pri_level_request_kind,
            )
        except CommandFailedError as e:
            if e.errorcode != status.BT_MESH_PUBLISH_NOT_CONFIGURED:
                raise

    def pri_level_transition_complete(self):
        """Callback to a generic level request on primary element with non-zero transition time."""
        self.pri_level_current = (
            self.pri_level_target)
        self.CTLLightbulbLightness.lightness_current = (
            self.CTLLightbulbLightness.lightness_target)

        if self.CTLLightbulbLightness.lightness_target != 0:
            self.CTLLightbulbLightness.lightness_last = (
                self.CTLLightbulbLightness.lightness_target)

        self.log.info("Transition complete. New level is " +
                      f"{self.pri_level_current}")
        # Save the state in flash after a small delay
        self.lighting_nvm_save_timer_start()

        self.pri_level_update_and_publish(0, IMMEDIATE)

    def pri_level_recall(self, event):
        """ Handle generic level recall events on primary element. """
        param = event.parameters
        actual_request = ((param[1] << 8) + param[0]).to_bytes(2, 'little')
        actual_request = int.from_bytes(actual_request, byteorder='little', signed=True)
        self.CTLLightbulbLightness.pri_level_target = actual_request

        if (self.pri_level_current
            == self.pri_level_target):
            self.log.info("Request for current Generic Level state received.")
        else:
            self.log.info(f"Recall lightness to {self.pri_level_target} " +
                          f"with transition {event.transition_time_ms}")

            if event.transition_time_ms == IMMEDIATE:
                self.pri_level_current = (
                    self.pri_level_target)
            else:
                # Lightbulb current state will be updated when transition is complete
                threading.Timer(event.transition_time_ms * 0.001,
                                self.pri_level_transition_complete).start()

            # Save the state in flash after a small delay
            self.lighting_nvm_save_timer_start()

        self.pri_level_update_and_publish(event.elem_index, event.transition_time_ms)

    def pri_level_change(self, event):
        """ Handle generic level change events on primary element. """
        param = event.parameters
        current = ((param[1] << 8) + param[0]).to_bytes(2, 'little')
        current = int.from_bytes(current, byteorder='little', signed=True)

        if self.pri_level_current == current:
            self.log.info("Request for current Generic Level state received: " +
                          f"{self.pri_level_current}.")
        else:
            self.log.info("Primary level update from " +
                          f"{self.pri_level_current} " +
                          f"to {current}")
            self.pri_level_current = current
            self.lighting_nvm_save_timer_start()

    def pri_level_move_schedule_next_request(self, remaining_delta):
        """ Schedule the next generic level move request on primary element. """
        transition_ms = 0
        if abs(remaining_delta) < abs(self.move_pri_level_delta):
            transition_ms = (
                self.LightnessServerTiming.move_pri_level_trans * remaining_delta
            ) / self.move_pri_level_delta

            self.lighting_set_level(
                self.CTLLightbulbLightness.lightness_target, transition_ms)

        else:
            transition_ms = self.LightnessServerTiming.move_pri_level_trans
            self.lighting_set_level(
                self.CTLLightbulbLightness.lightness_current
                + self.move_pri_level_delta,
                self.LightnessServerTiming.move_pri_level_trans,)

        self.LightnessServerTiming.level_move_timer = threading.Timer(transition_ms * 0.001,
                                                self.pri_level_move_request)
        self.LightnessServerTiming.level_move_timer.start()

    def pri_level_move_request(self):
        """ Handle generic level move requests on primary element. """
        self.log.info(f"Primary level move: level " +
                      f"{self.pri_level_current} -> " +
                      f"{self.pri_level_target}," +
                      f"delta {self.move_pri_level_delta} " +
                      f"in {self.LightnessServerTiming.move_pri_level_trans} ms")

        remaining_delta = (
            self.pri_level_target
            - self.pri_level_current)
        if abs(remaining_delta) < abs(self.move_pri_level_delta):
            # end of move level as it reached target state
            self.pri_level_current = (
                self.pri_level_target)
            self.CTLLightbulbLightness.lightness_current = (
                self.CTLLightbulbLightness.lightness_target)

        else:
            self.pri_level_current += self.move_pri_level_delta
            self.CTLLightbulbLightness.lightness_current += self.move_pri_level_delta

        # Save the state in flash after a small delay
        self.lighting_nvm_save_timer_start()
        self.pri_level_update_and_publish(0, self.LightnessServerTiming.unknown_remaining_time)

        remaining_delta = (
            self.pri_level_target
            - self.pri_level_current)

        if remaining_delta != 0:
            self.pri_level_move_schedule_next_request(remaining_delta)

    def pri_level_move_stop(self):
        """ Stop generic level move on primary element. """
        if hasattr(self, "level_move_timer"):
            self.LightnessServerTiming.level_move_timer.cancel()
            self.move_pri_level_delta = 0
            self.LightnessServerTiming.move_pri_level_trans = 0

        elif hasattr(self, "delayed_pri_level_timer"):
            self.LightnessServerTiming.delayed_pri_level_timer.cancel()
            self.move_pri_level_delta = 0
            self.LightnessServerTiming.move_pri_level_trans = 0

##### HELPER FUNCTIONS #####

def linear2actual(linear):
    """ Convert lightness linear value to lightness actual value. """
    actual = math.sqrt(65535 * linear)
    return actual

def actual2linear(actual):
    """ Convert lightness actual value to lightness linear value. """
    linear = (actual * actual + 65534) / 65535
    return linear
```
