## Supplementary material for "VesperFlow: An automated pipeline for decoding multisensor data from biologging tags": onoff_server.py

```
#!/usr/bin/env python3
"""
BtMesh NCP Light Server, OnOff Model implementation.
"""

# Copyright 2022 Silicon Laboratories Inc. www.silabs.com
#
# SPDX-License-Identifier: Zlib
#
# The licensor of this software is Silicon Laboratories Inc.
#
# This software is provided 'as-is', without any express or implied
# warranty. In no event will the authors be held liable for any damages
# arising from the use of this software.
#
# Permission is granted to anyone to use this software for any purpose,
# including commercial applications, and to alter it and redistribute it
# freely, subject to the following restrictions:
#
# 1. The origin of this software must not be misrepresented; you must not
#    claim that you wrote the original software. If you use this software
#    in a product, an acknowledgment in the product documentation would be
#    appreciated but is not required.
# 2. Altered source versions must be plainly marked as such, and must not be
#    misrepresented as being the original software.
# 3. This notice may not be removed or altered from any source distribution.

from dataclasses import dataclass
import os.path
import struct
import sys
import threading
import math
from lighting_server_gui import MainPage
from bgapi.bglib import CommandFailedError

sys.path.append(os.path.join(os.path.dirname(__file__), "../.."))
from common.util import BtMeshApp
from common import status
import common.btmesh_models as model

IMMEDIATE = 0
STATE_OFF = 0
STATE_ON = 1

class OnOffServer(BtMeshApp):
    """" Implementation of Generic On/Off Server. """
    def __init__(self, connector, **kwargs):
        super().__init__(connector=connector, **kwargs)
        # Copy of transition delay parameter, needed for delayed on/off request
        self.delayed_onoff_trans = 0
        # NVM save time
        self.nvm_save_time = 5000
        # Lighting server key to store data to nvm
        self.lighting_server_key = 0x4004
        # NVM save timer
        self.lighting_nvm_save_timer = threading.Timer(self.nvm_save_time * 0.001,
                                                       self.light_lightbulb_state_changed)
        # Current primary generic level value
        self.pri_level_current = -32768
        # Target primary generic level value
        self.pri_level_target = -32768
    @dataclass
    class CTLLightbulbLightness:
        """ Dataclass of lightbulb lightness state. """
        # Current lightness value
        lightness_current = 0
        # Target lightness value
        lightness_target = 0
        # Last lightness value
        lightness_last = 0xFFFF
        # Default lightness value
        lightness_default = 0x0000
        # Minimum lightness value
        lightness_min = 0x0001
        # Maximum lightness value
        lightness_max = 0xFFFF 
        # Minimum brightness
        min_brightness = 0

    @dataclass
    class CTLLightbulbOnOff:
        """ Dataclass of lightbulb on/off state. """
        # Current generic on/off value
        onoff_current = 0
        # Target generic on/off value
        onoff_target = 0
        # Transition time
        transtime = 0
        # On Power Up value
        onpowerup = 0


    def onoff_request(self, event):
        """ Process the requests for the generic on/off model. """
        request_state = int.from_bytes(event.parameters, byteorder="little")
        self.log.info(f"ON/OFF request: requested state = {request_state}, " +
                      f"trans = {event.transition_ms}, "+
                      f"delay= {event.delay_ms}")

        if self.CTLLightbulbOnOff.onoff_current == request_state:
            self.log.info("Request for current state received.")
        else:
            onoff_message = "ON" if request_state == 1 else "OFF"
            self.log.info(f"Turning lightbulb {onoff_message}")
            # Set target value
            self.CTLLightbulbOnOff.onoff_target = request_state
            # Immediate change
            if event.transition_ms == 0 and event.delay_ms == 0:
                self.CTLLightbulbOnOff.onoff_current = request_state

                if self.CTLLightbulbOnOff.onoff_current == STATE_OFF:
                    self.CTLLightbulbLightness.lightness_target = 0
                    self.CTLLightbulbLightness.lightness_current = (
                        self.CTLLightbulbLightness.lightness_target)
                else:
                    # Restore last brightness
                    self.CTLLightbulbLightness.lightness_target = (
                        self.CTLLightbulbLightness.lightness_last)
                    self.CTLLightbulbLightness.lightness_current = (
                        self.CTLLightbulbLightness.lightness_target)

                self.lighting_set_level(self.CTLLightbulbLightness.lightness_target,
                                        IMMEDIATE)

            elif event.delay_ms > 0:
                self.delayed_onoff_trans = event.transition_ms
                # Timer start
                threading.Timer(event.delay_ms * 0.001,
                                self.delayed_onoff_request).start()

            else:
                # No delay but transition time has been set.
                if self.CTLLightbulbOnOff.onoff_target == STATE_ON:
                    self.CTLLightbulbOnOff.onoff_current = STATE_ON

                self.onoff_update(event.elem_index, event.transition_ms)

                if request_state == STATE_OFF:
                    self.CTLLightbulbLightness.lightness_target = 0

                else:
                    # Restore last brightness
                    self.CTLLightbulbLightness.lightness_target = (
                        self.CTLLightbulbLightness.lightness_last)

                self.lighting_set_level(self.CTLLightbulbLightness.lightness_target,
                                        event.transition_ms)
                # Lightbulb current state will be updated when transition is complete
                threading.Timer(event.transition_ms * 0.001,
                                self.onoff_transition_complete).start()

            # Save the state in flash after a small delay
            self.lighting_nvm_save_timer_start()
            # State has changed, so the current scene number is reset
            self.lib.btmesh.scene_server.reset_register(event.elem_index)

        remaining_ms = event.delay_ms + event.transition_ms
        if event.flags & 2:
            # Response required. If non-zero, the client expects a response from the server.
            self.onoff_response(event.elem_index,
                                event.client_address,
                                event.appkey_index,
                                remaining_ms)

        self.onoff_update_and_publish(event.elem_index, remaining_ms)

    def delayed_onoff_request(self):
        """ Handle delayed light on/off requests. """
        self.log.info("Starting delayed ON/OFF request:" +
                      f"{self.CTLLightbulbOnOff.onoff_current} ->" +
                      f"{self.CTLLightbulbOnOff.onoff_target},"
                      f"with {self.delayed_onoff_trans} ms transition")

        if self.delayed_onoff_trans == 0:
            # No transition delay, update state immediately
            self.CTLLightbulbOnOff.onoff_current = (
                self.CTLLightbulbOnOff.onoff_target)
            if self.CTLLightbulbOnOff.onoff_current == STATE_OFF:
                self.lighting_set_level(self.min_brightness, self.delayed_onoff_trans)
            else:
                # Restore last brightness level
                self.lighting_set_level(self.CTLLightbulbLightness.lightness_last,
                                        IMMEDIATE)
                self.CTLLightbulbLightness.lightness_current = (
                    self.CTLLightbulbLightness.lightness_last)
                self.CTLLightbulbLightness.lightness_target = (
                    self.CTLLightbulbLightness.lightness_last)

            # Save the state in flash after a small delay
            self.lighting_nvm_save_timer_start()
            self.onoff_update_and_publish(0, self.delayed_onoff_trans)

        else:
            # Delay and transition time is greather than 0
            if self.CTLLightbulbOnOff.onoff_target == STATE_OFF:
                self.CTLLightbulbLightness.lightness_target = 0
            else:
                self.CTLLightbulbLightness.lightness_target = (
                    self.CTLLightbulbLightness.lightness_last)
                self.CTLLightbulbOnOff.onoff_current = STATE_ON

                self.onoff_update(0, self.delayed_onoff_trans)

            self.lighting_set_level(self.CTLLightbulbLightness.lightness_current,
                                    self.delayed_onoff_trans,)
            # State is updated when transition is complete
            threading.Timer(self.delayed_onoff_trans * 0.001,
                            self.onoff_transition_complete).start()

    def onoff_update(self, elem_index, remaining_ms):
        """ Update generic on/off state. """
        onoff = struct.pack("<BB",
                            self.CTLLightbulbOnOff.onoff_current,
                            self.CTLLightbulbOnOff.onoff_target)

        self.lib.btmesh.generic_server.update(
            elem_index,
            model.BTMESH_GENERIC_ON_OFF_SERVER_MODEL_ID,
            remaining_ms,
            self.lib.btmesh.generic_client.GET_STATE_TYPE_STATE_ON_OFF,
            onoff,
        )

    def onoff_response(self, elem_index, client_addr, appkey_index, remaining_ms):
        """ Respond to generic on/off requests. """
        self.log.info("Response sent")
        onoff = struct.pack("<BB",
                            self.CTLLightbulbOnOff.onoff_current,
                            self.CTLLightbulbOnOff.onoff_target)

        self.lib.btmesh.generic_server.respond(
            client_addr,
            elem_index,  # 0
            model.BTMESH_GENERIC_ON_OFF_SERVER_MODEL_ID,
            appkey_index,
            remaining_ms,
            0x00,
            self.lib.btmesh.generic_client.GET_STATE_TYPE_STATE_ON_OFF,
            onoff,
        )

    def onoff_update_and_publish(self, elem_index, remaining_ms):
        """ Update onoff state and publish model state to the network. """
        self.onoff_update(elem_index, remaining_ms)

        try:
            self.lib.btmesh.generic_server.publish(
                elem_index,  # 0
                model.BTMESH_GENERIC_ON_OFF_SERVER_MODEL_ID,
                self.lib.btmesh.generic_client.GET_STATE_TYPE_STATE_ON_OFF,
            )
        except CommandFailedError as e:
            # Application key or publish address are not set
            if e.errorcode != status.BT_MESH_PUBLISH_NOT_CONFIGURED:
                raise

    def onoff_transition_complete(self):
        """ Callback to light on/off request with non-zero transition time. """
        # Transition done -> set state, update and publish
        self.CTLLightbulbOnOff.onoff_current = (
            self.CTLLightbulbOnOff.onoff_target)
        onoff_message = (
            "ON" if self.CTLLightbulbOnOff.onoff_current == 1 else "OFF")
        self.log.info(f"{onoff_message}")
        # Save the state in flash after a small delay
        self.lighting_nvm_save_timer_start()
        self.onoff_update_and_publish(0, IMMEDIATE)

    def onoff_recall(self, event):
        """ Handle generic on/off recall events. """
        request_state = int.from_bytes(event.parameters, byteorder="little")
        if (self.CTLLightbulbOnOff.onoff_current
            == self.CTLLightbulbOnOff.onoff_target):
            self.log.info("Request for current OnOff state received.")
        else:
            onoff_message = "ON" if request_state == 1 else "OFF"
            self.log.info(f"Turning lightbulb {onoff_message}")

            self.CTLLightbulbOnOff.onoff_target = request_state
            if event.transition_time_ms == IMMEDIATE:
                self.CTLLightbulbOnOff.onoff_current = (
                    self.CTLLightbulbOnOff.onoff_target)
            else:
                if self.CTLLightbulbOnOff.onoff_target == STATE_ON:
                    self.CTLLightbulbOnOff.onoff_current = STATE_ON

                # Lightbulb current state will be updated when transition is complete
                threading.Timer(event.transition_time_ms * 0.001,
                                self.onoff_transition_complete).start()

            # Save the state in flash after a small delay
            self.lighting_nvm_save_timer_start()

        self.onoff_update_and_publish(event.elem_index, event.transition_time_ms)

    def onoff_change(self, event):
        """ Handle generic on/off change events. """
        length = len(event.parameters)
        param = event.parameters
        if length >= 1:
            current = param[0]

        if current != self.CTLLightbulbOnOff.onoff_current:
            self.log.info(f"On/Off state changed " +
                          f"from {self.CTLLightbulbOnOff.onoff_current} " +
                          f"to {current}")
            self.CTLLightbulbOnOff.onoff_current = current
            self.lighting_nvm_save_timer_start()
        else:
            self.log.info("On/Off change - same state as before")

    def lighting_set_level(self, level, trans_ms):
        """ Set GUI lightness level in given transition time. """
        self.lightness_current = level
        temp = MainPage.get_instance(self).CTLLightbulbState.temperature_value.get()
        temperature = self.temperature_to_rgb(temp)
        color = self.rgb_to_lightnessrgb(temperature, self.CTLLightbulbLightness.lightness_current)
        MainPage.get_instance(self).set_lightness_value(
            self.actual_value_to_precentage(self.CTLLightbulbLightness.lightness_current)
        )
        MainPage.get_instance(self).fade(
            MainPage.get_instance(self).lightbulb_image, color, trans_ms
        )

    ##### HELPER FUNCTIONS #####

    def actual_value_to_precentage(self, value):
        """ Convert actual lightness value to percentage. """
        return round((value * 100) / 0xFFFF)

    def temperature_to_rgb(self, temperature):
        """ Convert temperature to RGB color. """
        temp_red = 0
        temp_green = 0
        temp_blue = 0

        if temperature < 6563:
            temp_red = 255
            if temperature < 1925:
                temp_blue = 0
            else:
                temp_blue = temperature - 1918.74282
                temp_blue = 2.55822107 * pow(temp_blue, 0.546877914)

            if temperature < 909:
                temp_green = 0
            else:
                temp_green = temperature - 636.62578769
                temp_green = 73.13384712 * math.log(temp_green) - 383.76244858

        else:
            temp_red = temperature - 5882.02392431
            temp_red = -29.28670147 * math.log(temp_red) + 450.50427359
            temp_red = temp_red + 0.5
            temp_green = temperature - 5746.13180276
            temp_green = -18.69512921 * math.log(temp_green) + 377.39334366
            temp_blue = 255

        temp_max = max(temp_red, temp_green, temp_blue)

        temp_red = temp_red * 255 / temp_max
        temp_green = temp_green * 255 / temp_max
        temp_blue = temp_blue * 255 / temp_max

        color_red = 255 if temp_red > 255 else (round(temp_red) if temp_red >= 0 else 0)
        color_green = (255 if temp_green > 255 else (round(temp_green) if temp_green >= 0 else 0))
        color_blue = (255 if temp_blue > 255 else (round(temp_blue) if temp_blue >= 0 else 0))

        rgb = {"R": color_red, "G": color_green, "B": color_blue}

        return rgb

    def rgb_to_lightnessrgb(self, color, level):
        """ Change lightness of given color temperature. """
        new_color_red = round((color["R"] * level) / 65535)
        new_color_green = round((color["G"] * level) / 65535)
        new_color_blue = round((color["B"] * level) / 65535)

        return ("#"
                + f"{new_color_red:02X}"
                + f"{new_color_green:02X}"
                + f"{new_color_blue:02X}")

    def light_lightbulb_state_changed(self):
        """ Save the state to the nvm after the lighting lightbulb state is changed. """
        self.lib.bt.nvm.save(self.lighting_server_key, self.lightbulb_state_serialize())

    def lighting_nvm_save_timer_start(self):
        """ Start or restart lighting NVM save timer. """
        if self.lighting_nvm_save_timer.is_alive():
            self.lighting_nvm_save_timer.cancel()
        self.lighting_nvm_save_timer = threading.Timer(self.nvm_save_time * 0.001,
                                                       self.light_lightbulb_state_changed)
        self.lighting_nvm_save_timer.start()

    def lightbulb_state_serialize(self):
        """ Pack dataclass to a struct for NVM save. """
        lightbulb_state = struct.pack(
            "<BBHHHHHHHHhh",
            self.CTLLightbulbOnOff.onoff_current,
            self.CTLLightbulbOnOff.onoff_target,
            self.CTLLightbulbOnOff.transtime,
            self.CTLLightbulbOnOff.onpowerup,
            self.CTLLightbulbLightness.lightness_current,
            self.CTLLightbulbLightness.lightness_target,
            self.CTLLightbulbLightness.lightness_last,
            self.CTLLightbulbLightness.lightness_default,
            self.CTLLightbulbLightness.lightness_min,
            self.CTLLightbulbLightness.lightness_max,
            self.pri_level_current,
            self.pri_level_target
        )
        return lightbulb_state
```
