## Supplementary material for "VesperFlow: An automated pipeline for decoding multisensor data from biologging tags": README.md

# Bluetooth Mesh - NCP Host Light (Lighting Server)

This example demonstrates the operation of a Bluetooth Mesh Lighting Server (Light). 
The graphical user interface (GUI) simulates the behevior of a lightbulb that can be controlled by
another device implementing the client role (for example Bluetooth Mesh - SoC Switch). The 
virtual lightbulb can be switched on/off, and the lighting intensity, color temperature, and 
the Delta UV can also be set.

The following BT Mesh Models are demonstrated:
 * Generic OnOff Server
 * Generic Level Server
 * Light Lightness Server
 * Light CTL Server
 * Scene Server

## Getting Started

To learn Bluetooth Mesh technology fundamentals, see [Bluetooth Mesh Network - An Introduction for Developers](https://www.bluetooth.com/wp-content/uploads/2019/03/Mesh-Technology-Overview.pdf).

To get to know the Bluetooth Mesh SDK and Simplicity Studio, see [ QSG176: Bluetooth® Mesh SDK v2.x Quick Start Guide](https://www.silabs.com/documents/public/quick-start-guides/qsg176-bluetooth-mesh-sdk-v2x-quick-start-guide.pdf).

### Follow this procedure to get started

1. Install pyBGAPI.
	```
    pip install pybgapi
    ```
2. Clone the [repo](https://github.com/SiliconLabs/pybgapi-examples) from GitHub and check out the master barnch.
3. Install the 'colour library' and 'pillow library' python packages
	```
	pip install colour
	```
	and
	```
	pip install pillow
	```
4. Flash the **Bluetooth Mesh - NCP Empty** demo binary on your [Wireless Development Kit](https://www.silabs.com/development-tools/wireless)
    using [Simplicity Studio](https://docs.silabs.com/simplicity-studio-5-users-guide/latest/ss-5-users-guide-building-and-flashing/flashing).
5. Run the btmesh-light example from the repo.
	```
	python3 example/btmesh_light/app.py
	```
6. Provision the light example into the mesh network with a Provisioner device.
	Recommended option:<br>
	Silicon Labs Bluetooth Mesh smartphone application ([ QSG176: Bluetooth® Mesh SDK v2.x Quick Start Guide](https://www.silabs.com/documents/public/quick-start-guides/qsg176-bluetooth-mesh-sdk-v2x-quick-start-guide.pdf)).

	Other options:
	 * Bluetooth NCP Commander within Simplicity Studio
	 * Host Provisioner example in Bluetooth Mesh SDK.
7. Make sure that a device implementing the Client role is in the mesh network.
	Recommended option: pyBGAPI switch example ([Bluetooth mesh - Switch](example/btmesh_switch))

	Other options:
	 * Bluetooth Mesh - SoC Switch in Bluetooth Mesh SDK
	 *  Silicon Labs Bluetooth Mesh smartphone application
8.  How to use:
	* Use the switch app's GUI to switch on the lightbulb

	![OnOff](images/OnOff.png)
	
	* Use the switch app's Lightness slider to change the lightbulb lightness

	![Lightness](images/Lightness.png)

	* Use the switch app's Temperature slider to change the color temperature

	![CTL](images/CTL.png)

## Structure of the Python Application

![Class](images/class.png)

- OnOffServer class: this class implements the Generic OnOff Server model.<br>
The class includes a dataclass that represents the lightbulb state. It also handles onoff requests, delayed onoff requests, onoff recalls and onoff changes from the stack or a remote device.
After a change the class can send onoff responses and onoff updates, and can update the GUI.

- LightnessServer class: this class implements the Generic Level Server model and the Light Lightness Server model and extends the OnOffServer class.<br>
It handles generic level (pri_level) and lightness requests, delayed generic level (pri_level) and delayed lightness requests, generic level (pri_level) and lightness recalls, generic level (pri_level) and lightness changes from the stack or a remote device. After a change the class can send generic level (pri_level) and lightness responses, lightness updates, and can update the GUI.

- CTLServer class: the class implements the Light CTL Server model and extends the LightnessServer class.<br>
The class includes a dataclass that represents the CTL lightbulb state. 
It handles ctl requests, delayed ctl requests, ctl recalls and ctl changes from the stack or a remote device.
After a change the class can send ctl responses and ctl updates, and can update the GUI.

- FactoryReset class: this class extends the BtMeshApp class and implements the node reset and full factory reset functions.

- App class: the class extends the CTLServer and the FactoryReset class.
The main fuction of this class is to handle the received events and forward them to the right class for processing.
It restores the data from NVM and loads the data to the given dataclass.
After running the application two threads are created. One of the threads is responsible for the BT Mesh event handling while the other thread is responsible for the GUI.

- windows and MainPage class: this class extends the Tk class (tkinter library).
The windows class is a general class for making tkinter windows, where you can add the name of the window, the size of the window, and so on.
The MainPage class defines the actual layout of the window, variables and functions.

The example extends the BtMeshApp class. The BtMeshApp class implements the following: Check BGAPI Version, Initialize Mesh Node,
Factory and Node Reset. It uses both Bluetooth and Bluetooth Mesh XAPI files.
