## Supplementary material for "VesperFlow: An automated pipeline for decoding multisensor data from biologging tags": app.py

```
"""
BtMesh Switch NCP-host Example Application.
"""

# Copyright 2022 Silicon Laboratories Inc. www.silabs.com
#
# SPDX-License-Identifier: Zlib
#
# The licensor of this software is Silicon Laboratories Inc.
#
# This software is provided 'as-is', without any express or implied
# warranty. In no event will the authors be held liable for any damages
# arising from the use of this software.
#
# Permission is granted to anyone to use this software for any purpose,
# including commercial applications, and to alter it and redistribute it
# freely, subject to the following restrictions:
#
# 1. The origin of this software must not be misrepresented; you must not
#    claim that you wrote the original software. If you use this software
#    in a product, an acknowledgment in the product documentation would be
#    appreciated but is not required.
# 2. Altered source versions must be plainly marked as such, and must not be
#    misrepresented as being the original software.
# 3. This notice may not be removed or altered from any source distribution.

import os
import sys
import threading
import switch_gui

# Btmesh model classes and the general class
from onoff_client import OnOffClient
from lightness_client import LightnessClient
from ctl_client import CTLClient
from scene_client import SceneClient
from reset import Reset
from lpn import LPN
sys.path.append(os.path.join(os.path.dirname(__file__), "../.."))
from common.util import ArgumentParser, get_connector

# Advertising options
PB_ADV = 0x1
PB_GATT = 0x2

# Characteristic values
GATTDB_DEVICE_NAME = b"BtMesh Switch Example"
GATTDB_MANUFACTURER_NAME_STRING = b"Silicon Labs"

class App(OnOffClient, LightnessClient, CTLClient, Reset, SceneClient, LPN):
    """ 
    Application derived from OnOffClient, LightnessClient, 
    CTLClient, General, SceneClient, LPN. 
    """
    def node_reset(self):
        self.lpn_feature_deinit()
        return super().node_reset()

    def factory_reset(self):
        self.lpn_feature_deinit()
        return super().factory_reset()

    # Common event callbacks
    def btmesh_evt_node_initialized(self, evt):
        """ Bluetooth mesh event callback """
        if evt.provisioned:
            self.log.info("Node initialized and provisioned")
            self.lpn_feature_init()
            self.log.info("LPN feature initialized")
        else:
            self.lib.btmesh.node.start_unprov_beaconing(PB_ADV | PB_GATT)
            self.log.info("Node is ready to be provisioned")

        self.lib.btmesh.generic_client.init()
        self.log.info("All generic client models initialized")
        # Scene model setup and init
        self.lib.btmesh.scene_client.init(0)

    def btmesh_evt_node_provisioned(self, _evt):
        """ Bluetooth mesh event callback """
        self.set_configuraton_timer(self.LPNTimeout.lpn_timeout_after_provisioned)

    def btmesh_evt_node_model_config_changed(self, _evt):
        """ Bluetooth mesh event callback """
        self.set_configuraton_timer(self.LPNTimeout.lpn_timeout_after_config_model_changed)

    def btmesh_evt_node_config_set(self, _evt):
        """ Bluetooth mesh event callback """
        self.set_configuraton_timer(self.LPNTimeout.lpn_timeout_after_config_set)

    def btmesh_evt_node_key_added(self, _evt):
        """ Bluetooth mesh event callback """
        self.set_configuraton_timer(self.LPNTimeout.lpn_timeout_after_key)

    def btmesh_evt_lpn_friendship_established(self, evt):
        """ Bluetooth mesh event callback """
        self.log.info(f"Friendship established. Friend address = {evt.friend_address}")

    def btmesh_evt_lpn_friendship_terminated(self, evt):
        """ Bluetooth mesh event callback """
        self.log.info(f"Friendship terminated. Reason = {evt.reason}")
        if self.num_mesh_proxy_conn == 0:
            self.lpn_friend_find_timer(self.LPNTimeout.lpn_friend_find_timeout)

    def btmesh_evt_lpn_friendship_failed(self, evt):
        """ Bluetooth mesh event callback """
        self.log.info(f"Friendship failed. Reason = {evt.reason}")
        self.lpn_friend_find_timer(self.LPNTimeout.lpn_friend_find_timeout)

    def btmesh_evt_proxy_connected(self, _evt):
        """ Bluetooth mesh event callback """
        self.log.info("Proxy connected")
        self.num_mesh_proxy_conn += 1
        # Turn off LPN feature after GATT proxy connection is opened.
        self.lpn_feature_deinit()

    def btmesh_evt_proxy_disconnected(self, _evt):
        """ Bluetooth mesh event callback """
        self.log.info("Proxy disconnected")
        if self.num_mesh_proxy_conn > 0:
            self.num_mesh_proxy_conn -= 1
            if self.num_mesh_proxy_conn == 0:
                # Initialize lpn when there is no active proxy connection.
                self.lpn_feature_init()

        ####################################
        # Add further event handlers here.
        ####################################

# Script entry point.
if __name__ == "__main__":
    # Instantiate and run application.
    parser = ArgumentParser(description=__doc__)
    args = parser.parse_args()
    connector = get_connector(args)
    app = App(connector=connector)
    t_app = threading.Thread(target=app.run, args=[], daemon=True)
    t_app.name = "App_thread"
    t_app.start()
    switch_gui.switch_gui_start(app)
```
