## Supplementary material for "VesperFlow: An automated pipeline for decoding multisensor data from biologging tags": ctl_client.py

```
"""
BtMesh Switch NCP-host Light CTL Client Model.
"""

# Copyright 2022 Silicon Laboratories Inc. www.silabs.com
#
# SPDX-License-Identifier: Zlib
#
# The licensor of this software is Silicon Laboratories Inc.
#
# This software is provided 'as-is', without any express or implied
# warranty. In no event will the authors be held liable for any damages
# arising from the use of this software.
#
# Permission is granted to anyone to use this software for any purpose,
# including commercial applications, and to alter it and redistribute it
# freely, subject to the following restrictions:
#
# 1. The origin of this software must not be misrepresented; you must not
#    claim that you wrote the original software. If you use this software
#    in a product, an acknowledgment in the product documentation would be
#    appreciated but is not required.
# 2. Altered source versions must be plainly marked as such, and must not be
#    misrepresented as being the original software.
# 3. This notice may not be removed or altered from any source distribution.

from dataclasses import dataclass
import os
import sys
import threading
sys.path.append(os.path.join(os.path.dirname(__file__), "../.."))
from common.util import BtMeshApp
from common import btmesh_models as model

# Immediate transition time is 0 seconds
IMMEDIATE = 0
# No flags used for message
NO_FLAGS = 0

class CTLClient(BtMeshApp):
    """Implement the Light CTL Client Model specific APIs."""
    def __init__(self, connector, **kwargs):
        super().__init__(connector=connector, **kwargs)
        # ctl transaction identifier
        self.ctl_trid = 0
        # Delay time (in milliseconds) before starting the state change
        self.request_delay = 50
        # Lightness level converted from percentage to actual value, range 0..65535
        self.ctl_lightness = 0
        # Maximum lightness percentage value
        self.lightness_pct_max = 100
        # How many times CTL model messages are to be sent out for reliability
        # Using three by default
        self.ctl_request_count = 3

    @dataclass
    class Temperature:
        """ Dataclass of temperature values. """
        # Current temperature value
        min = 800
        # Target temperature value
        max = 20000
        # Default temperature value
        level = 800

    @dataclass
    class DeltaUV:
        """ Dataclass of deltaUV values. """
        # Delta UV, default value is 0
        value = 0
        # Maximum Delta UV value
        max = 1
        # Minimum Delta UV value
        min = -1

    def send_light_ctl_request(self):
        """
        Call 'ctl_request' with 'request_delay' 
        intervals for 'ctl_request_count' times.
        """
        # Increment transaction ID for each request, unless it's a retransmission.
        self.ctl_trid += 1
        self.ctl_trid %= 256

        # Starting two new timer threads for the second and third message.
        for count in range(1, self.ctl_request_count, 1):
            threading.Timer(count * self.request_delay * 0.001,
                            self.ctl_request,
                            args=[self.ctl_request_count - count]).start()

        # First message with 0ms delay
        self.ctl_request(self.ctl_request_count)

    def ctl_request(self, count):
        """
        Serialize and publish CTL requests, and calculate the delay for them.

        :param temperature: holds the latest desired temperature value
        :param count: number of repetition
        """
        delay = (count - 1) * self.request_delay
        if count > 0:
            self.lib.btmesh.generic_client.publish(
                0,
                model.BTMESH_LIGHTING_CTL_CLIENT_MODEL_ID,
                self.ctl_trid,
                IMMEDIATE,
                delay,
                NO_FLAGS,
                self.lib.btmesh.generic_client.SET_REQUEST_TYPE_REQUEST_CTL,
                self.serialize(self.ctl_lightness,
                               self.Temperature.level,
                               self.DeltaUV.value)
            )

            self.log.info(f"Ctl request, trid: {self.ctl_trid}, delay: {delay}")

    def serialize(self, lightness, temperature, delta_uv):
        """ Serialize lightness, temperature and Delta UV values. """
        bytes_lightness = lightness.to_bytes(2, byteorder='little')
        bytes_temperature = temperature.to_bytes(2, byteorder='little')
        bytes_deltauv = delta_uv.to_bytes(2, byteorder='little')
        return bytes_lightness + bytes_temperature + bytes_deltauv

    def convert_lightness_percentage(self, lightness):
        """
        Converts the lightness from percentage to actual value,
        rage 0..65535.

        :param lightness: lightness in percentage
        """
        return (lightness * 0xFFFF) // self.lightness_pct_max

    def convert_delta_uv(self, delta_uv):
        """
        Convert Delta UV value from a scale of -1 to 1 to a 16 bit signed interger.

        :param delta_uv: Delta UV value in float
        """
        temp = int(delta_uv * 32768)
        if temp > 32767:
            temp = 32767 # 0x7FFF
        elif temp < -32768:
            temp = -32768  # -0x8000

        return temp & 0xFFFF

    def set_temperature(self, set_lightness, set_temperature):
        """
        Check if the given value is valid. Valid value range is between
        'temperature_min' and 'temperature_max'. If the value is not valid
        then adjust it to be a proper one and call the ctl request function.

        :param set_lightness: the current lightness value
        :param set_temperature: desired temperature state given by the user
        """
        if set_temperature > self.Temperature.max:
            self.Temperature.level = self.Temperature.max
        elif set_temperature < self.Temperature.min:
            self.Temperature.level = self.Temperature.min
        else:
            self.Temperature.level = set_temperature

        self.ctl_lightness = self.convert_lightness_percentage(set_lightness)
        self.send_light_ctl_request()

    def set_delta_uv(self, set_lightness, set_delta_uv):
        """
        Set the converted velues of Delta UV and lightness
        and call the ctl request function.

        :param set_lightness: the current lightness value
        :param set_delta_uv: desired Delta UV state given by the user
        """
        self.DeltaUV.value = self.convert_delta_uv(set_delta_uv)
        self.ctl_lightness = self.convert_lightness_percentage(set_lightness)
        self.send_light_ctl_request()
```
