## Supplementary material for "VesperFlow: An automated pipeline for decoding multisensor data from biologging tags": lightness_client.py

```
"""
BtMesh Switch NCP-host Light Lightness Client Model.
"""

# Copyright 2022 Silicon Laboratories Inc. www.silabs.com
#
# SPDX-License-Identifier: Zlib
#
# The licensor of this software is Silicon Laboratories Inc.
#
# This software is provided 'as-is', without any express or implied
# warranty. In no event will the authors be held liable for any damages
# arising from the use of this software.
#
# Permission is granted to anyone to use this software for any purpose,
# including commercial applications, and to alter it and redistribute it
# freely, subject to the following restrictions:
#
# 1. The origin of this software must not be misrepresented; you must not
#    claim that you wrote the original software. If you use this software
#    in a product, an acknowledgment in the product documentation would be
#    appreciated but is not required.
# 2. Altered source versions must be plainly marked as such, and must not be
#    misrepresented as being the original software.
# 3. This notice may not be removed or altered from any source distribution.

from dataclasses import dataclass
import os
import sys
import threading
sys.path.append(os.path.join(os.path.dirname(__file__), "../.."))
from common.util import BtMeshApp
import common.btmesh_models as model

# Immediate transition time is 0 seconds
IMMEDIATE = 0
# No flags used for message
NO_FLAGS = 0

class LightnessClient(BtMeshApp):
    """ Implementation of the Light Lightness Client Model specific APIs. """
    def __init__(self, connector, **kwargs):
        super().__init__(connector=connector, **kwargs)
        # Lightness transaction identifier
        self.lightness_trid = 0
        # Delay time (in milliseconds) before starting the state change
        self.request_delay = 50
        # Transition time (in milliseconds) for the state change
        # Using zero transition time by default
        self.transtime = 0
        # How many times Lightness model messages are to be sent out for reliability
        # Using three by default
        self.lightness_request_count = 3

    @dataclass
    class Lightness:
        """ Dataclass of lightness values. """
        # Lightness level converted from percentage to actual value, range 0..65535
        lightness_level = 0
        # Lightness level percentage
        lightness_percentage = 0
        # Maximum lightness percentage value
        lightness_pct_max = 100

    def send_lightness_actual_request(self):
        """
        This function publishes light lightness request to change the lightness
        level of light(s) in the group. The lightness_level variable holds
        the latest desired light level.
        """
        # Increment transaction ID for each request, unless it's a retransmission.
        self.lightness_trid += 1
        self.lightness_trid %= 256

        # Starting two new timer threads for the second and third message.
        for count in range(1, self.lightness_request_count, 1):
            threading.Timer(count * self.request_delay * 0.001,
                            self.lightness_actual_request,
                            args=[self.Lightness.lightness_level,
                                  self.lightness_request_count - count]).start()

        # First message with 0ms delay
        self.lightness_actual_request(self.Lightness.lightness_level,
                                      self.lightness_request_count)

    def lightness_actual_request(self, lightness, count):
        """
        This function serializes Light Lightness request and calculates the delay for it.

        :param lightness: holds the latest desired lightness state in percentage
        :param count: number of repetition
        """
        delay = (count - 1) * self.request_delay
        if count > 0:
            self.lib.btmesh.generic_client.publish(
                0,
                model.BTMESH_LIGHTING_LIGHTNESS_CLIENT_MODEL_ID,
                self.lightness_trid,
                IMMEDIATE,
                delay,
                NO_FLAGS,
                self.lib.btmesh.generic_client.SET_REQUEST_TYPE_REQUEST_LIGHTNESS_ACTUAL,
                lightness.to_bytes(2, byteorder="little"),
            )

            self.log.info(f"Lightness actual request, trid: {self.lightness_trid}, delay: {delay}")

    def convert_lightness_percentage(self, lightness):
        """
        This function converts the lightness from percentage to actual value,
        rage 0..65535.

        :param lightness: lightness in percentage
        """
        return (lightness * 0xFFFF) // self.Lightness.lightness_pct_max

    def set_lightness(self, set_percentage):
        """
        This function checks if the given value is valid. Valid values are between 0 and 100.
        If the value is not valid than the function adjusts it to be a proper value.

        :param set_percentage: desired lightness state given by the user
        """
        if set_percentage > self.Lightness.lightness_pct_max:
            self.Lightness.lightness_percentage = self.Lightness.lightness_pct_max
        elif set_percentage < 0:
            self.Lightness.lightness_percentage = 0
        else:
            self.Lightness.lightness_percentage = set_percentage

        self.Lightness.lightness_level = self.convert_lightness_percentage(set_percentage)
        self.send_lightness_actual_request()
```
