## Supplementary material for "VesperFlow: An automated pipeline for decoding multisensor data from biologging tags": lpn.py

```
"""
BtMesh Switch Low Power Node implementation.
"""

# Copyright 2022 Silicon Laboratories Inc. www.silabs.com
#
# SPDX-License-Identifier: Zlib
#
# The licensor of this software is Silicon Laboratories Inc.
#
# This software is provided 'as-is', without any express or implied
# warranty. In no event will the authors be held liable for any damages
# arising from the use of this software.
#
# Permission is granted to anyone to use this software for any purpose,
# including commercial applications, and to alter it and redistribute it
# freely, subject to the following restrictions:
#
# 1. The origin of this software must not be misrepresented; you must not
#    claim that you wrote the original software. If you use this software
#    in a product, an acknowledgment in the product documentation would be
#    appreciated but is not required.
# 2. Altered source versions must be plainly marked as such, and must not be
#    misrepresented as being the original software.
# 3. This notice may not be removed or altered from any source distribution.

from dataclasses import dataclass
import os
import sys
import threading
import bgapi.bglib
sys.path.append(os.path.join(os.path.dirname(__file__), "../.."))
from common.util import BtMeshApp

class LPN(BtMeshApp):
    """ Implementation of the LPN node functionalities. """
    def __init__(self, connector, **kwargs):
        super().__init__(connector=connector, **kwargs)
        # Flag for indicating that LPN feature is active
        self.lpn_active = 0
        # Flag for proxy connection
        self.num_mesh_proxy_conn = 0
        # Minimum queue length the friend must support (2-128)
        self.min_queue_length = 2
        # Receive delay in milliseconds (10-255)
        self.received_delay = 50
        # The number of retry attempts to repeat (0-10)
        self.request_retries = 8
        # Time interval between retry attempts in milliseconds (0-100)
        self.retry_interval = 100
        # List of network IDs.
        self.lpn_friend_netkey_idx = 0
    @dataclass
    class LPNTimeout:
        """ Dataclass of LPN timeout. """
        # Poll timeout in milliseconds (1000-345599900:100)
        poll_timeout = 5000
        # Timeout for initializing LPN after an already provisioned node is initialized
        lpn_timeout_after_provisioned = 30000
        # Timeout for initializing LPN after the Configuration Model changed
        lpn_timeout_after_config_model_changed = 5000
        # Timeout for initializing LPN after the Configuration Model Set Message
        lpn_timeout_after_config_set = 5000
        # Timeout for initializing LPN after Security Key was added
        lpn_timeout_after_key = 5000
        # Timeout between retries to find a friend
        lpn_friend_find_timeout = 30000
        # Friendship timer for timeouts
        friendship_timer = None

    # Key values to identify LPN configurations
    lpn_conf ={
        'lpn_queue_length':0,
        'lpn_poll_timeout':1,
        'lpn_receive_delay':2,
        'lpn_request_retries':3,
        'lpn_retry_interval':4,
        'lpn_clock_accuracy':5
    }

    def lpn_friend_find_timer(self, delay_ms):
        """ Try to find a friend after the time expires. """
        self.LPNTimeout.friendship_timer = threading.Timer(delay_ms * 0.001,
                                                           self.lpn_establish_friendship,
                                                           args=[self.lpn_friend_netkey_idx])
        self.LPNTimeout.friendship_timer.start()

    def lpn_feature_init(self):
        """ Initialize LPN functionality with configuration and friendship establishment. """
        # Do not initialize LPN if LPN is currently active or any GATT proxy connection is open.
        if self.lpn_active or self.num_mesh_proxy_conn:
            return

        # Initialize LPN functionality.
        self.lib.btmesh.lpn.init()
        self.lpn_active = 1

        # LPN configurations.
        self.lib.btmesh.lpn.config(self.lpn_conf["lpn_queue_length"],
                                   self.min_queue_length)
        self.lib.btmesh.lpn.config(self.lpn_conf["lpn_poll_timeout"],
                                   self.LPNTimeout.poll_timeout)
        self.lib.btmesh.lpn.config(self.lpn_conf["lpn_receive_delay"],
                                   self.received_delay)
        self.lib.btmesh.lpn.config(self.lpn_conf["lpn_request_retries"],
                                   self.request_retries)
        self.lib.btmesh.lpn.config(self.lpn_conf["lpn_retry_interval"],
                                   self.retry_interval)

        # Get a list of networks supported by the node.
        netkey_bytes_written = self.lib.btmesh.node.get_networks()

        if len(netkey_bytes_written) < 2:
            self.log.warning("LPN get networks provided invalid number of netkey bytes " +
                             f"({len(netkey_bytes_written)})")

        else:
            # List of network IDs. Each ID is two bytes in little-endian format.
            networks = netkey_bytes_written.networks
            self.lpn_friend_netkey_idx = (networks[1] << 8) | networks[0]
            self.lpn_establish_friendship(self.lpn_friend_netkey_idx)

    def lpn_feature_deinit(self):
        """ Deinitialize LPN functionality. """
        if self.lpn_active == 0:
            # LPN feature is currently inactive
            return
        # Cancel active friendship timer
        if self.LPNTimeout.friendship_timer is not None:
            if self.LPNTimeout.friendship_timer.is_alive():
                self.LPNTimeout.friendship_timer.cancel()
        # Terminate friendship if exist
        try:
            self.lib.btmesh.lpn.terminate_friendship(self.lpn_friend_netkey_idx)
        except bgapi.bglib.CommandFailedError as err:
            self.log.info(f"Friendship termination failed {err}")

        self.lib.btmesh.lpn.deinit()
        self.lpn_active = 0
        self.log.info("LPN deinitialized")

    def lpn_establish_friendship(self, netkey_idx):
        """ Establish friendship. """
        self.lib.btmesh.lpn.establish_friendship(netkey_idx)
        self.log.info("Establish friendship message sent")

    def set_configuraton_timer(self, delay_ms):
        """ 
        Set the timer that delays LPN initialization to enable 
        quick configuration over advertising bearer. 
        """
        if self.lpn_active != 1:
            threading.Timer(delay_ms * 0.001, self.lpn_feature_init).start()
```
