## Supplementary material for "VesperFlow: An automated pipeline for decoding multisensor data from biologging tags": onoff_client.py

```
"""
BtMesh Switch NCP-host Generic OnOff Client Model.
"""

# Copyright 2022 Silicon Laboratories Inc. www.silabs.com
#
# SPDX-License-Identifier: Zlib
#
# The licensor of this software is Silicon Laboratories Inc.
#
# This software is provided 'as-is', without any express or implied
# warranty. In no event will the authors be held liable for any damages
# arising from the use of this software.
#
# Permission is granted to anyone to use this software for any purpose,
# including commercial applications, and to alter it and redistribute it
# freely, subject to the following restrictions:
#
# 1. The origin of this software must not be misrepresented; you must not
#    claim that you wrote the original software. If you use this software
#    in a product, an acknowledgment in the product documentation would be
#    appreciated but is not required.
# 2. Altered source versions must be plainly marked as such, and must not be
#    misrepresented as being the original software.
# 3. This notice may not be removed or altered from any source distribution.

import threading
import os.path
import sys
sys.path.append(os.path.join(os.path.dirname(__file__), "../.."))

from common.util import BtMeshApp
import common.btmesh_models as model

# No flags used for message
NO_FLAGS = 0
class OnOffClient(BtMeshApp):
    """ Implement Generic OnOff Client Model specific APIs. """
    def __init__(self, connector, **kwargs):
        super().__init__(connector=connector, **kwargs)
        # On/off transaction identifier
        self.onoff_trid = 0
        # Delay time (in milliseconds) before starting the state change
        self.request_delay = 50
        # Transition time (in milliseconds) for the state change
        # Using zero transition time by default
        self.transtime = 0
        # How many times OnOff model messages are to be sent out for reliability
        self.onoff_request_count = 3
        # Latest desired light state
        # switch_pos = 1 -> turn lights on
        # switch_pos = 0 -> turn lights off
        self.switch_pos = 0

    def send_onoff_request(self):
        """ 
        Call 'onoff_request' with 'request_delay' intervals for 
        'ctl_request_count' times. 
        """
        # Increment transaction ID for each request, unless it's a retransmission
        self.onoff_trid += 1
        self.onoff_trid %= 256

        # Starting two new timer threads for the second and third message
        for count in range(1, self.onoff_request_count, 1):
            threading.Timer(count * self.request_delay * 0.001,
                            self.onoff_request,
                            args=[self.switch_pos, self.onoff_request_count - count]).start()

        # First message with 0ms delay
        self.onoff_request(self.switch_pos, self.onoff_request_count)

    def onoff_request(self, switch_pos, count):
        """
        Serialize and publish Generic On/Off requests and calculate the delay for them.

        :param switch_pos: holds the latest desired light state
        :param count: number of repetition
        """
        delay = (count - 1) * self.request_delay
        if count > 0:
            self.lib.btmesh.generic_client.publish(
                0,
                model.BTMESH_GENERIC_ON_OFF_CLIENT_MODEL_ID,
                self.onoff_trid,
                self.transtime,
                delay,
                NO_FLAGS,
                self.lib.btmesh.generic_client.SET_REQUEST_TYPE_REQUEST_ON_OFF,
                switch_pos.to_bytes(1, byteorder='little')
            )

            self.log.info(f"On/off request, trid: {self.onoff_trid}, delay: {delay}")

    def set_switch(self, set_switch):
        """
        Check if the given value is valid. Valid values are 0 or 1.
        If the value is not valid then the function adjusts it to be a proper value.

        :param set_switch: desired light state given by the user
        """
        if set_switch > 1:
            self.switch_pos = 1
        elif set_switch < 0:
            self.switch_pos = 0
        else:
            self.switch_pos = set_switch

        self.send_onoff_request()
```
