## Supplementary material for "VesperFlow: An automated pipeline for decoding multisensor data from biologging tags": README.md

# Bluetooth Mesh - NCP Host Switch (Lighting Client)

This example demonstrates the operation of a Bluetooth Mesh Lighting Client (Switch). 
The graphical user interface (GUI) simulates the behavior of a switch that controls
another device implementing the server role (for example Bluetooth Mesh - SoC Light). The 
switch can control the state, lightness, color temperature, Delta UV and
scenes on a remote device. It also demonstrates LPN feature to optimize battery life.

The following BT Mesh Models are demonstrated:
 * Generic OnOff Client
 * Light Lightness Client
 * Light CTL Client
 * Scene Client

## Getting Started

To learn Bluetooth Mesh technology fundamentals, see [Bluetooth Mesh Network - An Introduction for Developers](https://www.bluetooth.com/wp-content/uploads/2019/03/Mesh-Technology-Overview.pdf).

To get to know the Bluetooth Mesh SDK and Simplicity Studio, see [ QSG176: Bluetooth® Mesh SDK v2.x Quick Start Guide](https://www.silabs.com/documents/public/quick-start-guides/qsg176-bluetooth-mesh-sdk-v2x-quick-start-guide.pdf).

### Follow this procedure to get started

1. Install pyBGAPI.
	```
    pip install pybgapi
    ```
2. Clone the [repo](https://github.com/SiliconLabs/pybgapi-examples) from GitHub and check out the master barnch.
3. Flash the **Bluetooth Mesh - NCP Empty** demo binary on your [Wireless Development Kit](https://www.silabs.com/development-tools/wireless)
    using [Simplicity Studio](https://docs.silabs.com/simplicity-studio-5-users-guide/latest/ss-5-users-guide-building-and-flashing/flashing).

4. Run the btmesh-switch example from the repo.
	```
	python3 example/btmesh_switch/app.py
	```
5. Provision the switch example into the mesh network with a Provisioner device.<br>
	Recommended option: Silicon Labs Bluetooth Mesh smartphone application ([ QSG176: Bluetooth® Mesh SDK v2.x Quick Start Guide](https://www.silabs.com/documents/public/quick-start-guides/qsg176-bluetooth-mesh-sdk-v2x-quick-start-guide.pdf)).
	
	Other options:
	 * Bluetooth NCP Commander within Simplicity Studio
	 * Host Provisioner example in Bluetooth Mesh SDK
6. Make sure that a device implementing the Server role is in the mesh network.
	Recommended option: pyBGAPI light example ([Bluetooth mesh - Light](example/btmesh_light))

	Other options:
	* Bluetooth Mesh - SoC Light in Bluetooth Mesh SDK
7. How to use:<br>
	Note: if you configure the devices with the mobile app, in order to have the desired performance you need to have the same client-server functionalities configured on the devices. For example if you want to recall a saved scene then the functionality for the devices should be Scene Client and Scene Server.

	* Use the switch app's GUI to switch on the lightbulb

	![OnOff](images/OnOff.png)
	
	* Use the switch app's Lightness slider to change the lightbulb lightness

	![Lightness](images/Lightness.png)

	* Use the switch app's Temperature slider to change the color temperature

	![CTL](images/CTL.png)

	* To recall, store or delete a scene enter the scene number and push the desired button to execute

	![Scene](images/Scene.png)

## Structure of the Python Application

![Class](images/class.png)

- OnOffClient class: this class implements the Generic OnOff Client model.

- LightnessClient class: this class implements the Light Lightness Server model.

- CTLClient class: this class implements the Light CTL Client model.

- SceneClient class: this class implements the Scene Client model.

- Reset class: this class extends the BtMeshApp class and implements the node reset and full factory reset functions.

- LPN class: this class implements LPN functionalities and tries to establish friendship.

- App class: this class handles the received events and forwards them to the right class for processing.
After running the application two threads are created. One of the threads is responsible for the BT Mesh event handling while the other thread is responsible for the GUI.

- windows and MainPage class: this class extends the Tk class (tkinter library).
The windows class is a general class for making tkinter windows, here you can add the name of the window, the size of the window etc.
The MainPage class defines the actual layout of the window, variables and functions.

The example extends the BtMeshApp class. The BtMeshApp class implements the following: Check BGAPI Version, Initialize Mesh Node, Factory and Node Reset. It uses both Bluetooth and Bluetooth Mesh XAPI files.
