## Supplementary material for "VesperFlow: An automated pipeline for decoding multisensor data from biologging tags": scene_client.py

```
"""
BtMesh Switch NCP-host Scene Client Model.
"""

# Copyright 2022 Silicon Laboratories Inc. www.silabs.com
#
# SPDX-License-Identifier: Zlib
#
# The licensor of this software is Silicon Laboratories Inc.
#
# This software is provided 'as-is', without any express or implied
# warranty. In no event will the authors be held liable for any damages
# arising from the use of this software.
#
# Permission is granted to anyone to use this software for any purpose,
# including commercial applications, and to alter it and redistribute it
# freely, subject to the following restrictions:
#
# 1. The origin of this software must not be misrepresented; you must not
#    claim that you wrote the original software. If you use this software
#    in a product, an acknowledgment in the product documentation would be
#    appreciated but is not required.
# 2. Altered source versions must be plainly marked as such, and must not be
#    misrepresented as being the original software.
# 3. This notice may not be removed or altered from any source distribution.

import os
import sys
import threading
import bgapi.bglib
sys.path.append(os.path.join(os.path.dirname(__file__), "../.."))
from common.util import BtMeshApp

IMMEDIATE = 0
IGNORED = 0
NO_FLAGS = 0
class SceneClient(BtMeshApp):
    """ Implement Scene Client Model specific APIs. """

    def __init__(self, connector, **kwargs):
        super().__init__(connector=connector, **kwargs)
        # Scene recall transaction identifier
        self.scene_recall_trid = 0
        # How many times scene model messages are to be sent out for reliability
        self.scene_request_count = 3
        # Delay time (in milliseconds) before starting the state change
        self.request_delay = 50
        # Currently selected scene
        self.scene_number = 0
        # Address zero is used in scene client commands to indicate that message should be published
        self.publish_address = 0x0000
        # Parameter ignored for publishing
        self.app_key = 0
        # Scene number to store in a state of the light
        self.scene_store_number = 0

    def send_scene_recall_request(self):
        """ Publish scene recall request to recall a saved state. """
        # Increment transaction ID for each request, unless it's a retransmission.
        self.scene_recall_trid += 1
        self.scene_recall_trid %= 256

        # Starting two new timer threads for the second and third message.
        for count in range(1, self.scene_request_count, 1):
            threading.Timer(count * self.request_delay * 0.001,
                            self.scene_recall,
                            args=[self.scene_request_count - count]).start()

        # First message with 0ms waiting
        self.scene_recall(self.scene_request_count)

    def check_scene_recall(self, scene_to_recall):
        """
        Check if the given value is valid. Ignore invalid values.

        :param scene_to_recall: desired state to be recalled given by the user
        """
        if scene_to_recall <= 0 or scene_to_recall > 255:
            self.log.error("Wrong scene number! Possible values are between 1 and 255.")
        else:
            self.scene_number = scene_to_recall
            self.send_scene_recall_request()

    def check_scene_delete(self, scene_to_delete):
        """
        Check if the given value is valid. Ignore invalid values.

        :param scene_to_recall: desired state to be recalled given by the user
        """
        if scene_to_delete <= 0 or scene_to_delete > 255:
            self.log.error("Wrong scene number! Possible values are between 1 and 255.")
        else:
            self.scene_delete(scene_to_delete)

    def check_scene_store(self, scene_to_store):
        """
        Check if the given value is valid. Ignore invalid values.

        :param scene_to_store: actual state to be stored in the user given number
        """
        if scene_to_store <= 0 or scene_to_store > 255:
            self.log.error("Wrong scene number! Possible values are between 1 and 255.")
        else:
            self.scene_store(scene_to_store)

    def scene_recall(self, count):
        """
        Serialize and publish Scene recall requests, and calculate the delay for them.

        :param count: number of the scene to be recalled
        """
        delay = (count - 1) * self.request_delay
        if count > 0:
            try:
                self.lib.btmesh.scene_client.recall(
                    self.publish_address,
                    0,
                    self.scene_number,
                    IGNORED,
                    NO_FLAGS,
                    self.scene_recall_trid,
                    IMMEDIATE,
                    delay,
                )

                self.log.info(
                    f"Scene recall request, trid: {self.scene_recall_trid}, delay: {delay}"
                )
            except bgapi.bglib.CommandFailedError as err:
                self.log.error("%s", err.errorcode)

    def scene_store(self, scene_to_store):
        """Serialize scene store request."""
        try:
            self.lib.btmesh.scene_client.store(
                self.publish_address,
                0,
                scene_to_store,
                self.app_key,
                NO_FLAGS,
            )
            self.log.info(f"Scene store request, scene store number: {scene_to_store}")
        except bgapi.bglib.CommandFailedError as err:
            self.log.error("%s", err.errorcode)

    def scene_delete(self, scene_to_delete):
        """Serialize scene delete request."""
        try:
            self.lib.btmesh.scene_client.delete(
                self.publish_address,
                0,
                scene_to_delete,
                self.app_key,
                NO_FLAGS,
            )

            self.log.info(f"Scene store request, scene delete number: {scene_to_delete}")
        except bgapi.bglib.CommandFailedError as err:
            self.log.error("%s", err.errorcode)
```
