## Supplementary material for "VesperFlow: An automated pipeline for decoding multisensor data from biologging tags": switch_gui.py

```
"""
BtMesh Switch NCP-host GUI.
"""

# Copyright 2022 Silicon Laboratories Inc. www.silabs.com
#
# SPDX-License-Identifier: Zlib
#
# The licensor of this software is Silicon Laboratories Inc.
#
# This software is provided 'as-is', without any express or implied
# warranty. In no event will the authors be held liable for any damages
# arising from the use of this software.
#
# Permission is granted to anyone to use this software for any purpose,
# including commercial applications, and to alter it and redistribute it
# freely, subject to the following restrictions:
#
# 1. The origin of this software must not be misrepresented; you must not
#    claim that you wrote the original software. If you use this software
#    in a product, an acknowledgment in the product documentation would be
#    appreciated but is not required.
# 2. Altered source versions must be plainly marked as such, and must not be
#    misrepresented as being the original software.
# 3. This notice may not be removed or altered from any source distribution.

from dataclasses import dataclass
import tkinter
import logging

class ResizableWindow(tkinter.Tk):
    """ Root window for the frame. """
    def __init__(self, app, *args, **kwargs):
        tkinter.Tk.__init__(self, *args, **kwargs)
        self.protocol("WM_DELETE_WINDOW", lambda args=app: self.exit(args))
        self.resizable(1, 1)

        # Adding a title to the window
        self.wm_title("BT Mesh Switch")

        # Specifying the region where the frame is grided
        self.geometry("410x210")
        self.rowconfigure(0, weight=1)
        self.columnconfigure(0, weight=1)

        frame = MainPage(self, app)
        frame.grid(row=0, column=0, sticky=tkinter.NSEW)

    def exit(self, app):
        """ Shuting down the app. """
        app.stop()
        self.destroy()
        
class MainPage(tkinter.Frame):
    """ Implementation of the used frame. """
    __instance = None
    def get_instance(self):
        """ Getting GUI instance. """
        if MainPage.__instance is None:
            self.gui_log.info("No instance for GUI yet")
        return MainPage.__instance

    def __init__(self, parent, app):
        self.gui_log = logging.getLogger(str(type(self)))
        tkinter.Frame.__init__(self, parent)
        if MainPage.__instance is not None:
            self.gui_log.error("GUI instance is already present")
            return
        self.root = self
        self._job = None
        self.app = app
        MainPage.__instance = self

        # Variables for the scale inputs
        self.CTLLightbulbState.lightness_value = tkinter.IntVar()
        self.CTLLightbulbState.temperature_value = tkinter.IntVar()
        self.CTLLightbulbState.temperature_value.set(6500)
        self.CTLLightbulbState.delta_uv_value = tkinter.DoubleVar()
        self.CTLLightbulbState.last_lightness_value = tkinter.IntVar()

        # Lightness
        slider_label = tkinter.Label(
            self,
            text='Lightness:'
        )

        slider_label.grid(
            column=0,
            row=0,
            sticky=tkinter.EW,
            columnspan=2
        )

        slider = tkinter.Scale(
            self,
            from_=0,
            to=100,
            orient='horizontal',
            command=self.lightness_slider_changed,
            variable=self.CTLLightbulbState.lightness_value
        )

        slider.grid(
            column=2,
            row=0,
            sticky=tkinter.EW,
            columnspan=4
        )

        self.lightness_value_label = tkinter.Label(
            self,
            text=f"{self.CTLLightbulbState.lightness_value.get()}%"
        )

        self.lightness_value_label.grid(
            row=0,
            column=6,
            sticky=tkinter.EW,
            padx=5,
            pady=5
        )

        # Temperature
        slider_label = tkinter.Label(
            self,
            text='Temperature:'
        )

        slider_label.grid(
            column=0,
            row=1,
            sticky=tkinter.EW,
            columnspan=2
        )

        slider = tkinter.Scale(
            self,
            from_=800,
            to=20000,
            orient='horizontal',
            command=self.temperature_slider_changed,
            variable=self.CTLLightbulbState.temperature_value,
        )

        slider.grid(
            column=2,
            row=1,
            sticky=tkinter.EW,
            columnspan=4
        )

        self.temperature_value_label = tkinter.Label(
            self,
            text=f"{self.CTLLightbulbState.temperature_value.get()}K"
        )

        self.temperature_value_label.grid(
            row=1,
            column=6,
            sticky=tkinter.EW,
            padx=5,
            pady=5
        )

        # Delta UV
        slider_label = tkinter.Label(
            self,
            text='Delta UV:'
        )

        slider_label.grid(
            column=0,
            row=2,
            sticky=tkinter.EW,
            columnspan=2
        )

        slider = tkinter.Scale(
            self,
            from_=-1.00,
            to=1.00,
            orient='horizontal',
            command=self.delta_uv_slider_changed,
            variable=self.CTLLightbulbState.delta_uv_value,
            resolution=0.01,
            digits=3
        )

        slider.grid(
            column=2,
            row=2,
            sticky=tkinter.EW,
            columnspan=4
        )

        self.delta_uv_value_label = tkinter.Label(
            self,
            text=f"{self.CTLLightbulbState.delta_uv_value.get():.2f}"
        )
        self.delta_uv_value_label.grid(
            row=2,
            column=6,
            sticky=tkinter.EW,
            padx=5,
            pady=5
        )

        # On/Off button
        self.on_off_button = tkinter.Button(
            self,
            text='On',
            command=self.on_off_button_pushed
        )

        self.on_off_button.grid(
            row=0,
            column=7,
            sticky=tkinter.EW,
            padx=5,
            pady=5,
        )

        # Factory reset button
        factory_reset = tkinter.Button(
            self,
            text="Factory reset",
            command=self.factory_reset
        )

        factory_reset.grid(
            row=3,
            column=4,
            sticky=tkinter.EW,
            columnspan=2
        )

        # Node reset button
        node_reset = tkinter.Button(
            self,
            text="Node reset",
            command=self.node_reset
        )

        node_reset.grid(
            row=3,
            column=2,
            sticky=tkinter.EW,
            padx=(0, 5),
            columnspan=2
        )

        # Scene
        scene_label = tkinter.Label(
            self,
            text="Scene: "
        )

        scene_label.grid(
            row=4,
            column=0,
            sticky=tkinter.EW,
            padx=5,
            pady=5,
            columnspan=2
        )

        self.scene_entry = tkinter.Entry(
            self,
            width=10,
            borderwidth=5,
        )

        self.scene_entry.grid(
            row=4,
            column=2,
            sticky=tkinter.W,
            pady=5,
            columnspan=2
        )

        scene_recall = tkinter.Button(
            self,
            text="Recall",
            command=self.recall_scene
        )

        scene_recall.grid(
            row=4,
            column=5,
            sticky=tkinter.EW,
            pady=5
        )

        scene_delete = tkinter.Button(
            self,
            text="Delete",
            command=self.delete_scene
        )

        scene_delete.grid(
            row=4,
            column=6,
            sticky=tkinter.EW,
            padx=5,
            pady=5
        )

        store_scene = tkinter.Button(
            self,
            text="Store",
            command=self.store_scene
        )

        store_scene.grid(
            row=4,
            column=4,
            sticky=tkinter.EW,
            padx=(0, 5),
            pady=5
        )

        self.columnconfigure((0,1,2,3,4,5,6,7), weight=1, uniform='column')
        self.rowconfigure((0,1,2,3,4), weight=1, uniform='row')

    @dataclass
    class CTLLightbulbState:
        """ Dataclass of CTL lightbulb state. """
        # Value of lightness
        lightness_value = None
        # Value of temperature
        temperature_value = None
        # Value of deltauv
        delta_uv_value = None
        # Value of last lightness
        last_lightness_value = None

    def lightness_slider_changed(self, _event):
        """ Manage lightness slider changes. """
        self.lightness_value_label.configure(text=
                                             f"{self.CTLLightbulbState.lightness_value.get()}%")
        if self._job:
            self.root.after_cancel(self._job)
        self._job = self.root.after(120, self.app.set_lightness(
            self.CTLLightbulbState.lightness_value.get()))
        self.CTLLightbulbState.last_lightness_value.set(
            self.CTLLightbulbState.lightness_value.get())
        if self.CTLLightbulbState.lightness_value.get() != 0:
            self.on_off_button.config(text="Off")
        else:
            self.on_off_button.config(text="On")

    def temperature_slider_changed(self, _event):
        """ Manage the temperature slider changes. """
        self.temperature_value_label.configure(
            text=f"{self.CTLLightbulbState.temperature_value.get()}K")
        if self._job:
            self.root.after_cancel(self._job)
        self._job = self.root.after(120,
                                    self.app.set_temperature(
                                        self.CTLLightbulbState.last_lightness_value.get(),
                                        self.CTLLightbulbState.temperature_value.get()))

    def delta_uv_slider_changed(self, _event):
        """ Manage the delta UV slider changes. """
        self.delta_uv_value_label.configure(text=self.CTLLightbulbState.delta_uv_value.get())
        if self._job:
            self.root.after_cancel(self._job)
        self._job = self.root.after(120,
                                    self.app.set_delta_uv(
                                        self.CTLLightbulbState.lightness_value.get(),
                                        self.CTLLightbulbState.delta_uv_value.get()))

    def on_off_button_pushed(self):
        """ Send out On/Off messages and updete the gui. """
        if self.CTLLightbulbState.lightness_value.get() == 0:
            if self.CTLLightbulbState.last_lightness_value.get() != 0:
                self.CTLLightbulbState.lightness_value.set(
                    self.CTLLightbulbState.last_lightness_value.get())
                self.lightness_value_label.configure(
                    text=f"{self.CTLLightbulbState.lightness_value.get()}%")
            else:
                self.CTLLightbulbState.lightness_value.set(100)
                self.lightness_value_label.configure(
                    text=f"{self.CTLLightbulbState.lightness_value.get()}%")
            self.on_off_button.config(text='Off')
            self.app.set_switch(1)

        else:
            self.CTLLightbulbState.lightness_value.set(0)
            self.lightness_value_label.configure(text=
                                                 f"{self.CTLLightbulbState.lightness_value.get()}%")
            self.on_off_button.config(text='On')
            self.app.set_switch(0)

    def factory_reset(self):
        """ Call the factory reset function from the General class. """
        self.app.factory_reset()

    def node_reset(self):
        """ Call the node reset function from the General class. """
        self.app.node_reset()

    def recall_scene(self):
        """ Call the scene store function from the SceneClient class. """
        to_recall = self.scene_entry_check()
        if to_recall is not None:
            self.app.check_scene_recall(to_recall)

    def delete_scene(self):
        """ Call the scene delete function from the SceneClient class. """
        to_delete = self.scene_entry_check()
        if to_delete is not None:
            self.app.check_scene_delete(to_delete)            

    def store_scene(self):
        """ Call the scene store function from the SceneClient class. """
        to_store = self.scene_entry_check()
        if to_store is not None:
            self.app.check_scene_store(to_store)

    def scene_entry_check(self):
        """ Check if empty parameter was given after a scene button press. """
        if self.scene_entry.get() == "":
            return None
        return int(self.scene_entry.get())


def switch_gui_start(app):
    """ Starting the GUI. """
    gui = ResizableWindow(app=app)
    gui.mainloop()
```
