## Supplementary material for "VesperFlow: An automated pipeline for decoding multisensor data from biologging tags": README.md

# Proxtit Proximity system downloader example based on PyBGAPI Examples

This repo contains example source code based on [pyBGAPI](https://pypi.org/project/pybgapi/) that
implement simple Bluetooth app to communicate with Proxtit proximity loggers via Bluetooth for demonstration purposes.
These examples can be used as references to implement custom Bluetooth in Python in just a few minutes without writing a single line of embedded code.

## TL;DR

1. Install pyBGAPI.
    ```
    pip install pybgapi
    ```

2. Plug provided USB Bluetooth communicator based on BG22 SoC.

3. Run an arbitrary example from the repo or the downloader/app.py.
    ```
    python3 example/bt_empty/app.py
    ```
    or
    ```
    python3 downloader/app.py
    ```

## Getting Started

To get started with Silicon Labs Bluetooth software, see
[QSG169: Bluetooth® SDK v3.x Quick Start Guide](https://www.silabs.com/documents/public/quick-start-guides/qsg169-bluetooth-sdk-v3x-quick-start-guide.pdf).
For Bluetooth Mesh, see
[QSG176: Bluetooth® Mesh SDK v2.x Quick-Start Guide](https://www.silabs.com/documents/public/quick-start-guides/qsg176-bluetooth-mesh-sdk-v2x-quick-start-guide.pdf).

In the NCP context, the application runs on a host MCU or a PC, which is the NCP Host, while the
Bluetooth stack runs on an EFR32, which is the NCP Target.

The NCP Host and Target communicate via a serial interface (UART). The communication between the NCP
Host and Target is defined in the Silicon Labs proprietary protocol, BGAPI. pyBGAPI is the reference
implementation of the BGAPI protocol in Python for the NCP Host.

On Linux systems, an additional CPC layer is available that provides additional security and
robustness. For further details, see [Co-Processor Communication](https://docs.silabs.com/gecko-platform/latest/service/cpc/overview).

[AN1259: Using the v3.x Silicon Labs Bluetooth® Stack in Network CoProcessor Mode](https://www.silabs.com/documents/public/application-notes/an1259-bt-ncp-mode-sdk-v3x.pdf)
provides a detailed description how NCP works and how to configure it for CPC and custom hardware.

For the latest BGAPI documentation, see [docs.silabs.com](https://docs.silabs.com/bluetooth/latest/).
