## Supplementary figures and images for "VesperFlow: An automated pipeline for decoding multisensor data from biologging tags"

### class.png

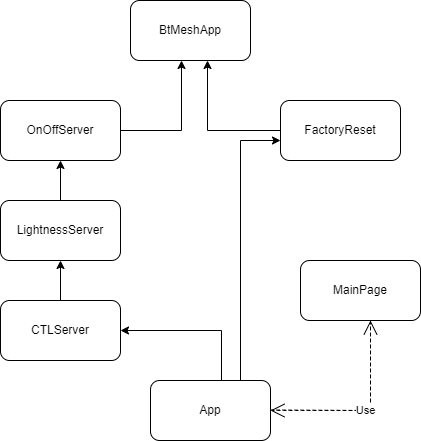

### class.png

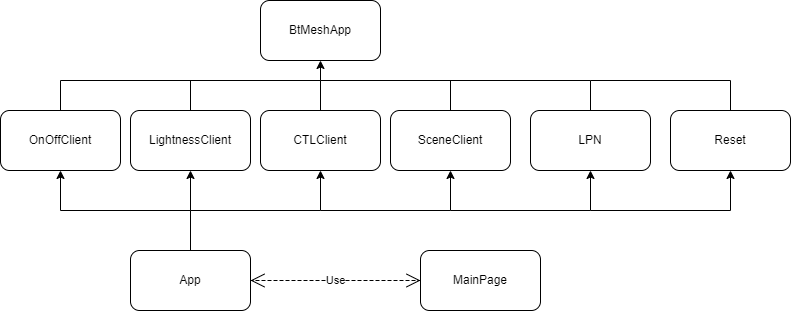

### CTL.png

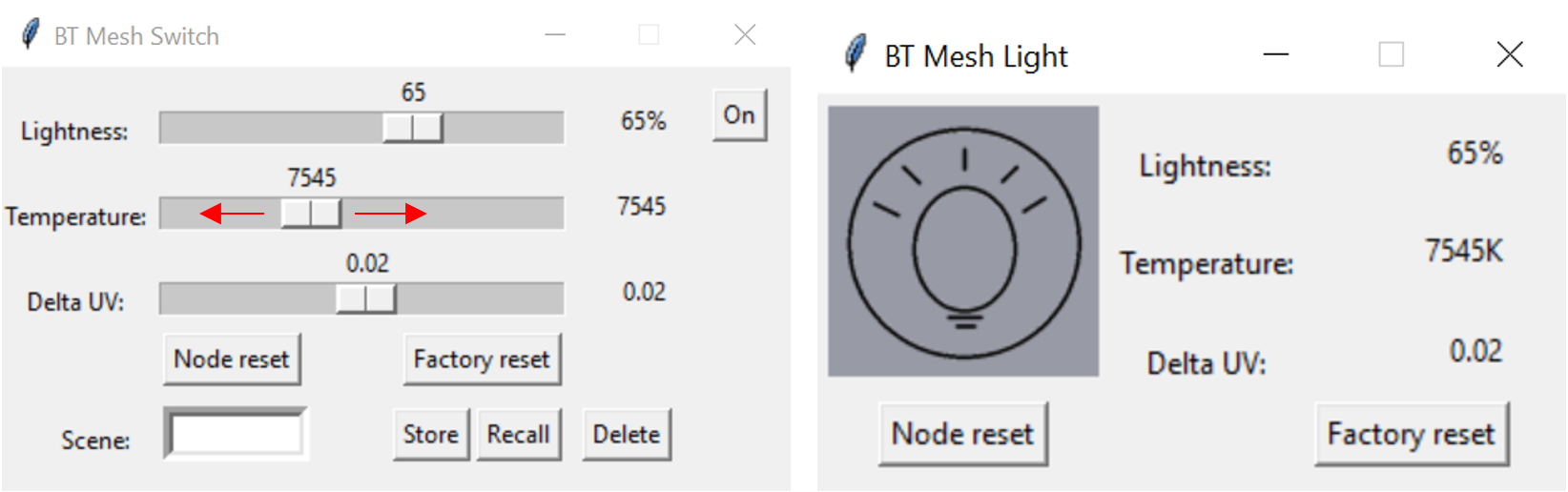

### graphviz_installer.png

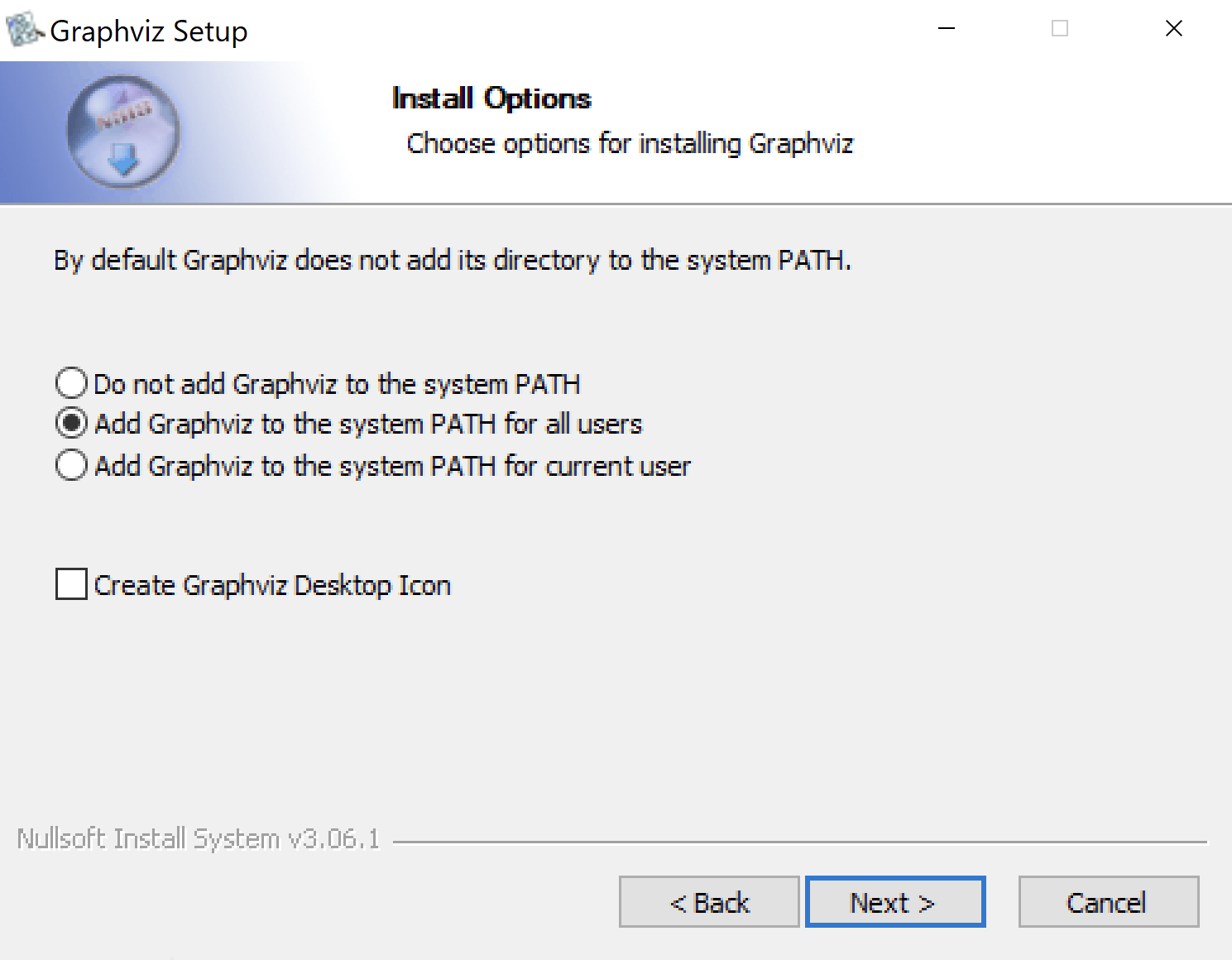

### lightbulb.png

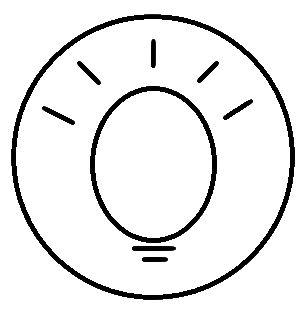

### Lightness.png

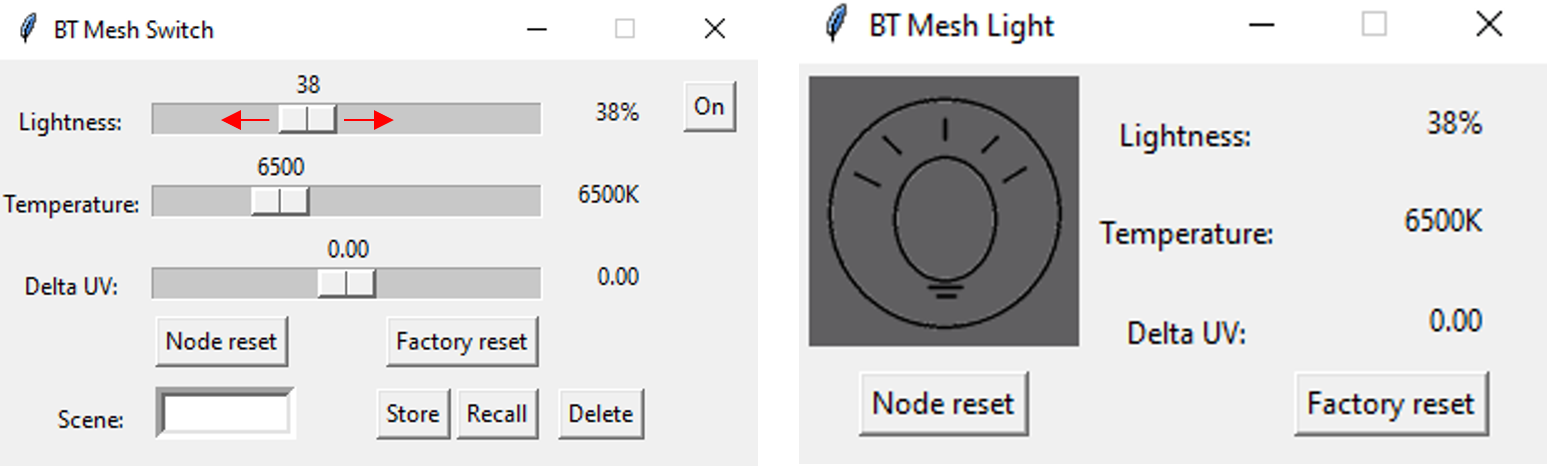

### OnOff.png

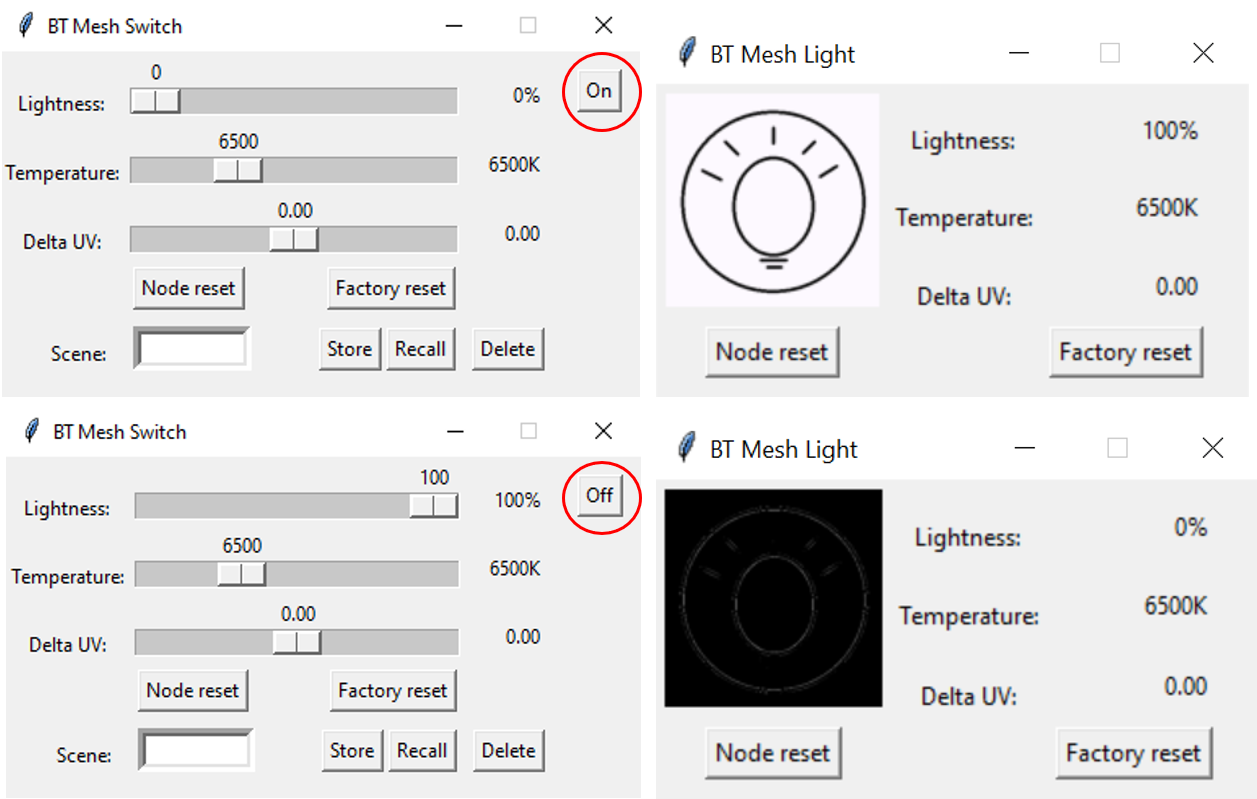

### roaming_network.png

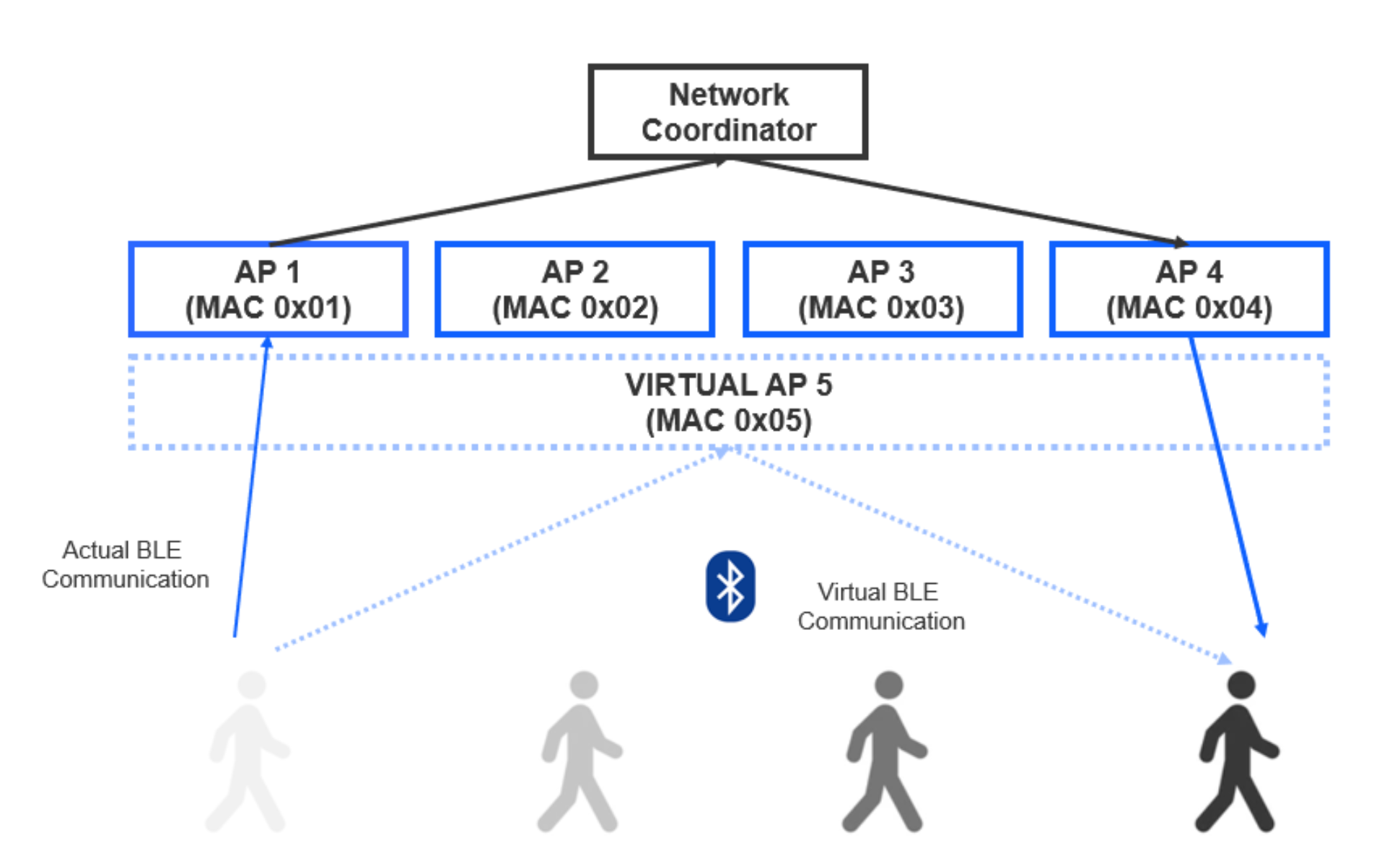

### Scene.png

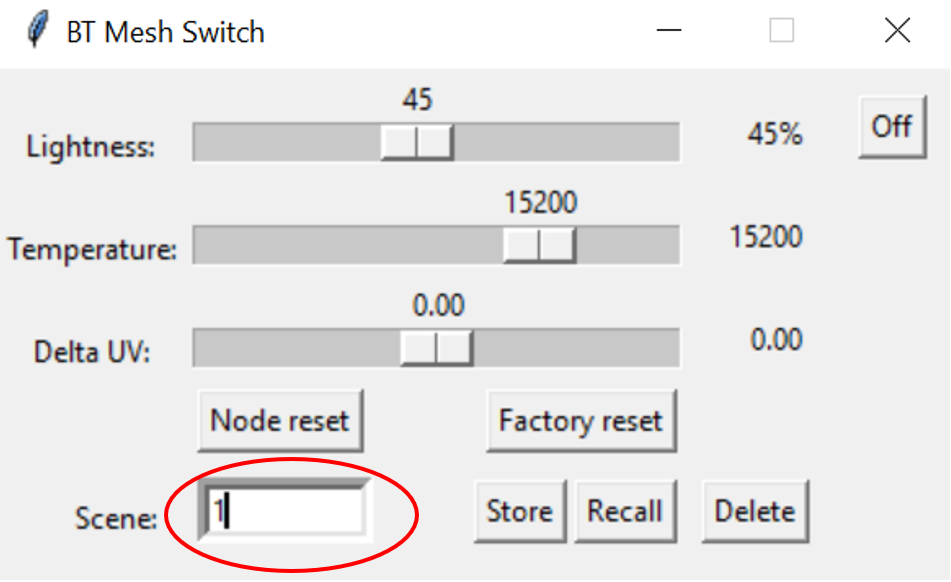
